# Demographic history shapes genomic variation in an intracellular parasite with a wide geographic distribution

**DOI:** 10.1101/2021.11.02.466881

**Authors:** Pascal Angst, Dieter Ebert, Peter D. Fields

## Abstract

Analyzing variation in a species’ genomic diversity can provide insights into its historical demography, biogeography and population structure, and thus, its ecology and evolution. Although such studies are rarely undertaken for parasites, they can be highly revealing because of the parasite’s coevolutionary relationships with hosts. Modes of reproduction and transmission are thought to be strong determinants of genomic diversity for parasites and vary widely among microsporidia (fungal-related intracellular parasites), which are known to have high intraspecific genetic diversity and interspecific variation in genome architecture. Here we explore genomic variation in the microsporidium *Hamiltosporidium*, a parasite of the freshwater crustacean *Daphnia magna*, looking especially at which factors contribute to nucleotide variation. Genomic samples from 18 Eurasian populations and a new, long-read based reference genome were used to determine the roles that reproduction mode, transmission mode and geography play in determining population structure and demographic history. We demonstrate two main *H. tvaerminnensis* lineages and a pattern of isolation-by-distance, but note an absence of congruence between these two parasite lineages and the two Eurasian host lineages. We suggest a comparatively recent parasite spread through Northern Eurasian host populations after a change from vertical to mixed-mode transmission and the loss of sexual reproduction. While gaining knowledge about the ecology and evolution of this focal parasite, we also identify common features that shape variation in genomic diversity for many parasites, e.g., distinct modes of reproduction and the intertwining of host–parasite demographies.

## Introduction

Uncovering the ecological and evolutionary forces that drive variation in genetic diversity within and among populations across a species’ range is important for understanding biological processes of basic and applied interest. Factors such as historical conditions (e.g., phylogeography and population structure), environmental conditions (e.g., adaptation to local climate), and stochastic processes (e.g., genetic drift and founder events) may contribute to patterns of genetic variation in a species (Bellis et al., 2020; Levicoy et al., 2021; Sendell-Price et al., 2021). For parasites, such studies take on an additional level of complexity, as parasite evolution is influenced by its dependence on the host and by issues such as host demography, parasite epidemiology, and coevolutionary interactions between hosts and parasites (Ebert & Fields, 2020). While a great deal of research has examined the effect of parasitism on host evolution, we know little about the factors that drive changes in the distribution of parasite genomic diversity, such as parasite life history, host shifts, mode of transmission and mode of reproduction (Janssen et al., 2004; Nieberding et al., 2008; Sandrock et al., 2011). The phylogeography of human parasites, which has been successfully studied to illuminate genetic epidemiology and variation in several systems, has proved to be a powerful tool in understanding, for example, *Mycobacterium tuberculosis* and viruses like the recently emerged coronavirus SARS-CoV-2 (Dellicour et al., 2021; Gagneux & Small, 2007). For the Gram-negative bacterium *Helicobacter pylori*, a known risk factor for stomach cancer in humans, the link between host and parasite biogeography and the parasite’s mode of transmission was shown to be an important factor in the evolution of the system (Montano et al., 2015): similarly, *H. pylori*, like its human host, shows a continuous loss of genetic diversity with increasing geographic distance from East Africa. Its vertical mode of transmission most likely contributed to the geographical distribution of genetic diversity shared between the parasite and its host. The interplay between co-phylogeography and mode of transmission here highlights the importance of studying interactions between factors that drive genetic diversity when trying to understand host and parasite evolution. In this article, we present a population genomic study of a parasitic microsporidium, focusing on the distribution of genomic diversity and relating it to key factors of parasite biology.

Genetic variation in parasites and their hosts may be influenced by the same or different factors; moreover they might influence each other’s evolution in both specific and general ways (Langerhans, 2008). In *H. pylori*, for example, the parasite’s genomic signature was so strong that it could be used to infer the history of the human host across the Pacific islands (Moodley et al., 2009). However, investigating interrelationships between multiple factors such as mode of transmission and biogeography in *H. pylori* is challenging, as both host and parasite need to be considered; therefore, such studies are still rare in parasite evolutionary biology.

One taxon that has recently gained attention as a potential model for parasite evolution is the microsporidia, due to its high variation in many aspects relevant to evolutionary and ecological theory (Murareanu et al., 2021; Wadi & Reinke, 2020). Microsporidia are obligate intracellular parasites with a phylogenetic position close to the fungi (James et al., 2013), although they show many highly derived features: for example, they have no mitochondria, have fewer genes and a very small genome size compared to other fungi, including the smallest known eukaryote genomes (Corradi, 2015). However, the latter two features are highly variable among microsporidia (Wadi & Reinke, 2020; Williams et al., 2008). Not only do microsporidia genomes vary tremendously in length and in gene numbers, but de-novo genome comparisons of microsporidia also found high levels of intraspecific genomic diversity (Pelin et al., 2015; Pombert et al., 2013). Moreover, microsporidia show high variation in their modes of reproduction and transmission, with ~18 % of species being thought to transmit vertically (Murareanu et al., 2021). Because disentangling and understanding the factors that drive genomic diversity may be easier in such a fast-evolving clade, microsporidia present a powerful model clade for evolutionary biology research.

The freshwater planktonic crustacean *Daphnia magna* is a well-established model system in ecology and evolution, including studies of host–parasite interactions (Altermatt & Ebert, 2008; Ebert, 2008; Orlansky & Ben-Ami, 2019). *D. magna* is parasitized by different species of microsporidia (Ebert, 2005), including the microsporidian genus *Hamiltosporidium* (Ebert, 2005; Haag et al., 2020; Pombert et al., 2015), of which two species, *H. magnivora* and *H. tvaerminnensis*, infect *D. magna* exclusively (Haag et al., 2011). Host and parasites are widespread across Eurasia. Both *Hamiltosporidium* species have, as compared to other microsporidia, relatively large, gene-sparse genomes, a feature that has been linked to a high number of repetitive, transposable elements (Parisot et al., 2014) that may have spread in the *Hamiltosporidium* genome due to the parasite’s ability to transmit vertically, which leads to bottlenecks and thus a decreased effective population size, *N_e_*, and reduced effectiveness of selection (de Albuquerque et al., 2020; Haag et al., 2020). Previous studies have found that *H. tvaerminnensis* reproduces asexually and has a mixed-mode transmission (i.e., horizontal and vertical), whereas its sister species, *H. magnivora*, reproduces sexually and transmits among *D. magna* hosts only vertically (Haag et al., 2011; Haag et al., 2013b). *H. magnivora* is believed to have a second, as yet unknown host, to which it is transmitted horizontally (Mangin et al., 1995). While diversity in aspects like modes of reproduction and transmission makes the *Hamiltosporidium* genus a good model for studying parasite evolution in general, it might represent a more unique system in terms of understanding the consequences of gaining or losing a second host and of changing the amount of recombination following a switch in reproductive mode.

Several studies on the distribution of genetic variation in microsporidia in diverse hosts have shown that host demography contributes to parasite phylogeny, population structure, and demographic history (e.g., Gómez-Moracho et al., 2015; Shafer et al., 2009; Wang et al., 2019). Existing knowledge about *D. magna* demographic history (Fields et al., 2018; Orsini et al., 2012, 2013; Stollewerk, 2010) allows us to go one step further and investigate the parasite’s genetic demography in relation to that of its host, looking at host–parasite co-phylogeny, co-biogeography, and coevolution. Previous studies on the *D. magna*–microsporidia host–parasite system have focused on mode of reproduction, demographic history, and transmission mode (Haag et al., 2013b; Haag et al., 2020) in an effort to understand the system’s colonization history and its transition from sexual to clonal reproduction. Here, we go a step further, reevaluating these questions using entire genomes sampled from a much larger swath of the species’ range, including Europe and Asia, and combining various approaches to investigate the evolutionary history and underlying mechanisms shaping genomic variation in this system. First, we ask how much intraspecific genomic variation exists in the microsporidian species *H. tvaerminnensis* using a new, highly improved long-read based reference genome. Based on a genomic study that estimated genomic variation in the microsporidium *Nosema ceranae* (Pelin et al., 2015), a close relative of *H. tvaerminnensis* with the same mode of transmission (Haag et al., 2020), a reasonable expectation was to find similar magnitudes of genetic diversity in *H. tvaerminnensis*. Second, using a multicontinental dataset, we ask how historical processes such as demography and phylogeography can explain the current distribution of genetic variation in *H. tvaerminnensis* across continental biogeographic scales? Knowing the phylogeography and demographic history of the host allowed us to test the co-phylogeography hypothesis. A finding that the parasite shares the same population structure and pattern of isolation-by-distance (IBD) as has been reported in the host (Andras et al., 2018; Fields et al., 2015) would support a model of phylogeographic co-cladogenesis. Since effects of demography and selection on genetic diversity are often hard to distinguish (Pavlidis et al., 2008), and nonadaptive processes may be an especially powerful force in the focal microsporidian genus (Haag et al., 2020), we also quantified the strength of positive selection in *H. tvaerminnensis*. Confirming weak selection strength would give us more confidence in our demography focused hypothesis and support previous findings (Haag et al., 2020).

## Material and Methods

### Daphnia magna Diversity Panel

The parasites examined in this study were derived from material collected within the framework of a large-scale biogeographic study of the host species, *D. magna* (Fields et al., 2015, 2018; Seefeldt & Ebert, 2019). Animals collected from across the species range were brought to the laboratory, and one iso-female line was created from each population (these iso-female lines are termed ‘clones’ for the course of this study). Animals from each clone were checked for microsporidia infections with phase-contrast microscopy, using squash-preparations or samples of the gut. The *D. magna* clones were sequenced with Illumina paired-end (PE) sequencing using a HiSeq 2500, with the respective parasite genome simultaneously sequenced in several cases. Whole-genome sequences of three *H. magnivora* and 15 *H. tvaerminnensis*, each from a different host clone collected from a different population, were obtained and used in the present study (Table 1). *H. tvaerminnensis* and *H. magnivora* are very closely related. Their species designation is based on morphological traits (Haag et al., 2011), but a genome-wide assessment of their relatedness has not been conducted thus far. Here, one common reference genome for the two *Hamiltosporidium* species was used.

**Table 1:** Sample information. Average whole-genome coverage for the parasites is denoted in the last column.

| Sample ID | Country | Latitude | Longitude | Parasite species | Coverage |
| --- | --- | --- | --- | --- | --- |
| BE-OM-2 | Belgium | 50.863280 | 4.721370 | *H. magnivora* | 594× |
| BE-T1-2 | Belgium | 50.823050 | 4.593940 | *H. magnivora* | 145× |
| BE-WH2-11 | Belgium | 51.334780 | 3.348130 | *H. magnivora* | 149× |
| CY-PA3-2 | Cyprus | 35.034060 | 33.954930 | *H. tvaerminnensis* | 598× |
| ES-DO1-1 | Spain | 36.978360 | -6.477640 | *H. tvaerminnensis* | 282× |
| ES-OM-5 | Spain | 37.254353 | -6.969114 | *H. tvaerminnensis* | 348× |
| FI-BR1-39-6 | Finland | 59.845253 | 23.274526 | *H. tvaerminnensis* | 219× |
| FI-FHS2-11-3 | Finland | 60.273780 | 27.218750 | *H. tvaerminnensis* | 94× |
| FI-OER-3-3 | Finland | 59.788590 | 23.174140 | *H. tvaerminnensis* | 1097× |
| IL-G-3 | Israel | 32.228840 | 34.831140 | *H. tvaerminnensis* | 703× |
| IL-PS-6 | Israel | 32.255710 | 34.853270 | *H. tvaerminnensis* | 689× |
| IL-SH-4 | Israel | 31.701030 | 34.713760 | *H. tvaerminnensis* | 600× |
| RU-BAYA1-1 | Russia | 53.017833 | 106.886833 | *H. tvaerminnensis* | 756× |
| RU-TY1-3 | Russia | 50.255500 | 89.546830 | *H. tvaerminnensis* | 389× |
| RU-TY2-2 | Russia | 50.212170 | 89.717830 | *H. tvaerminnensis* | 523× |
| RU-ZB1-1 | Russia | 50.413170 | 114.724703 | *H. tvaerminnensis* | 374× |
| SE-G4-20 | Sweden | 60.253040 | 18.306090 | *H. tvaerminnensis* | 562× |
| SE-H1-4 | Sweden | 58.342330 | 11.218030 | *H. tvaerminnensis* | 336× |

### Reference genome assembly

To obtain *D. magna* free of microbiota, except microsporidia, we followed a procedure described by Fields et al. (2015), using Qiagen Genomic Tips to obtain high molecular weight DNA from the FI-OER-3-3 clone, which was infected with *H. tvaerminnensis*. We used a Blue Pippin to isolate DNA fragments greater than 15 Kbp. A standard PacBio library was prepared, and one SMRTCELL was sequenced in CLR mode on a Sequel I at the D-BSSE (Basel, Switzerland).

After receiving the raw data in a subread BAM file, we initially mapped all reads to the *D. magna* reference genome (Fields et al., in prep). In order to achieve the most complete assembly of *H. tvaerminnensis*, we used Canu v.2.1.1 (Koren et al., 2017) with high sensitivity parameters to effectively work out the *H. tvaerminnensis* genome as part of the larger metagenome of *D. magna* and *H. tvaerminnensis* in the read set. We used BlobToolKit v.2 (Challis et al., 2020) to further isolate contigs specific to *H. tvaerminnensis*. By mapping Illumina data from an uninfected host genotype, we were able to further disentangle suspected *H. tvaerminnensis* contigs from contigs that likely belonged to the host. Afterwards, we used purge_haplotigs v.1.1.1 (Roach et al., 2018) to haplodize the assembly of contigs that presumably derived only from the parasite genome. Finally, we polished the resulting *H. tvaerminnensis* assembly using Genomic Concensus v.2.3.3 (Chin et al., 2013) and Pilon v.1.23 (Walker et al., 2014).

To annotate the new *H. tvaerminnensis* genome, we lifted the previously published annotation from Haag et al. (2020) over using RaGOO v.1.1 (Alonge et al., 2019) and evaluated the biological completeness of this new genome draft using BUSCO v.4.0.1 (Seppey et al., 2019) and its microsporidia_odb10 (Creation date: 2020-08-05).

### Mapping and variant calling

Raw reads were assessed for quality with FastQC v.0.11.7 (http://www.bioinformatics.babraham.ac.uk/projects/fastqc) and subsequently trimmed with Trimmomatic v.0.38 (Bolger et al., 2014) to remove low quality sequence and adapter contamination. Trimming success was assessed by doing a second run of FastQC. Quality trimmed reads were mapped to the reference genome with BWA MEM v.0.7.17 (H. Li, 2013). SAMtools v.1.9 (H. Li et al., 2009) was used to convert the SAM files to BAM files, coordinate sort individual BAM files, and remove unmapped reads. Read groups were added and duplicates marked for individual BAM files using Picard Toolkit v.2.18.16 (Broad Institute, 2019). The average read depths were computed using SAMtools function depth. GATK v.3.8 HaplotypeCaller was used to generate variant calls (McKenna et al., 2010; Van der Auwera et al., 2013), specifically GVCFs, which were first generated for individual BAM files. We used the GATK function GenotypeGVCF to combine GVCFs into all-site VCFs. Variant calls were generated for *H. tvaerminnensis–H. magnivora* in the form of a VCF file, which was filtered to exclude INDEL variants with VCFtools v.0.1.16 (Danecek et al., 2011).

### Coding sequences – single copy orthologs

Protein coding regions are of special interest for phylogenetic analyses and for identifying signals of selection (including its efficacy) across the genome. In order to characterize variation in these particular subsections of the genome, we needed to extract subsets of the genome-wide VCF. Protein sequences of *H. magnivora* (BEOM2 v1; GenBank accession: PITI00000000.1) were downloaded from NCBI and protein datasets of *H. tvaerminnensis* and *H. magnivora* were used as input for OrthoMCL v.2.0.9 to find one-to-one orthologs between the two (L. Li et al., 2003, following specifically the automation pipeline described at the Github repository: https://github.com/apetkau/orthomcl-pipeline). The orthologous sequences of *H. tvaerminnensis* and *H. magnivora* were aligned with PRANK v.170427 (Löytynoja, 2014) using a custom script adapted from Fields et al. (in review). After an initial survey of pairwise alignment quality, we implemented a masking step, in which excessively divergent or poorly aligned sequences were excluded from downstream analysis. We then used the R v.3.5.1 (R Core Team, 2018) package *seqinR* v.3.4-5 (Charif & Lobry, 2007) to import FASTA alignments and *PopGenome* v.2.7.1 (Pfeifer et al., 2014) to manipulate the VCF files. To produce multiple sequence alignments, pseudo-haplotypes (i.e., a random assignment of SNPs to one or the other haplotype) were generated by recoding the structure of the VCF file. One variant-imputed version of the reference per pseudo-haplotype was produced for each *H. tvaerminnensis* sample with the function vcf2fasta of Vcflib v.1.0.0-rc2 (Garrison, 2019), and then coding sequences of interest were cut out of the variant-imputed versions of the reference with the script gff2fasta.pl (https://github.com/ISUgenomics/common_scripts). The resultant coding sequences were aligned to the initial *H. tvaerminnensis* and *H. magnivora* alignments using MAFFT v.7.407 (Katoh et al., 2002; Katoh & Standley, 2013) and its --add option. Next, all positions masked in the two species reference alignments were masked in the multi sequence-alignments with generate_masked_ranges.py (https://gist.github.com/danielecook) and BEDtools v.2.27.1 (Quinlan, 2014) function maskfasta.

### Sequence variation and population genetic analyses

Per-site nucleotide differences (π) were calculated along the genome for *H. tvaerminnensis* in 1 Kbp windows using pixy v.0.95 (Korunes & Samuk, 2021), and the difference between lineages was statistically tested using a Wilcoxon signed rank test in R. Alleles of less than half and more than double the average sample coverage were filtered out for the calculation of π. Similarly, π was calculated for the coding sequences of *H. tvaerminnensis* using the script selectionStats.py (https://github.com/tatumdmortimer/popgen-stats). Both methods consider invariant sites in their calculations, making calculations more consistent with theoretical expectations and more comparable among species (Korunes & Samuk, 2021). This analysis was not conducted with *H. magnivora*, as only three genomes were available.

### Population structure and phylogenetic analyses

We conducted both a principle component analysis (PCA) and a cluster analysis with the *Hamiltosporidium* whole-genome polymorphism data using the R packages *SNPRelate* v.1.14.0 and *gdsfmt* v.1.16.0 (Zheng et al., 2012). Additionally, the pattern of IBD was tested for *H. tvaerminnensis* by comparing the pairwise genetic differentiation of the samples with their pairwise geographical distance. The R base *stats* function dist was used to calculate the pairwise Euclidean distance, while *geodist* v.0.0.3 (Padgham & Sumner, 2019) was used to calculate the geographical distance between the samples. *VCFR* and *adegenet* v.2.1.2 (Jombart, 2008) were used in R for file import and format conversions. The correlation between differentiation and geographical distance was tested with a dbMEM analysis by RDA. Specifically, we transformed the explanatory variable, geographic distance, into dbMEMs using the R package *adespatial* v.0.3-14 (Dray et al., 2021) and decomposed the response variable, genetic differentiation, into principal components using the R base *stats* function prcomp. The RDA was done in R using the package *vegan* v.2.5-7 (Oksanen et al., 2020) with significance assessed with 1,000 permutations. In addition to looking at whole-genome diversity, we also examined the phylogenetic signal in protein-coding regions of the genome directly. As the aforementioned pseudo-haplotypes are inappropriate for a number of phylogenetic methods that rely on haplotype information, we used ambiguity codes, an alternative method for unknown phase. We used the GATK-functions SelectVariants and FastaAlternateReferenceMaker to generate variant-imputed versions of the *H. tvaerminnensis* genome with ambiguity codes. Single copy ortholog genes were extracted from the alternative references using gff2fasta.pl and concatenated into a single sequence. The larger alignment of these individual samples was filtered for four-fold degenerate sites with MEGA v.7.16.0617 (Kumar et al., 2016). A Bayesian phylogenetic analysis was conducted using BEAST2 v.2.5.1 (Bouckaert et al., 2019; Drummond et al., 2005) with the ambiguity code option enabled. The obtained tree was combined with each sample’s geographical coordinates using the R package *phytools* v.0.7-20 (Revell, 2012). Similarly, for the host–parasite co-phylogeny, we applied the same methodology using the host genome (V3.0; Daphnia Genome Consortium).

### Demographic history

We attempted to reconstruct the demographic history of the *H. tvaerminnensis* and *H. magnivora* samples by characterizing the unfolded site frequency spectrum (uSFS) of single copy orthologs, distinguishing the uSFS for synonymous and nonsynonymous sites. The script siteFrequencySpectrum.R (https://github.com/tatumdmortimer/popgen-stats) was used to interface with *PopGenome* and modified to compute the uSFS for both site classes. In the uSFS of *H. tvaerminnensis*, we used *H. magnivora* as an outgroup to determine the state of the variants (ancestral or derived), and vice versa. We used the function gap.barplot from the R package *plotrix* v.3.7-7 (Lemon, 2006) to plot the uSFS.

To estimate changes in the effective population size history of *Hamiltosporidium* samples, we used PSMC v.0.6.5 (H. Li & Durbin, 2011). Therefore, all BAM files were downsampled to an average whole-genome coverage of 50× with the SAMtools function view. Afterwards, one consensus sequence per sample was produced using SAMtools option mpileup, BCFtools v.1.9 option call (H. Li, 2011) and vcfutils.pl vcf2fq, which is included in BCFtools. SNPs with a coverage of less than a third or more than double of the average whole-genome coverage were filtered as suggested by the developer (https://github.com/lh3/psmc). The script plotPsmc.r (Liu & Hansen, 2017) was used for plotting the inferred historical dynamics of *N_e_*. Similarly, “pseudo-diploid” samples were analyzed. Therefore, downsampled BAM files of two samples were jointly input to SAMtools to produce two-sample VCF files using BCFtools. WhatsHap v.0.18 (Martin et al., 2016) was then used for read-based phasing. Unphased sites were masked and haplotypes were combined in all four possible ways before converting them to four FASTQ files using vcf2fq.

### Rate of adaptive nucleotide substitutions

The magnitude of positive selection, or the rate of adaptive substitution (*α*), was inferred for *H. tvaerminnensis* by using both the classical McDonald–Kreitman test (MKT; McDonald & Kreitman, 1991) and more recently derived extensions. Thereby, intraspecific diversity and divergence counts as compared to the outgroup (i.e., counts of polymorphisms segregating within the species and counts of fixed differences between the ingroup and the outgroup) were created from a multi-sequence alignment of single copy orthologs. These alignments were first concatenated and then used as input for iMKT v.0.2 (Murga-Moreno et al., 2019), a web-based application (accessed on 05-21-2021) to perform standard MKTs (McDonald & Kreitman, 1991). We also used iMKT to estimate *α* based on the asymptotic MKT approach of Haller and Messer (2017). This approach differs from a traditional MKT in that it makes an explicit attempt to control for the confounding effects of low frequency, deleterious allele classes. Finally, we estimated the distribution of fitness effects to acquire a model-based estimate of *α* and *ω_A_* (an estimate of adaptation that may sometimes be more useful than *α*) using dfe-alpha v.2.16 (Eyre-Walker & Keightley, 2009; Keightley & Eyre-Walker, 2007), after masking internal stop codons with MACSE v.2.03 (Ranwez et al., 2011). We ran dfe-alpha using a wrapper script adapted from Fields et al. (in review).

## Results

### *A reference genome of* H. tvaerminnensis *based on long reads*

As the tissue of the host and the intracellular parasite are difficult to separate, we sequenced host and *H. tvaerminnensis* together using long-read PacBio technology. Our initial assembly of the host and parasite genomes resulted in an assembly length of 123 megabase pair (Mbp). While the host genome had an average coverage < 50×, the parasite genome had a coverage mostly > 100× (Fig. S1A). The *H. tvaerminnensis* genome had an assembly length of 28.5 Mbp and showed nearly 0× coverage when we analyzed reads from a sequenced *D. magna* host known to be uninfected by *H. tvaerminnensis* (Fig. S1B). We used purge_haplotigs to compile our final, haploid assembly of *H. tvaerminnensis*.

This new reference genome significantly improved the contiguity of the previous *H. tvaerminnensis* genome, which was based solely on short-read Illumina sequencing (Haag et al., 2020) and which had a total length of 18.3 Mbp, a maximum contig length of 0.06 Mbp, an N50 of 0.01 Mbp, and was composed of 2915 contigs. In contrast, our new long-read based assembly had a total length of 21.6 Mbp, a max contig length of 2.7 Mbp, an N50 of 1.03 Mbp, and 27 contigs. Along with the improved contiguity, the biological completeness of the new reference genome improved, with a ~15 % increase in both genome length and BUSCO completeness score (Table S1). However, without cytological data, this improved assembly may still not represent complete chromosomal contigs.

### Samples, mapping, and sequence variation

Infected clonal host lines were sequenced together with the parasites, resulting in 15 genomic samples, each from a different Eurasian population with sufficient representation of *H. tvaerminnensis* reads to describe its genomic variation (Table 1). Genomic samples from three populations of our outgroup *H. magnivora* were also included to extend the number of analytical approaches for characterizing genome-wide variation in the *H. tvaerminnensis* sample. The percentage of sequencing reads mapped to the parasite’s new reference genome ranged from 6.28 % to 63.70 %; most of the remaining reads were from the *D. magna* host. The average whole-genome coverage was greater than 94× in all cases of *Hamiltosporidium* infections. The overall SNP density in *H. tvaerminnensis* was 25 SNPs per kilobase pair (Kbp), with the total number of SNPs being 554,935. The average whole-genome π (Nei & Li, 1979) value was 0.009, or approximately 1 %.

### Population structure and phylogenetic analyses

We assessed the amount of genomic variation that differed or was shared between samples by applying a PCA to the whole-genome SNP data (Table S2). The first two eigenvectors of the PCA (Fig. S2) divided the *Hamiltosporidium* samples into three clusters: PC1 (explaining 36.09 % of the variance) separated *H. tvaerminnensis* samples from those of *H. magnivora*; and PC2 (22.35 %) split *H. tvaerminnensis* samples into two clusters, one Northern (N = 9) encompassing Scandinavian and East Asian samples, and one Southern (N = 6), encompassing all Mediterranean samples. A cluster analysis applied to the whole-genome SNP data resulted in the same groupings (Fig. S2), as did a Bayesian phylogenetic tree estimation based on four-fold degenerate sites of the 2,229 single copy orthologs (Fig. 1). After finding these groupings consistently in all of our population structure analyses, we refer to them as Northern and Southern *H. tvaerminnensis* lineages.

**Fig. 1:**
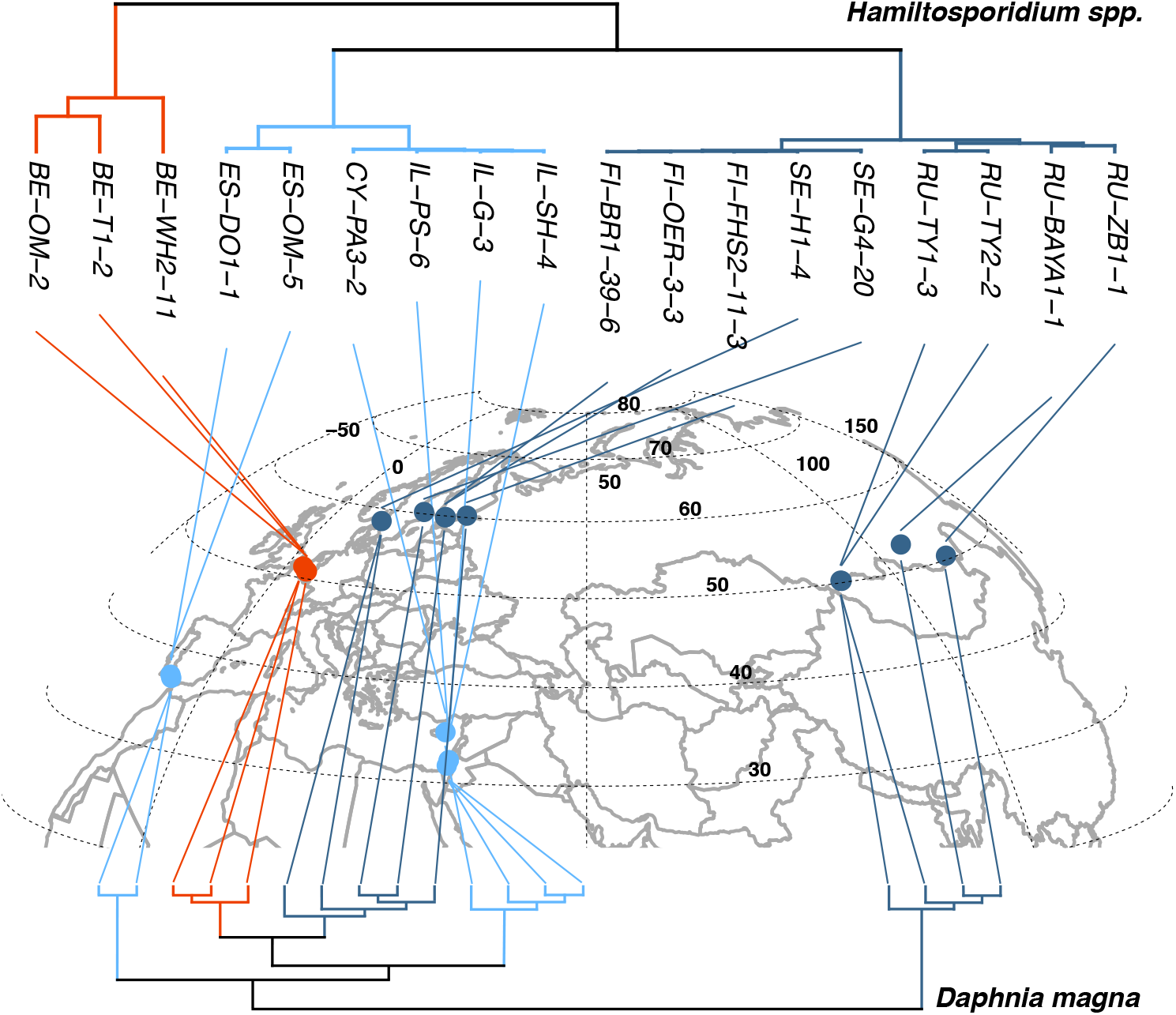
Geophylogeny of *Hamiltosporidium* spp. with co-phylogeny to the *D. magna* host. Both phylogenetic trees are based on four-fold degenerate sites but are not on the same scale. Colored lines indicate the main clusters derived from our population structure analyses: *H. magnivora* (orange), Northern *H. tvaerminnensis* (dark blue), Southern *H. tvaerminnensis* (cyan), and connect the tree tips of *Hamiltosporidium* spp. (above) and *D. magna* (below) to the sampling locality. Eurasia appears as an orthographic projection with numbers indicating latitudes and longitudes. Host and *H. tvaerminnensis* phylogenetic trees are mostly congruent, except that East Asian samples of the parasite cluster with Northern European samples rather than forming a separate clade as in the host.

We inferred the demographic history for the two *H. tvaerminnensis* lineages separately. The SNP density and the π value showed that the Northern lineage (11 SNPs per Kbp) was considerably less SNP-dense and less diverse (π = 0.005) than the Southern lineage (19 SNPs per Kbp; π = 0.008; Wilcoxon signed rank test: *p* < 0.001). We found a positive correlation between pairwise differentiation and geographical distance for the overall *H. tvaerminnensis* data and the species’ lineages using distance-based Moran’s eigenvector maps (dbMEM) analysis by redundancy analysis (RDA) (overall: *R^2^* = 0.15, *p* = 0.197; Northern: *R^2^* = 0.43, *p* = 0.001; Southern: *R^2^* = 0.82, *p* = 0.003), which has been suggested as preferable over the formerly used Mantel tests (Legendre et al., 2015). The much higher correlation coefficients within the lineages, as compared to the total dataset, strongly underline the importance of considering the phylogeography within the lineages.

### Demographic history

We examined the demographic history of *H. tvaerminnensis* looking at both the full dataset and each the Northern and Southern lineages individually. The unfolded site frequency spectra (uSFS) of *H. tvaerminnensis* suggest strong deviation from what one would expect in a Wright-Fisher population (Crawford & Lazzaro, 2012), i.e., a panmictic population in mutation-drift balance (Fig. 2A, C, and D). Specifically, we do not observe any left-skewed distribution in *H. tvaerminnensis*, although it was seen in *H. magnivora* (Fig. 2B). The Northern *H. tvaerminnensis* lineage had a clear peak around intermediate frequencies consistent with the earlier observation of fixed heterozygosity at many northern European sample sites (Haag et al., 2013b). Such a clear peak was not observed in the Southern *H. tvaerminnensis* lineage, however, which showed a more even spread than expected from a Wright-Fisher population.

**Fig. 2:**
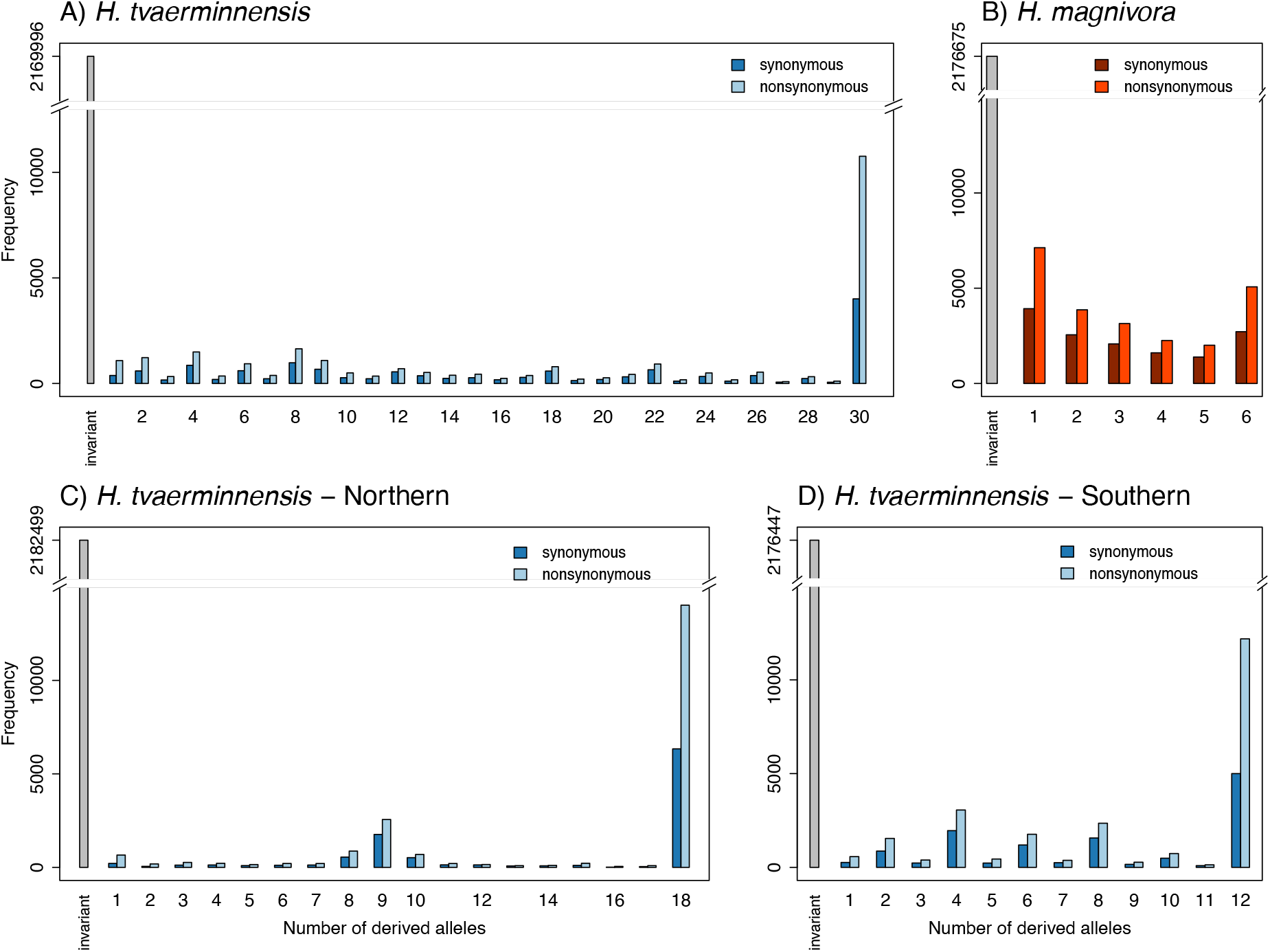
Unfolded site frequency spectra of (A) all *H. tvaerminnensis* samples, (B) all *H. magnivora* samples, (C) the Northern and (D) the Southern *H. tvaerminnensis* lineage. Distribution of the allele frequencies in the single copy orthologs is shown after identifying each position as either ancestral, i.e., invariant to the outgroup, or derived, i.e., different form the outgroup. Non-/synonymous substitutions are shown separately. Of particular interest is the peak at intermediate frequencies in the Northern lineage, as it represents mostly fixed heterozygote sites originating from the transition from sexual to asexual reproduction.

To further understand the evolution of these lineages, we analyzed the changes of *N_e_* across time using software based on a Pairwise Sequentially Markovian Coalescent (PSMC) model to infer population history. Additionally, we did a very rough time calibration using publicly available estimates of mutation rates from the well-studied ‘red yeast’ fungi *Rhodotorula toruloides* (Long et al., 2016) and previously reported generation times of microsporidia (Kramer, 1965). PSMC revealed a relatively consistent pattern across samples from the same species and lineage (Fig. 3). Ancient *N_e_*, (i.e., around 100 thousand (K) to 1 million (M) years ago) was similar among all species and lineages but diverged in more recent times, with *N_e_* for *H. magnivora* being less dynamic across time than *N_e_* for all *H. tvaerminnensis* samples. The biggest difference between the species was the recent, strong reduction in *N_e_* a few thousand years ago, seen only in the *H. tvaerminnensis* samples. There were also differences among the *H. tvaerminnensis* samples. Specifically, a more ancient decrease in *N_e_*, between around 15 K and 50 K years ago was seen in all *H. tvaerminnensis* samples except those from Southeast Europe/Middle East, suggesting distinct population histories for the eastern and western samples of the Southern lineage. Apart from that, the consistency within the Northern lineage and the two groups of the Southern lineage gives us confidence about the quality of this population history measure.

**Fig. 3:**
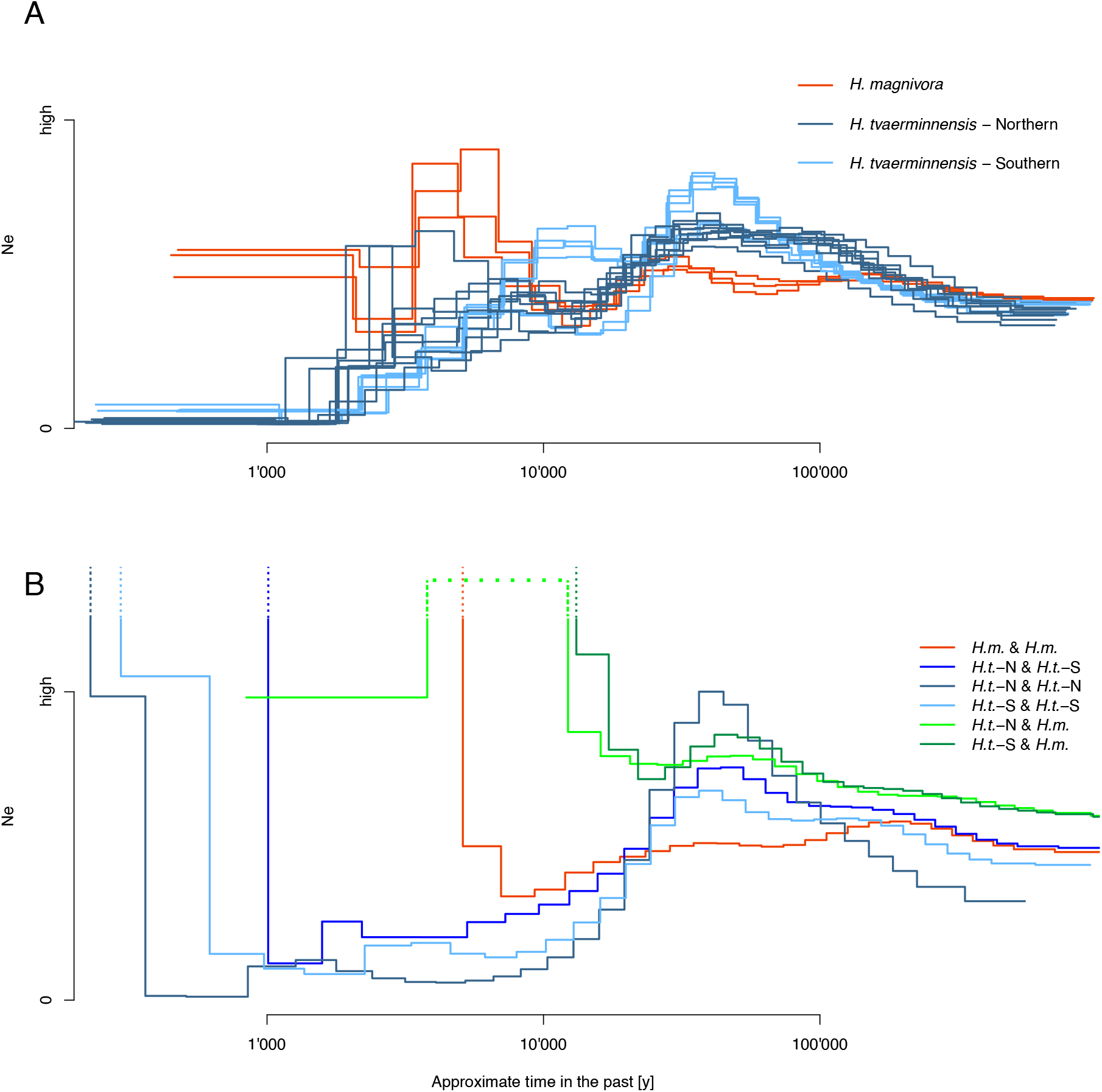
PSMC results of *Hamiltosporidium* samples. The effective population size history of *Hamiltosporidium* samples estimated using PSMC (A). Each line represents one sample, with colors indicating its cluster affiliation according to our population structure analyses. Samples from Spain (Southern *H. tvaerminnensis* cluster) show decreases in *N_e_* compared with the other samples from the Southern *H. tvaerminnensis* cluster, which are stable over the same time period. A base-10 log scale is used for the x-axis, which is time calibrated using a rough approximation. No extreme upward vertical trajectories are observed in contrast to (B), which shows the PSMC results of *Hamiltosporidium* “pseudo-diploid” samples. In (B), each line represents one *in silico*, “pseudo-diploid” sample, i.e., single phased haplotypes of our diploid samples combined in a pairwise manner. Orange indicates an intraspecies combination of *H. magnivora* (*H.m.*) haplotypes; blue shows intraspecies combinations of *H. tvaerminnensis* (*H.t.*) haplotypes, and green depicts interspecies combinations of *H. tvaerminnensis* (*H.t.*) haplotypes and *H. magnivora* (*H.m.*) haplotypes. Here, the Northern *H. tvaerminnensis* lineage is abbreviated as *H.t*.-N, and the Southern *H. tvaerminnensis* lineage as *H.t*.-S. The *H. tvaerminnensis* intraspecies combinations do not show extreme upward vertical trajectories until very recently, which suggests a rather young species rather than a diploid species that has been reproducing asexually for a long time. Only one “pseudo-diploid” sample turns downwards after an extreme upward vertical trajectory, the interspecies haplotype combination of Northern *H. tvaerminnensis* and *H. magnivora* haplotypes, which suggests gene flow between the two. This is indicated by the dotted light green line, which is drawn out of scale for the ease of presentation. The axes cover the same ranges in both graphs.

According to the PSMC results, neither *Hamiltosporidium* species showed extreme upward vertical trajectories in *N_e_*, which one would expect in a diploid species that reproduces asexually over a long time period and has non-recombining chromosomes that separately accumulate mutations (i.e., the Meselson effect; Ceplitis, 2003; Weir et al., 2016). This result could suggest that the supposedly asexual *H. tvaerminnensis* experiences some recombination, as is believed to be true for the sexual *H. magnivora*. To further understand how mode of reproduction evolved, we ran the PSMC method with *in silico*, “pseudo-diploid” samples, i.e., single phased haplotypes of our diploid samples that we combined in a pairwise manner. While this method is frequently used to estimate the relative divergence times of species (Cahill et al., 2016; Mattle-Greminger et al., 2018), we used it to investigate the degree of separate chromosomal evolution within and between *Hamiltosporidium* species. The PMSC results using “pseudo-diploids” differed strongly from the results using the original samples (compare Fig. 3A and B), especially in recent times. *N_e_* in ancient times was similar in both analyses, with none of the intraspecies “pseudo-diploid” samples’ PSMC results showing extreme upward vertical trajectories; however, in recent times, *N_e_* showed extreme upward vertical trajectories in all the intraspecies “pseudo-diploid” samples (Fig. 3B). This confutes long-time asexuality and represents spatially based population differentiation. Of the intraspecies combinations, *H. magnivora* showed the deepest upward vertical trajectories in time (orange in Fig. 3B), suggesting the highest differentiation (see also tree in Fig. 1); however, upward vertical trajectories in inter-lineage and interspecies “pseudo-diploid” samples are further in the past (green lines in Fig. 3B), which is expected for more diverged haplotype combinations. Interestingly, there may have been lineage-specific events, including hybridization with *H. magnivora* (light green in Fig. 3B): in contrast to the other “pseudo-diploid” haplotype combinations, the PSMC results of Northern *H. tvaerminnensis–H. magnivora* “pseudo-diploid” haplotype combinations show upward vertical trajectories in *N_e_* followed by a return to lower values, which we interpret as putative gene flow between the species.

### Rate of adaptive nucleotide substitutions

As genomic signals of demography and selection are often hard to disentangle, we assessed the importance of selection in *H. tvaerminnensis* by estimating the rate of adaptive substitutions (*α*) using a standard McDonald–Kreitman test (MKT). The *α* values of *H. tvaerminnensis* ranged from 0.121 to 0.227 (Table 2). Asymptotic MKT derived *α* values, which are expected to be less affected by demographic factors, ranged from 0.222 to 0.279. However, the asymptotic methodology needed more data in some frequency categories than was available in our study (i.e., more neutral and selected diversity) to perform the asymptotic fit over the data (Jesús Murga-Moreno, pers. comm.). Therefore, the estimation of *α* using asymptotic MKT resulted in large confidence intervals and was unfeasible for the Northern lineage. The *α* values of *H. tvaerminnensis* estimated using dfe-alpha ranged from −0.026 to 0.203. Estimates of *α* have previously been found to be above 0.5 in many species and to reach up to 0.9 (Galtier, 2016). The consistently low estimates of *α* for *H. tvaerminnensis* support previous findings that genomic variation in *Hamiltosporidium* spp. might be predominantly driven by nonadaptive processes rather than by adaptive evolution (Haag et al., 2020).

**Table 2:** Proportion of adaptive amino-acid substitutions (*α*) estimated for different *H. tvaerminnensis* lineages with McDonald–Kreitman tests (MKT) and dfe-alpha. *α* derived from Standard MKT presented with p-values (Fisher’s exact test), and *α* derived from dfe-alpha with a CI derived from 1,000 bootstrap samples. Rate of adaptive divergence relative to neutral divergence (*ω_A_*), also with a CI derived from 1,000 bootstrap samples, was estimated using dfe-alpha.

| Lineage | Standard MKT |  |  |  |  |  |
| --- | --- | --- | --- | --- | --- | --- |
| | $\alpha$ | P-value | | | | |
| *H. tvaerminnensis* - all | 0.227 | <0.001 |  |  |  |  |
| *H. tvaerminnensis* - north | 0.121 | <0.001 |  |  |  |  |
| *H. tvaerminnensis* - south | 0.202 | <0.001 |  |  |  |  |

| Lineage | dfe-alpha |  |  |  |  |  |
| --- | --- | --- | --- | --- | --- | --- |
| | $\alpha$ | $\alpha$ low | $\alpha$ high | $\omega_A$ | $\omega_A$ low | $\omega_A$ high |
| *H. tvaerminnensis* - all | 0.203 | 0.142 | 0.272 | 0.066 | 0.041 | 0.093 |
| *H. tvaerminnensis* - north | -0.026 | -0.104 | 0.061 | -0.008 | -0.030 | 0.021 |
| *H. tvaerminnensis* - south | 0.139 | 0.004 | 0.156 | 0.047 | 0.001 | 0.054 |

## Discussion

A core aim of population genetics is to understand genetic variation over space and time and link it to variation in the ecology of the species. To understand genetic variation in parasites, it is further helpful to know the host’s biology and features that determine interactions, such as parasite transmission and host–parasite coevolution. Our population genomic study provides insights into the variation of genomic diversity, population structure, and demographic history in a highly specific parasite that exclusively infects the planktonic freshwater species *Daphnia magna*, a model system in host-parasite evolutionary ecology. This parasite, the microsporidium *Hamiltosporidium*, is widely spread and locally very common (Decaestecker et al., 2005; Ebert et al., 2001; Goren & Ben-Ami, 2013). Our analyses use samples spanning the Eurasian region of the host’s Holarctic range where *D. magna* forms two clades, a Western Eurasian and an East Asian clade (Fields et al., 2018). We also find two clades for *H. tvaerminnensis*, but the host and parasite clades are not congruent to each other, thus are not the result of co-cladogenesis. Furthermore, we find relatively low levels of adaptive evolution, which is consistent with the hypothesis that a genome expansion may have resulted from high levels of genetic drift, corroborating a mechanism for the evolution of highly divergent genome architectures in microsporidia in general (Haag et al., 2020). Within the genus, we hypothesize that *H. tvaerminnensis* might be derived from *H. magnivora* (de Albuquerque et al., 2020), as *H. tvaerminnensis*, which presumably no longer depends on a second host, has been able to colonize geographic regions inaccessible to *H. magnivora*. This apparently became possible with the evolution of direct horizontal transmission among *D. magna* individuals (only vertical in *H. magnivora*), thus eliminating the need for a second host (*H. magnivora* is still believed to depend on a second, yet unknown, host for horizontal transmission). Living without a second host, also resulted in a loss of sexual reproduction (Fig. 4), which has been suggested to take place in the second host (Mangin et al., 1995), with profound consequences for the evolution of genetic diversity of *H. tvaerminnensis*.

**Fig. 4:**
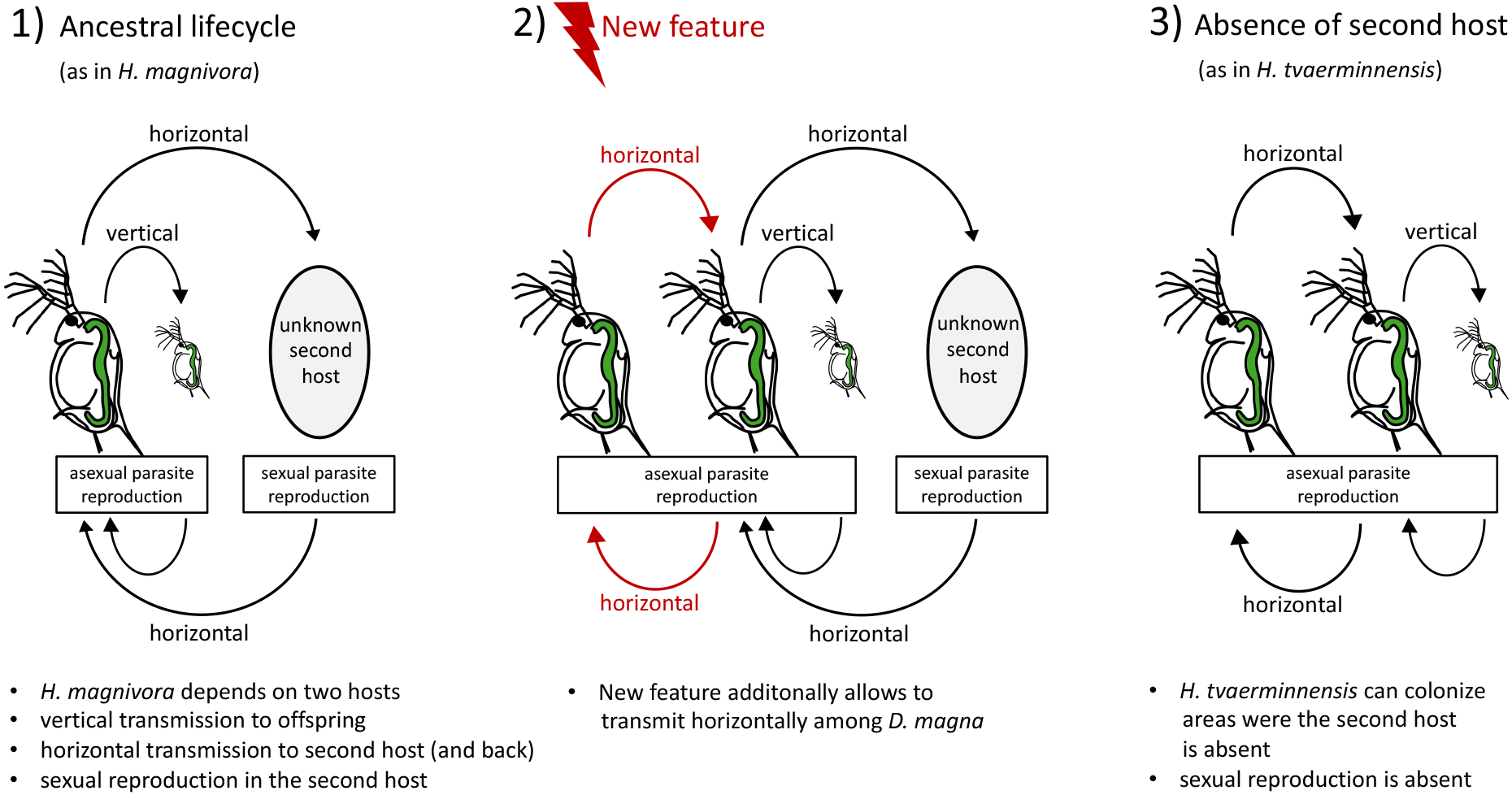
Putative evolutionary change in the *Hamiltosporidium* lifecycle. 1) The ancestral lifecycle of *Hamiltosporidium* might have involved asexual vertical transmission to offspring and transmission to a second host where sexual reproduction could take place, as is the case for *H. magnivora*. 2) However, a novel transmission strategy arose wherein horizontal transmission amongst the host *D. magna* without sexual reproduction 3) allowing this derived *Hamiltosporidium* group to colonize geographical regions where the second host is absent, as is the case for *H. tvaerminnensis*. Sexual reproduction would then only be possible in the presence of the now facultative second host.

### No pattern of host-parasite (co-)phylogeography

Because microsporidian parasites and their hosts interact closely, it might be assumed that parasite and host phylogeography should be congruent—the within species equivalent of Fahrenholz rule (Fahrenholz, 1913), which states that host and parasite phylogenies are expected to be congruent to each other, i.e., show a pattern of co-cladogenesis. It is now known, however, that this is rarely the case (Page, 2003). A strict co-phylogeography may be expected if the parasite’s mode of transmission is uniquely vertical (Werren et al., 2008) and when host and parasite co-disperse (Page, 2003). A previous attempt by Pelin et al. (2015) to reconstruct a phylogeography based on whole-genome data for the microsporidium *N. ceranae* remained inconclusive due to high, human-driven migration and lack of isolation-by-distance (IBD). In a different parasite of *D. magna*, the horizontally transmitted bacterium *Pasteuria ramosa*, a pattern of IBD was observed (Andras et al., 2018; Fields et al., 2015), but the issue of co-phylogeography was not addressed in the *Pasteuria–Daphnia* system. In the *H. tvaerminnensis–D. magna* system studied here, vertical transmission is common, and natural dispersal as well as co-dispersal seems likely (Haag et al., 2013b), especially because the unstable pond habitat of *D. magna* requires a high propensity for migration. We therefore hypothesized that we would find a pattern of host–parasite co-phylogeography and co-cladogenesis. Although *D. magna* shows two main clades in Eurasia, a Western Eurasian and an East Asian clade (Fields et al., 2018), our phylogeographic analysis suggests that the split between the two main lineages of *H. tvaerminnensis* is clearly distinct from that of the host. Instead of a Western Eurasia/Eastern Asia split, we found a North–South split, a signal consistent across all the phylogeographic analyses we conducted (Fig. 1, Fig. S2). Thus, the phylogeography of *D. magna* and *H. tvaerminnensis* is non-overlapping and cannot be explained by vertical transmission and co-dispersal alone. Nevertheless, a pattern of IBD (as has been shown for *D. magna* by Fields et al. (2015) and Andras et al. (2018)) was observed for *H. tvaerminnensis*, supporting the assumption of low or moderate gene flow between populations and reinforcing the intimate host–parasite association.

### The divergence of Northern and Southern parasite lineages

Our phylogenetic analysis of *Hamiltosporidium* revealed a clear separation between a Northern and a Southern clade of *H. tvaerminnensis*. Not surprisingly, therefore, the correlation between pairwise genetic differentiation and geographical distance was clearer within these two parasite lineages than in the combined dataset. We speculate that the Northern *H. tvaerminnensis* lineage resulted from the parasite spreading after the host expanded northwards out of its glacial refugium about 10,000 years ago when the ice retreated from the North. This spread of the Northern lineage of *H. tvaerminnensis* might have accompanied a population bottleneck, which resulted in reduced genetic diversity. Furthermore, the uSFS of Northern *H. tvaerminnensis* indicates fixed heterozygosity at many positions (Fig. 2C), a finding consistent with Haag et al.’s (2013b) suggestions that the spread to the North coincided with the emergence of asexuality and a subsequent population expansion. The Southern lineage’s samples, in comparison, are more diverged from each other, even though their currently known geographic spread is smaller than that of the Northern lineage. This could have resulted from an older transition to asexual reproduction or an additional glacial refugium other than the one in Southeast Europe/Middle-East for *D. magna* (Fields et al., 2018). However, this needs further investigation. Additionally, estimating divergence times between the parasite lineages and comparing them to the host clades’ divergence time could indicate whether co-dispersal, i.e., range expansion of host and parasite at a similar time and in the same direction, played an essential role for the evolution of the host and *H. tvaerminnensis*.

### Relative importance of selection

Models to determine the demographic history of a population are typically based on the assumption of neutrality, expecting that deviations from neutrality will lead to biased results (Johri et al., 2021). Discussing demographic history with regard to selection is therefore important. In *Hamiltosporidium* spp. It has been proposed that nonadaptive processes reduce the efficiency of adaptive evolution (Haag et al., 2020). Briefly, horizontal transmission is thought to have been the transmission mode of the *Hamiltosporidium* ancestors until, at some point, an ancestral lineage developed the ability to transmit vertically among *D. magna* as well, which is now the case for all *Hamiltosporidium* species (Haag et al., 2020). A shift from horizontal transmission to mixed-mode transmission, especially if vertical transmission predominated and played an important role in co-dispersal, could have reduced effective population size, thus making it easier for slightly deleterious mutations to increase in frequency by genomic drift and making purifying and positive selection less efficient (Haag et al., 2020). Consequently, the number of nonsynonymous substitutions fixed by genomic drift increases while the proportion of substitutions fixed by positive selection (*α*) decreases.

To test this theory, we used our whole-genome data to determine the relative strength of selection in *Hamiltosporidium*. Of the different methods used to estimate *α* in our study, dfe-alpha relies on a specific model of demography, any deviation from which might confound this method. The SFS, as well as historical changes in *N_e_* were inferred (see section below; Fig. 2 and Fig. 3) to determine how the demography of the *Hamiltosporidium* samples could violate dfe-alpha’s underlying demographic model. Indeed, Northern *H. tvaerminnensis* samples exhibited a clear recent bottleneck in their SFS and demographic history, and the estimated *α* value of this lineage was negative (Table 2), suggesting that (a) positive selection is either weak or constrained, or (b) the underlying likelihood model of dfe-alpha was not appropriate for this data. Test and reference regions were similarly affected by demography, as the synonymous and nonsynonymous SFS have the same shape (Fig. 2). To ensure a demographically robust analysis for our estimation of *α*, we conducted both a standard MKT and a more recent approach called asymptotic MKT (Haller & Messer, 2017). It has been suggested that this latter approach can accommodate a wider range of demographic histories by accounting for the effects of low frequency, deleterious variants. *α* values estimated using MKTs were similar to those using dfe-alpha. Also, the overall dataset always had the highest estimate, while the Northern lineage always had the lowest. Furthermore, among all the species for which *α* and the rate of adaptive divergence relative to neutral divergence (*ω_A_*), has been estimated, *H. tvaerminnensis* is on the lower end of the selection efficacy distribution (e.g., Galtier, 2016; Rousselle et al., 2020). Taken together, our findings support the hypothesis of weak selection efficacy, increased genetic drift, and a resulting genome expansion in *Hamiltosporidium* spp. proposed by Haag et al. (2020).

### Historical changes in N_e_

Historical processes are a major contributor to present-day distribution of genetic variation, influencing both similarities and differences in the co-phylogeography that a parasite might share with its host. The effective population size history of *H. tvaerminnensis* constructed with the PSMC method showed a rather consistent pattern across samples from the same lineage but revealed differences among lineages (Fig. 3A). Since the mutation rates and generation times of *H. tvaerminnensis* and *H. magnivora* are unknown, we used available estimates for related species to estimate time intervals of interest. The more ancient decrease in *N_e_*, which is observable in many species (Hewitt, 1996; Holder et al., 1999; Stewart et al., 2010), is thought to stem from the species’ range contraction, which happened when the glaciers expanded at the beginning of the last glaciation about 110,000 years ago. It is followed by an increase in *N_e_*, presumably after the glaciers retreated about 10,000 years ago, as also described for the host *D. magna* (Fields et al., 2018). Samples that originated from hosts in the Middle East, a region where glaciers had no direct impact and where the presumed glacial refugia of the Western Eurasian *D. magna* clade was located, consistently seem to be more stable during the whole period of glacial expansion and recession. Numerous other species have shown patterns consistent with southern refugia during this period (Stewart et al., 2010). Importantly, our Spanish samples of the Southern lineage show an *N_e_* pattern of change that is more similar to non-Middle Eastern samples (Fig. 3A), coinciding with the absence of a glacial refugium for hosts in southwestern Eurasia (Fields et al., 2018). Thus, *Daphnia* hosts and their parasites presumably recolonized Western Europe from the same refugium, including the Hispanic peninsula after the last glaciation.

A second decrease where *N_e_* dips to very low estimates is observed in more recent times for all *H. tvaerminnensis* samples but not for *H. magnivora*. A number of potential factors can decrease the effective population size: specialization of the parasite to a single host species; a clonal mode of reproduction (i.e. a switch to asexuality); an epidemic dynamic of boom and bust; restricted and local dispersal; frequent population extinctions and recolonizations; a short-lived, annual or ephemeral host; and small, fragmented host populations (Barrett et al., 2008). Although several of these factors likely contributed to the *N_e_* dynamics of *H. tvaerminnensis*, it is unclear which of them predominated. For example, *D. magna* varies strongly in susceptibility to *H. tvaerminnensis*, with the most susceptible hosts occurring in unstable habitats that have a high propensity to dry up in summer (like ephemeral rock or desert pools) (Cabalzar et al., 2019; Lange et al., 2015). These unstable habitats are also associated with frequent population extinctions and recolonizations as well as a loss of diversity because of frequent population bottlenecks.

However, as previous analyses have suggested that *H. tvaerminnensis* transitioned from a sexual to an asexual reproduction mode at some point in its recent history (Haag et al., 2013a), we hypothesize that the more recent decrease in *H. tvaerminnensis’ N_e_* might be related mainly to the evolution of horizontal transmission directly from *D. magna* to *D. magna* (Fig. 4). The loss of sexual reproduction in the absence of a second host, which is believed to be necessary for sexual reproduction in *H. magnivora* (Mangin et al., 1995), reduced *N_e_*. However, the transition from sexual to exclusively asexual reproduction would leave a very distinct historical pattern of change in *N_e_* as inferred via PSMC. Specifically, because individual homologous chromosomes can no longer recombine, one would expect to see an upward, nearly vertical trajectory in *N_e_* as mutations accumulate on separate chromosomal copies, which our reconstructed history of *H. tvaerminnensis*’ population size does not show, or at least not deep enough in time when such a transition would be easier to pin down with the PSMC method. This implies that exclusive asexuality is probably not ancient, coinciding with the SFS of the Northern lineage, which shows a distinct peak at 0.5 caused by fixed heterozygosity. Being exclusive asexual for a long period of time would provide sufficient time for individual haplotypes to accumulate their own distinct mutations, which is not what the SFS shows (Fig. 2). Alternatively, although *H. tvaerminnensis* may be a mostly asexual species, it could still show occasional recombination, for example, when it encounters its now presumably facultative second host. A loss of sexual reproduction with occasional recombination would explain the low estimates of *N_e_* without extreme upward vertical trajectories of non-recombining chromosomes, using the PSMC method. However, any discussion about the second decrease in *N_e_* should be undertaken with the mindfulness that estimates of PSMC from the recent past are generally more uncertain (H. Li & Durbin, 2011), and that PSMC has not often been used for understanding the demographic history of parasitic species (but see e.g., Hecht et al., 2018).

When comparing *H. magnivora* and *H. tvaerminnensis*, we found that their histories of *N_e_* do not align precisely. Along with differences in their demographies, differences in mutation rate and generation time, both of which used here may not be optimal for each species, can also cause aberrations with the PSMC method. The PSMC results for the *in silico*, “pseudo-diploid” samples support the idea that hybridization events happened between *H. magnivora* and the Northern *H. tvaerminnensis* lineage. The combinations of *H. tvaerminnensis* and *H. magnivora* haplotypes show the expected, nearly vertical trajectories of no longer recombining lineages in deeper time (Fig. 3B). However, combinations of the Northern *H. tvaerminnensis* lineage and *H. magnivora* reveal a demographic signal consistent with gene flow between the two species. Overall, our analyses support the suggestion that an ancestral *H. tvaerminnensis* became able to transmit horizontally directly from *D. magna* to *D. magna* (*H. magnivora* is unable to do this) and thus was able to persist in habitat without its second host. This allowed *H. tvaerminnensis* to colonize Northern habitats, but also made it reliant almost exclusively on asexual reproduction, with the latent ability to recombine if the second host were present.

### Comparatively high genomic variation in *Hamiltosporidium*

Large-sized microsporidia genomes are generally difficult to assemble due to a high number of repetitive elements (Parisot et al., 2014). *H. tvaerminnensis* has been target of previous assembly trials using shotgun and Illumina sequencing (Corradi et al., 2009; Haag et al., 2020). Here, we were able to produce a very contiguous genome assembly of this microsporidium with a large genome size using long-read PacBio sequencing. *H. tvaerminnensis* has a whole-genome SNP density of approximately 25 SNPs per Kbp, which is more diverse than its close relative *N. ceranae* (12.7) and on the high end compared to other microsporidia and fungi (Pelin et al., 2015). A likely reason why the multi-continent *N. ceranae* sample has a lower density estimate is because the species was recently introduced into the sampled region and may have experienced a bottleneck during range expansion (Pelin et al., 2015). In contrast, *D. magna*, the host of *H. tvaerminnensis*, is native to Eurasia.

## Conclusion

The evidence presented here of a non-overlapping phylogeography between two partners in an intimate host–parasite system with a wide geographic distribution has several implications. It suggests that the parasite, *H. tvaerminnensis*, might be a rather young parasite of its host, *D. magna*. Else, it could not be simultaneously wide-spread *and* have a phylogeography distinct from its host. Also, it implies that co-dispersal of host and parasite, while common, is not the only form of dispersal. Thus our research underscores the importance and potential of using samples from the whole-species range in population genomic studies. Regarding microsporidia with long genomes, we could strengthen the assumption of a reduced selection efficacy, likely as a result of high levels of genetic drift that ultimately caused genome expansion (Haag et al., 2020). By quantifying genomic variation and selection efficacy in other microsporidia, more evidence for this hypothesis and for the evolution of different genome architectures in this taxon could be provided. However, while our study reveals some general principles on the evolution of genomic variation in a parasite, it also shows how specific factors shape genomic variation, some of these (or their combination) being possibly unique to this system (e.g., loss of the second host and the shift to exclusive asexuality in the absence of the second host). It therefore provides a case study for the genomic evolution of a microsporidium at a fine scale, which awaits comparison to other systems that will help paint a bigger picture of the evolution of specific obligate parasites.

## Acknowledgements

We thank Jürgen Hottinger, Michelle Krebs and Andrea Cabalzar for help in the field and laboratory. Members of the Ebert group provided feedback on the study and the manuscript. This work was supported by the Swiss National Science Foundation (SNSF) (grant numbers 310030B_166677 and 310030_188887 to DE).

## Data accessibility

All analysis scripts as well as raw and processed data are available at https://github.com/pascalangst/Angst_etal_2021_MolEcol, NCBI SRA database (BioProject IDs XXXX), and NCBI GenBank, Zenodo XXXX (see git repo for all accessions).

## Author contributions

All authors designed the study. PA analysed the data and wrote the manuscript. All authors reviewed the manuscript.

