## Supplemental material for "Demographic history shapes genomic variation in an intracellular parasite with a wide geographic distribution"

### Supplementary figures

**Fig. S1: Blobplots for mapping and annotating contigs resulting from A) raw PacBio data used to generate the total assembly of both host and parasite DNA, and B) from Illumina data deriving from host genotype known to not be infected with *H. tvaerminnensis*.**

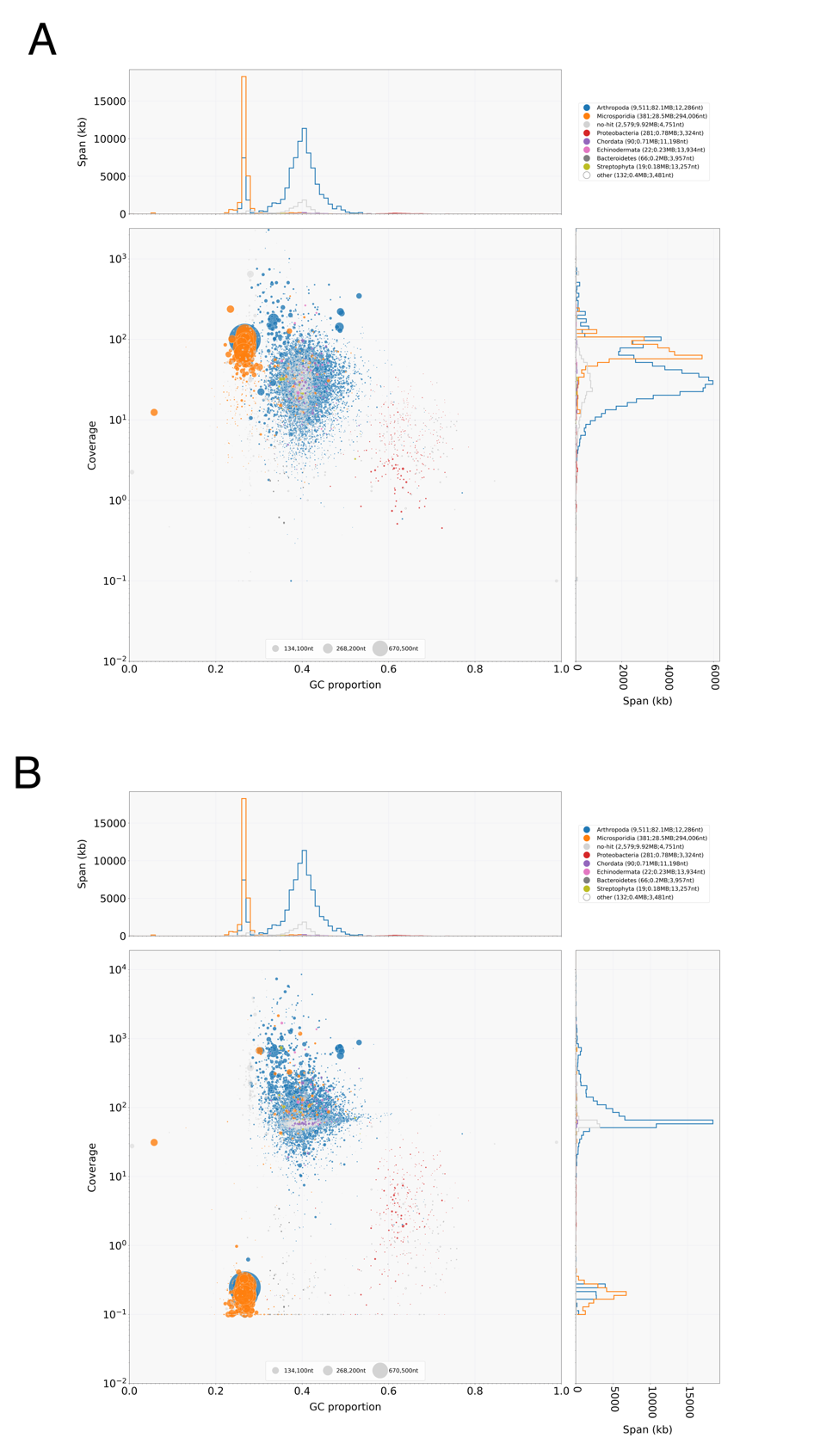

**Fig. S2: Population Structure of Hamiltosporidium.** (A) Principle component analysis (B) Cluster analysis

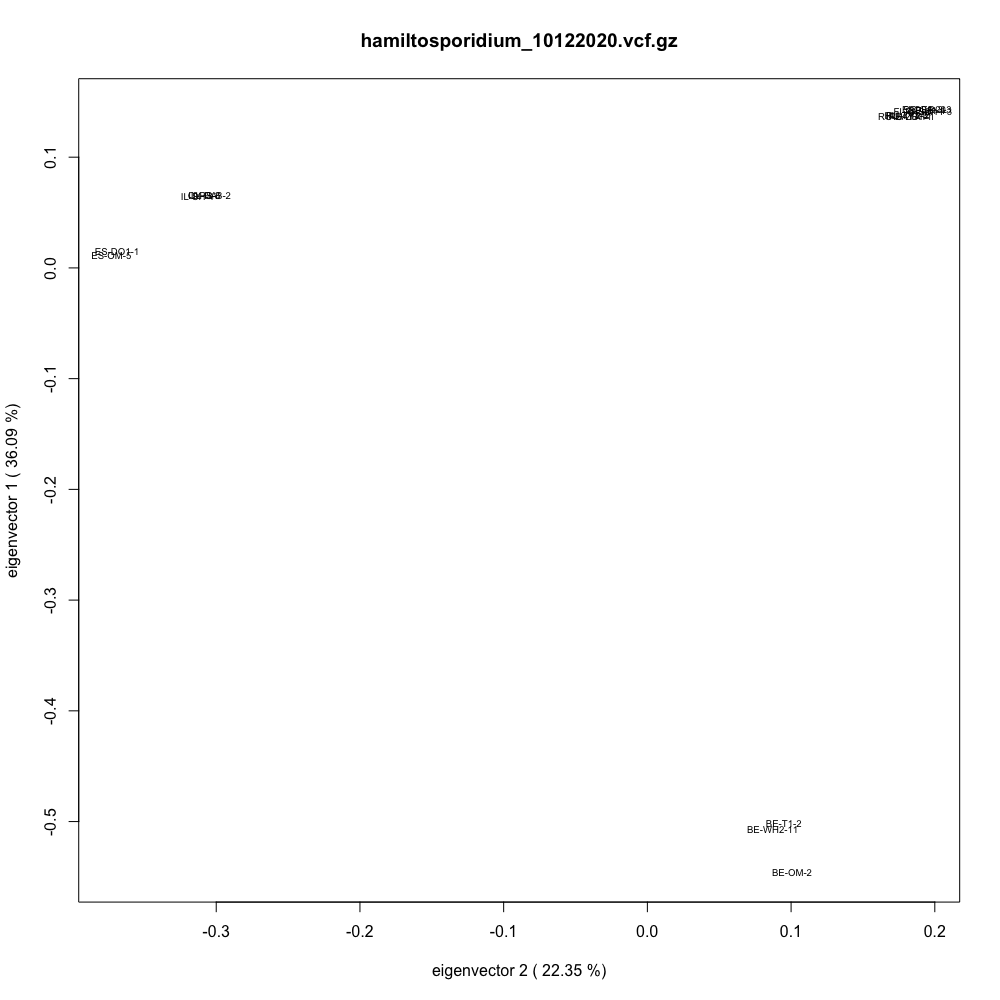

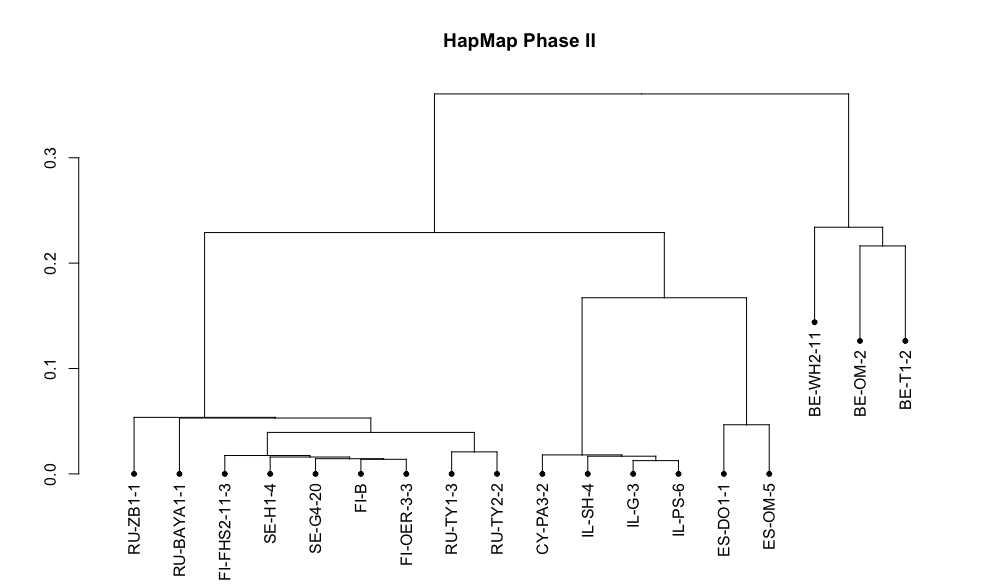

B

A

### Supplementary tables

**Table S1: BUSCO output for FIOER33 v1 assembly and new, improved FIOER33 v2 assembly.** V1 assembly is available from: NCBI database; Assembly name: FIOER33 v1; GenBank assembly accession: GCA_004325045.1, Bioprojects accession: PRJNA419750 (Haag et al., 2020).

| FIOER33 v1: | |
| --- | --- |
| C:60.4%[S:58.2%,D:2.2%],F:2.3%,M:37.3%,n:600 | |
| 362 | Complete BUSCOs (C) |
| 349 | Complete and single-copy BUSCOs (S) |
| 13 | Complete and duplicated BUSCOs (D) |
| 14 | Fragmented BUSCOs (F) |
| 224 | Missing BUSCOs (M) |
| 600 | Total BUSCO groups searched |
| FIOER33 v2: | |
| C:70.6%[S:65.8%,D:4.8%],F:3.8%,M:25.6%,n:600 | |
| 424 | Complete BUSCOs (C) |
| 395 | Complete and single-copy BUSCOs (S) |
| 29 | Complete and duplicated BUSCOs (D) |
| 23 | Fragmented BUSCOs (F) |
| 153 | Missing BUSCOs (M) |
| 600 | Total BUSCO groups searched |

**Table S2:** **List of *H. tvaerminnensis* coding sequence (CDS) alignments with single copy ortholog in *H. magnivora*.** The single copy orthologous protein of H. magnivora is denoted in the last column. Per-CDS π and π_N_/π_S_ value were calculated using selectionstats.py.

| locus_tag | π | π_N_/π_S_ | Ortholog |
| --- | --- | --- | --- |
| FUN_000007-T1 | 0.00099614 | 0.16272649 | TBU08418.1 |
| FUN_000008-T1 | 0.00492276 | 0.66321673 | TBU01812.1 |
| FUN_000009-T1 | 0.00586918 | 6.07696015 | TBT99657.1 |
| FUN_000010-T1 | 0.00207874 | 0 | TBU04614.1 |
| FUN_000012-T1 | 0.00252765 | 0.14314228 | TBU04613.1 |
| FUN_000013-T1 | 0.00397752 | 0.11333637 | TBU05979.1 |
| FUN_000014-T1 | 0.00142864 | 0.13247779 | TBU05978.1 |
| FUN_000015-T1 | 0.00159754 | 0.18325005 | TBU05977.1 |
| FUN_000016-T1 | 0.00317523 | 0.0487588 | TBU06570.1 |
| FUN_000017-T1 | 0.00300423 | 0.13346123 | TBU06571.1 |
| FUN_000018-T1 | 0.00264664 | 0.20936481 | TBU06572.1 |
| FUN_000021-T1 | 0.00040799 | Inf | TBT98599.1 |
| FUN_000022-T1 | 0.00359186 | 0.50714612 | TBU05748.1 |
| FUN_000024-T1 | 0.00431683 | 0.42217644 | TBU01508.1 |
| FUN_000025-T1 | 0.003788 | 0.04989015 | TBU01507.1 |
| FUN_000028-T1 | 0.00650875 | 0.20564388 | TBU01140.1 |
| FUN_000032-T1 | 0 | NA | TBU09119.1 |
| FUN_000034-T1 | 0.00423803 | 0.4553073 | TBU00420.1 |
| FUN_000036-T1 | 0.00360024 | 0.04676127 | TBU03543.1 |
| FUN_000037-T1 | 0.00370281 | 0.38558822 | TBU03544.1 |
| FUN_000039-T1 | 0.00179665 | 0 | TBT97492.1 |
| FUN_000059-T1 | 0.00134319 | 0 | TBU04336.1 |
| FUN_000062-T1 | 0.00831541 | 0.1700732 | TBU08648.1 |
| FUN_000063-T1 | 0.00346041 | 0.31522583 | TBU08647.1 |
| FUN_000065-T1 | 0.00159238 | 0.16891884 | TBU08646.1 |
| FUN_000067-T1 | 0 | NA | TBU07230.1 |
| FUN_000068-T1 | 0 | NA | TBU07231.1 |
| FUN_000070-T1 | 0.00086554 | 0.16050584 | TBU03170.1 |
| FUN_000071-T1 | 0.00365047 | 0.2846938 | TBU03171.1 |
| FUN_000072-T1 | 0.00205936 | 0.1262901 | TBT99595.1 |
| FUN_000073-T1 | 0.00138212 | Inf | TBT97970.1 |
| FUN_000074-T1 | 0.00075153 | Inf | TBU00970.1 |
| FUN_000075-T1 | 0.00011085 | 0 | TBU00969.1 |
| FUN_000076-T1 | 0.00090362 | 0.37480032 | TBU00968.1 |
| FUN_000083-T1 | 0.00133036 | Inf | TBU09301.1 |
| FUN_000087-T1 | 0.00027858 | 0.2787731 | TBU01787.1 |
| FUN_000089-T1 | 0 | NA | TBU02151.1 |
| FUN_000094-T1 | 0.00032749 | 0.12830655 | TBU06489.1 |
| FUN_000095-T1 | 0.00037078 | 0.09860909 | TBT99092.1 |
| FUN_000100-T1 | 0.00033602 | 0.34514385 | TBU05888.1 |
| FUN_000106-T1 | 0.00019201 | Inf | TBU04756.1 |
| FUN_000125-T1 | 0.00980947 | 0.69332909 | TBU02213.1 |
| FUN_000127-T1 | 0.0040004 | 0.05514241 | TBT99614.1 |
| FUN_000129-T1 | 0.00312644 | 0.87579077 | TBU00157.1 |
| FUN_000130-T1 | 0.00322201 | 0.44881932 | TBT98692.1 |
| FUN_000132-T1 | 0.0030722 | 1.12268357 | TBT96904.1 |
| FUN_000133-T1 | 0.0024306 | 0.07236282 | TBU04133.1 |
| FUN_000136-T1 | 0.01232384 | 0.32187936 | TBU06768.1 |
| FUN_000137-T1 | 0.00475319 | 0.28254793 | TBU07236.1 |
| FUN_000139-T1 | 0.00800942 | 0.3428881 | TBU07237.1 |
| FUN_000141-T1 | 0.00522039 | 0.27053626 | TBU05745.1 |
| FUN_000142-T1 | 0.01418619 | 1.14005439 | TBU05744.1 |
| FUN_000143-T1 | 0.00123907 | Inf | TBT98752.1 |
| FUN_000145-T1 | 0 | NA | TBU08663.1 |
| FUN_000147-T1 | 0.00232271 | 0.08739968 | TBU08665.1 |
| FUN_000151-T1 | 0.00193589 | 0.31308904 | TBU09228.1 |
| FUN_000157-T1 | 0.03594255 | 0.51194411 | TBT99902.1 |
| FUN_000164-T1 | 0.02175544 | 0.496569 | TBU08529.1 |
| FUN_000166-T1 | 0.01585508 | 0.34934563 | TBU04999.1 |
| FUN_000167-T1 | 0.0031118 | 0.47934001 | TBT96819.1 |
| FUN_000169-T1 | 0.00237953 | 0 | TBT96794.1 |
| FUN_000170-T1 | 0.0044011 | 0.23955158 | TBT98792.1 |
| FUN_000180-T1 | 0.00137368 | 0.27211915 | TBT99753.1 |
| FUN_000181-T1 | 0.00286738 | 0 | TBT99754.1 |
| FUN_000182-T1 | 0.0055001 | 0.22209006 | TBU04310.1 |
| FUN_000184-T1 | 0.02426712 | 1.2078101 | TBU04309.1 |
| FUN_000185-T1 | 0.0028921 | 0.06518211 | TBU05898.1 |
| FUN_000189-T1 | 0.00270102 | 1.31319216 | TBU08473.1 |
| FUN_000190-T1 | 0.00039459 | 0.24973288 | TBU08471.1 |
| FUN_000191-T1 | 0.00110852 | Inf | TBU00466.1 |
| FUN_000192-T1 | 0.00060089 | 0 | TBU00465.1 |
| FUN_000197-T1 | 0.00152927 | 0.9723317 | TBU03947.1 |
| FUN_000198-T1 | 0.00699785 | 0.37959774 | TBU03946.1 |
| FUN_000200-T1 | 0.0014515 | 0.06032864 | TBU03945.1 |
| FUN_000201-T1 | 0.00289728 | 0.14898088 | TBT97307.1 |
| FUN_000206-T1 | 0.00075457 | 0.29451639 | TBU08058.1 |
| FUN_000207-T1 | 0.00404649 | 0.49530241 | TBU08057.1 |
| FUN_000208-T1 | 0.00616154 | 1.52597535 | TBU08056.1 |
| FUN_000209-T1 | 0.00445152 | 0.29994556 | TBU08055.1 |
| FUN_000210-T1 | 0.03337501 | 0.81316268 | TBU08054.1 |
| FUN_000212-T1 | 0.00201779 | Inf | TBU05561.1 |
| FUN_000213-T1 | 0.00314958 | 0.74447791 | TBU05562.1 |
| FUN_000214-T1 | 0.0001593 | 0 | TBU05563.1 |
| FUN_000215-T1 | 0.00300858 | 0.33139662 | TBU05564.1 |
| FUN_000216-T1 | 0.00394053 | 0.01275948 | TBT99444.1 |
| FUN_000218-T1 | 0.00290835 | 0 | TBU00917.1 |
| FUN_000219-T1 | 0.00540644 | 0.09918531 | TBU00916.1 |
| FUN_000221-T1 | 0.00428667 | 0.09745555 | TBT98836.1 |
| FUN_000222-T1 | 0.0020795 | 0 | TBT98837.1 |
| FUN_000223-T1 | 0.00020326 | Inf | TBU03803.1 |
| FUN_000225-T1 | 0.00994135 | 0.91603679 | TBU06517.1 |
| FUN_000227-T1 | 0.00428109 | 0.14069715 | TBU03186.1 |
| FUN_000230-T1 | 0.00196698 | 0.05676448 | TBU08392.1 |
| FUN_000231-T1 | 0.00549994 | 0.40840333 | TBU08393.1 |
| FUN_000232-T1 | 0.00486037 | 1.38094659 | TBU08394.1 |
| FUN_000233-T1 | 0.00915892 | 0.15358643 | TBU08395.1 |
| FUN_000234-T1 | 0.00336479 | 0.28819431 | TBU08396.1 |
| FUN_000235-T1 | 0.00720695 | 0.20369301 | TBT99742.1 |
| FUN_000236-T1 | 0.00419681 | 0.08508727 | TBT99741.1 |
| FUN_000238-T1 | 0.00514659 | Inf | TBU06538.1 |
| FUN_000240-T1 | 0.03225806 | 0.85684647 | TBU03021.1 |
| FUN_000241-T1 | 0.0034708 | 0.41446535 | TBU01562.1 |
| FUN_000242-T1 | 0.00556083 | 0.63011797 | TBU05869.1 |
| FUN_000249-T1 | 0.0024569 | 0.32538649 | TBU02840.1 |
| FUN_000252-T1 | 0.00091107 | 0 | TBU09456.1 |
| FUN_000255-T1 | 0.00306743 | 0.12824675 | TBU09457.1 |
| FUN_000256-T1 | 0.00749616 | 2.38171552 | TBU09458.1 |
| FUN_000257-T1 | 0.0001044 | 0 | TBU09459.1 |
| FUN_000258-T1 | 0.00238575 | 0.24819126 | TBU09460.1 |
| FUN_000263-T1 | 0.0066736 | 2.95549985 | TBU04198.1 |
| FUN_000264-T1 | 0.0013244 | 0 | TBU04199.1 |
| FUN_000267-T1 | 0.00490669 | 0.01019154 | TBU04200.1 |
| FUN_000268-T1 | 0.00518141 | Inf | TBU03715.1 |
| FUN_000269-T1 | 0.00024028 | 0 | TBU05646.1 |
| FUN_000270-T1 | 0.0029899 | 0.29045795 | TBU05647.1 |
| FUN_000271-T1 | 0.00392225 | 0.14129516 | TBU05648.1 |
| FUN_000272-T1 | 0.0022633 | 0.20156142 | TBU06214.1 |
| FUN_000273-T1 | 0.01554355 | 0.64660145 | TBU06213.1 |
| FUN_000274-T1 | 0.00010314 | 0.26662167 | TBU02640.1 |
| FUN_000280-T1 | 0.00591398 | 0.20500878 | TBU07835.1 |
| FUN_000293-T1 | 0.00496848 | 0.57427144 | TBU06891.1 |
| FUN_000294-T1 | 0.00159229 | 0.16676375 | TBU06890.1 |
| FUN_000296-T1 | 0.00221529 | 0.03373448 | TBU06889.1 |
| FUN_000303-T1 | 0.00128359 | 0 | TBU00408.1 |
| FUN_000304-T1 | 0.0035195 | 0.82929336 | TBU03648.1 |
| FUN_000305-T1 | 0.00016109 | 0 | TBU03649.1 |
| FUN_000306-T1 | 0.00366388 | 0.11815898 | TBU05585.1 |
| FUN_000308-T1 | 0.01954069 | 1.16175181 | TBU05586.1 |
| FUN_000311-T1 | 0.00929624 | 2.3538586 | TBU00348.1 |
| FUN_000317-T1 | 0.01103816 | 0.28766471 | TBU06425.1 |
| FUN_000319-T1 | 0.01034959 | 0.28424885 | TBU01047.1 |
| FUN_000320-T1 | 0.00376413 | 0.07315506 | TBU04323.1 |
| FUN_000327-T1 | 0.00527728 | 1.64952218 | TBU09260.1 |
| FUN_000330-T1 | 0.00156154 | Inf | TBU03238.1 |
| FUN_000334-T1 | 0.00039279 | Inf | TBU01266.1 |
| FUN_000335-T1 | 0.00052791 | 0.04707705 | TBU01265.1 |
| FUN_000338-T1 | 0.00013399 | 0.22418601 | TBT99958.1 |
| FUN_000341-T1 | 0.00214384 | 0.15089877 | TBU00733.1 |
| FUN_000344-T1 | 0.02376093 | 0.53978046 | TBU04255.1 |
| FUN_000346-T1 | 0.00525687 | 0.66065175 | TBU08604.1 |
| FUN_000347-T1 | 0.00525235 | 0.35633792 | TBU08603.1 |
| FUN_000349-T1 | 0.00156469 | 3.06386533 | TBU02992.1 |
| FUN_000351-T1 | 0.00498406 | 0.34345164 | TBU02690.1 |
| FUN_000352-T1 | 0.00655131 | 0.34886084 | TBU02689.1 |
| FUN_000353-T1 | 0.00328554 | 0.12997324 | TBU01742.1 |
| FUN_000354-T1 | 0.00325011 | 0.30741891 | TBU01743.1 |
| FUN_000357-T1 | 0.00629426 | 0.93027208 | TBT99552.1 |
| FUN_000358-T1 | 0.00638471 | 0.32095908 | TBT99553.1 |
| FUN_000359-T1 | 0.00116995 | 0 | TBU09395.1 |
| FUN_000360-T1 | 0.00178429 | 0 | TBU09394.1 |
| FUN_000361-T1 | 0.00492057 | 0.29488154 | TBU09393.1 |
| FUN_000362-T1 | 0.00716846 | 0.39818711 | TBU09392.1 |
| FUN_000364-T1 | 0.00408941 | 0.20840681 | TBU09391.1 |
| FUN_000365-T1 | 0.0034079 | 0.2509908 | TBU09390.1 |
| FUN_000367-T1 | 0.01008269 | 0.95014337 | TBU07913.1 |
| FUN_000369-T1 | 0.00234723 | 0.17029368 | TBU07915.1 |
| FUN_000389-T1 | 0.00095902 | 0.04476185 | TBU09556.1 |
| FUN_000393-T1 | 0.00198736 | 0.24003055 | TBU03513.1 |
| FUN_000395-T1 | 0.00315598 | 0.04369714 | TBU03277.1 |
| FUN_000398-T1 | 0.00351906 | 0.1171077 | TBU00382.1 |
| FUN_000402-T1 | 0.0061634 | 0.45441507 | TBU08177.1 |
| FUN_000404-T1 | 0.00568183 | 0.78982476 | TBU04838.1 |
| FUN_000405-T1 | 0.00045966 | 0 | TBU04840.1 |
| FUN_000406-T1 | 0.00213545 | 0.13589345 | TBU00588.1 |
| FUN_000407-T1 | 0.01069101 | 0.39533599 | TBU05964.1 |
| FUN_000408-T1 | 0.0035812 | 0.21821942 | TBU05965.1 |
| FUN_000409-T1 | 0.00516401 | 0.13336415 | TBU05966.1 |
| FUN_000410-T1 | 0.00233541 | 0.10231908 | TBU05967.1 |
| FUN_000411-T1 | 0.00383076 | 0.0623098 | TBU09273.1 |
| FUN_000412-T1 | 0.00638903 | 0.46566447 | TBU09272.1 |
| FUN_000413-T1 | 0.00115251 | 0 | TBU09271.1 |
| FUN_000414-T1 | 0.00483454 | 0.47241877 | TBU09269.1 |
| FUN_000415-T1 | 0.00061659 | 0.02840171 | TBU09268.1 |
| FUN_000416-T1 | 0.00252167 | 0 | TBU09267.1 |
| FUN_000417-T1 | 0.0067352 | 0.19753372 | TBU09266.1 |
| FUN_000419-T1 | 0.00114354 | 2.02892354 | TBU03095.1 |
| FUN_000421-T1 | 0.0016129 | 0.17594787 | TBU03096.1 |
| FUN_000422-T1 | 0.00074245 | 0.14508671 | TBT98338.1 |
| FUN_000423-T1 | 0.00168067 | 0.11493076 | TBU08795.1 |
| FUN_000424-T1 | 0.00296099 | 0.33885644 | TBU08796.1 |
| FUN_000429-T1 | 0 | NA | TBU08798.1 |
| FUN_000430-T1 | 0.00022519 | 0.23491379 | TBT97522.1 |
| FUN_000431-T1 | 0.00106939 | 0.02654579 | TBU00851.1 |
| FUN_000433-T1 | 0.00011879 | Inf | TBU00850.1 |
| FUN_000434-T1 | 0.00074215 | 0.34172777 | TBU02881.1 |
| FUN_000435-T1 | 0.00093968 | 0.26120444 | TBU02880.1 |
| FUN_000437-T1 | 0.00033832 | 0.25234135 | TBU00777.1 |
| FUN_000440-T1 | 0.00049438 | 0.06945447 | TBT97228.1 |
| FUN_000442-T1 | 0.00287682 | 0.39375588 | TBU07969.1 |
| FUN_000443-T1 | 0.000812 | 0.03201069 | TBU07968.1 |
| FUN_000444-T1 | 0.0079825 | 0.13345755 | TBU07967.1 |
| FUN_000445-T1 | 0.00365451 | 0.05376677 | TBU07966.1 |
| FUN_000446-T1 | 0.00212108 | 0 | TBU07965.1 |
| FUN_000447-T1 | 0.00042426 | 0 | TBU07964.1 |
| FUN_000448-T1 | 0.00757434 | 0.2192048 | TBU06341.1 |
| FUN_000449-T1 | 0.00317292 | 0.04726629 | TBU06342.1 |
| FUN_000450-T1 | 0.00278628 | 0.19304907 | TBU06343.1 |
| FUN_000451-T1 | 0.00269045 | 0.1007992 | TBU06344.1 |
| FUN_000452-T1 | 0.00172763 | 0.18797141 | TBU06345.1 |
| FUN_000453-T1 | 0.00629621 | 0.4051346 | TBU02114.1 |
| FUN_000456-T1 | 0.00927942 | 2.01896239 | TBU01087.1 |
| FUN_000457-T1 | 0.00220773 | 1.89105781 | TBU05350.1 |
| FUN_000458-T1 | 0.00279424 | 0.11066901 | TBU05351.1 |
| FUN_000460-T1 | 0.0083476 | 2.99608303 | TBU04160.1 |
| FUN_000461-T1 | 0.00497381 | 0.57744684 | TBU04159.1 |
| FUN_000462-T1 | 0.00620065 | 0.63322336 | TBU04158.1 |
| FUN_000464-T1 | 0.00469263 | 0.1244793 | TBU06297.1 |
| FUN_000466-T1 | 0.00268817 | 0.46481277 | TBU01134.1 |
| FUN_000467-T1 | 0.00076176 | 1.54686558 | TBU01135.1 |
| FUN_000470-T1 | 0.00102407 | 0 | TBU00251.1 |
| FUN_000471-T1 | 0.00635811 | 0.39372309 | TBU08720.1 |
| FUN_000472-T1 | 0.0032129 | 0.17982227 | TBU08719.1 |
| FUN_000474-T1 | 0.00169176 | 0 | TBU08718.1 |
| FUN_000475-T1 | 0.00467989 | 0.14551447 | TBU08717.1 |
| FUN_000476-T1 | 0.00620398 | 0.20812277 | TBU08716.1 |
| FUN_000477-T1 | 0.00480287 | 0.15043668 | TBU08715.1 |
| FUN_000478-T1 | 0.00533313 | 0.12278847 | TBU08714.1 |
| FUN_000480-T1 | 0.01108899 | 0.42210972 | TBU08850.1 |
| FUN_000482-T1 | 0.00137944 | 0.08272671 | TBU08852.1 |
| FUN_000485-T1 | 0.02325015 | 1.07103227 | TBU02209.1 |
| FUN_000491-T1 | 0.00325889 | 2.95631701 | TBT98394.1 |
| FUN_000494-T1 | 0.00177653 | 0.1649695 | TBU00177.1 |
| FUN_000498-T1 | 0.01162961 | 0.16099524 | TBU09248.1 |
| FUN_000501-T1 | 0.00395217 | 0.4686792 | TBU09250.1 |
| FUN_000502-T1 | 0.00981468 | 0.53676875 | TBU09251.1 |
| FUN_000503-T1 | 0.00232122 | 0.0383018 | TBU04621.1 |
| FUN_000504-T1 | 0.0068709 | 0.12452362 | TBU04623.1 |
| FUN_000505-T1 | 0.00648633 | 0.30159397 | TBU04624.1 |
| FUN_000506-T1 | 0 | NA | TBT97240.1 |
| FUN_000508-T1 | 0.00477496 | 0.16878264 | TBT97331.1 |
| FUN_000509-T1 | 0.00378574 | 0.12646779 | TBT98377.1 |
| FUN_000511-T1 | 0.00809674 | 1.16635042 | TBU05798.1 |
| FUN_000512-T1 | 0.00422132 | 0.59541679 | TBU05799.1 |
| FUN_000513-T1 | 0.00356764 | 0.27963264 | TBU02433.1 |
| FUN_000514-T1 | 0.01333561 | 1.44527082 | TBU02432.1 |
| FUN_000515-T1 | 0.00509493 | 0.18592729 | TBU07132.1 |
| FUN_000516-T1 | 0.03018063 | 0.7810382 | TBU07133.1 |
| FUN_000517-T1 | 0.01169466 | 0.81632107 | TBU07134.1 |
| FUN_000518-T1 | 0.00347412 | 0.32488151 | TBU07135.1 |
| FUN_000522-T1 | 0.00365903 | 0.28969039 | TBU03743.1 |
| FUN_000540-T1 | 0 | NA | TBU05672.1 |
| FUN_000541-T1 | 0.0020676 | 0.10209289 | TBU05673.1 |
| FUN_000542-T1 | 0.0039179 | 0.14531827 | TBU05674.1 |
| FUN_000544-T1 | 0.00117036 | 1.66297526 | TBU05675.1 |
| FUN_000545-T1 | 0.00225568 | 0.12832614 | TBU05676.1 |
| FUN_000546-T1 | 0.00069372 | 0.33833656 | TBU05677.1 |
| FUN_000547-T1 | 0.00136169 | 0.2602301 | TBU06146.1 |
| FUN_000549-T1 | 0.00255525 | 0.2380592 | TBU08018.1 |
| FUN_000551-T1 | 0.0034298 | 1.23777126 | TBU08017.1 |
| FUN_000554-T1 | 0.00274271 | 0.08925661 | TBU02903.1 |
| FUN_000555-T1 | 0.00417764 | 0.59124717 | TBU02902.1 |
| FUN_000556-T1 | 0.00468527 | 0.06937683 | TBU02530.1 |
| FUN_000557-T1 | 0.01290323 | 0.26923077 | TBU02529.1 |
| FUN_000558-T1 | 0.00168496 | 0 | TBU02528.1 |
| FUN_000559-T1 | 0.00406943 | 0.42150395 | TBU03822.1 |
| FUN_000560-T1 | 0.00239909 | 0.36070771 | TBU03821.1 |
| FUN_000561-T1 | 0.00068922 | 0.01673912 | TBU03820.1 |
| FUN_000562-T1 | 0 | NA | TBU03819.1 |
| FUN_000567-T1 | 0.00323054 | 0.13994396 | TBU03560.1 |
| FUN_000573-T1 | 0.00368664 | 0.05127704 | TBU08684.1 |
| FUN_000575-T1 | 0.0095696 | 0.72738472 | TBU08685.1 |
| FUN_000576-T1 | 0.00353033 | 3.26963845 | TBU00846.1 |
| FUN_000579-T1 | 0.00054215 | 0.10990888 | TBU05665.1 |
| FUN_000580-T1 | 0.00165771 | 0.01927058 | TBU05664.1 |
| FUN_000581-T1 | 0.00165939 | 0.68215961 | TBU05663.1 |
| FUN_000582-T1 | 0.00208018 | 0.10614408 | TBU05662.1 |
| FUN_000586-T1 | 0.00136252 | 1.7552734 | TBU01329.1 |
| FUN_000587-T1 | 0.0040681 | 0.14529253 | TBU04040.1 |
| FUN_000590-T1 | 0.00285475 | 0.11731494 | TBU08111.1 |
| FUN_000592-T1 | 0.00554889 | 0.15736254 | TBU08110.1 |
| FUN_000594-T1 | 0.04100711 | 0.65322587 | TBU02773.1 |
| FUN_000597-T1 | 0.00595602 | 0.1868872 | TBU00614.1 |
| FUN_000598-T1 | 0.02197768 | 0.4854599 | TBU01759.1 |
| FUN_000606-T1 | 0.01753014 | 0.39235999 | TBU07874.1 |
| FUN_000607-T1 | 0.00061444 | 0 | TBU07873.1 |
| FUN_000608-T1 | 0.00617512 | 0 | TBU07872.1 |
| FUN_000609-T1 | 0.00133129 | 0.05192089 | TBU07871.1 |
| FUN_000612-T1 | 0.03707636 | 0.96454898 | TBT98799.1 |
| FUN_000613-T1 | 0.0037881 | 0.1152392 | TBU08377.1 |
| FUN_000614-T1 | 0.00351025 | 0.19664563 | TBU08378.1 |
| FUN_000615-T1 | 0.00111456 | 0.01523665 | TBU08379.1 |
| FUN_000617-T1 | 0.00677648 | 0.33124228 | TBU08380.1 |
| FUN_000618-T1 | 0.00256741 | 0.0317461 | TBU00738.1 |
| FUN_000619-T1 | 0.01636356 | 1.56656614 | TBU06102.1 |
| FUN_000620-T1 | 0.00360234 | 0.02732843 | TBU06104.1 |
| FUN_000621-T1 | 0.00865921 | 0.26208097 | TBT99538.1 |
| FUN_000622-T1 | 0.00700902 | 0.26775263 | TBU00275.1 |
| FUN_000631-T1 | 0.00534341 | 0.47980265 | TBU00153.1 |
| FUN_000634-T1 | 0.00834664 | 0.45705604 | TBU02673.1 |
| FUN_000642-T1 | 0.00291847 | 0.07755011 | TBU01844.1 |
| FUN_000645-T1 | 0.0036525 | 0 | TBU04725.1 |
| FUN_000646-T1 | 0.00471698 | 0.15482381 | TBU04723.1 |
| FUN_000647-T1 | 0.01327189 | 0.7487723 | TBU05792.1 |
| FUN_000648-T1 | 0.00700531 | 0.28580553 | TBU00564.1 |
| FUN_000649-T1 | 0.00495993 | 0.05546676 | TBU06023.1 |
| FUN_000650-T1 | 0.00541617 | 0.09588326 | TBU06024.1 |
| FUN_000651-T1 | 0.00344832 | 0.12118302 | TBU03302.1 |
| FUN_000654-T1 | 0.00910636 | 0.46926665 | TBU07455.1 |
| FUN_000655-T1 | 0.00455685 | 0.05351261 | TBU07456.1 |
| FUN_000658-T1 | 0.00156253 | 0.21882521 | TBT99478.1 |
| FUN_000659-T1 | 0.00247209 | 0.27765749 | TBU01778.1 |
| FUN_000660-T1 | 0.00179893 | 0.16176621 | TBU01779.1 |
| FUN_000661-T1 | 0.00834027 | 1.07038303 | TBU02958.1 |
| FUN_000662-T1 | 0.00647348 | 0.65260471 | TBU02957.1 |
| FUN_000663-T1 | 0.00526909 | 0.26282192 | TBU01037.1 |
| FUN_000666-T1 | 0.00403087 | 0.22686602 | TBU00393.1 |
| FUN_000669-T1 | 0.00555027 | 0.09374275 | TBT97550.1 |
| FUN_000670-T1 | 0 | NA | TBU06329.1 |
| FUN_000672-T1 | 0.00061201 | Inf | TBU06328.1 |
| FUN_000674-T1 | 0.0023858 | 2.08960253 | TBU06327.1 |
| FUN_000675-T1 | 0.00366272 | 0.10461252 | TBU03447.1 |
| FUN_000676-T1 | 0.00325029 | 0.41198431 | TBU03446.1 |
| FUN_000677-T1 | 0.00325787 | 0.39104379 | TBU03445.1 |
| FUN_000681-T1 | 0.02617574 | 0.73416729 | TBU07399.1 |
| FUN_000682-T1 | 0.00393358 | 0.46386465 | TBU07398.1 |
| FUN_000690-T1 | 0.00226473 | 0.03289087 | TBU05892.1 |
| FUN_000691-T1 | 0.01881973 | 0.35608953 | TBT96815.1 |
| FUN_000692-T1 | 0.00270232 | 0.18541126 | TBU05893.1 |
| FUN_000694-T1 | 0.00321869 | 0.40894345 | TBU02511.1 |
| FUN_000700-T1 | 0.0129491 | 0.32778882 | TBU01469.1 |
| FUN_000705-T1 | 0.02580645 | 0.93693694 | TBU06649.1 |
| FUN_000714-T1 | 0.00144347 | 3.56460568 | TBU06799.1 |
| FUN_000715-T1 | 0.00081067 | 0.22698073 | TBT97968.1 |
| FUN_000716-T1 | 0.00194163 | 0 | TBU02497.1 |
| FUN_000718-T1 | 0.00316921 | 0.05177902 | TBT99232.1 |
| FUN_000719-T1 | 0.00762724 | 0.21408786 | TBT98356.1 |
| FUN_000722-T1 | 0.02213951 | 1.70890581 | TBT97569.1 |
| FUN_000723-T1 | 0.00262393 | 3.02962775 | TBU01270.1 |
| FUN_000725-T1 | 0.00521705 | 0.85151908 | TBU01030.1 |
| FUN_000727-T1 | 0.0097105 | 0.59877394 | TBU09512.1 |
| FUN_000728-T1 | 0.01145067 | 0.43508777 | TBU09511.1 |
| FUN_000729-T1 | 0.00665323 | 0.78057395 | TBU09510.1 |
| FUN_000730-T1 | 0.00348352 | 1.03726978 | TBU09509.1 |
| FUN_000731-T1 | 0.00477345 | 0.47123038 | TBU09508.1 |
| FUN_000732-T1 | 0.00493652 | 0.43240278 | TBU09507.1 |
| FUN_000734-T1 | 0.00298854 | 0.70543211 | TBU06263.1 |
| FUN_000735-T1 | 0.00206344 | 0 | TBU06264.1 |
| FUN_000736-T1 | 0.00412295 | 11.6798335 | TBU06265.1 |
| FUN_000737-T1 | 0.00779332 | 1.74912554 | TBT99885.1 |
| FUN_000738-T1 | 0.02839356 | 1.22655762 | TBU08012.1 |
| FUN_000739-T1 | 0.0152231 | 0.62196999 | TBU08011.1 |
| FUN_000740-T1 | 0.00217018 | 0.01918409 | TBU08010.1 |
| FUN_000741-T1 | 0.015553 | 0.2512403 | TBU04570.1 |
| FUN_000743-T1 | 0.00318008 | 0 | TBU04569.1 |
| FUN_000747-T1 | 0.0052264 | 0.07052332 | TBU08785.1 |
| FUN_000749-T1 | 0.00958392 | 0.89929139 | TBU07616.1 |
| FUN_000750-T1 | 0.01032126 | 2.08753935 | TBU07617.1 |
| FUN_000751-T1 | 0.00268358 | 0.00709698 | TBU07618.1 |
| FUN_000752-T1 | 0.01755091 | 4.35101167 | TBU03798.1 |
| FUN_000753-T1 | 0.01192058 | 0.22707354 | TBT99899.1 |
| FUN_000754-T1 | 0.00020098 | Inf | TBT96875.1 |
| FUN_000755-T1 | 0.00528359 | 0.30371217 | TBU00594.1 |
| FUN_000756-T1 | 0.0060242 | 0.26343024 | TBU01550.1 |
| FUN_000758-T1 | 0.01273087 | 0.8891834 | TBU03871.1 |
| FUN_000760-T1 | 0.00033428 | 0.10904628 | TBU02071.1 |
| FUN_000761-T1 | 0.00361203 | 0.15197358 | TBU02070.1 |
| FUN_000762-T1 | 0.00425763 | 0.41929164 | TBU00830.1 |
| FUN_000763-T1 | 0.01519793 | 0.64633705 | TBU07279.1 |
| FUN_000764-T1 | 0.00534864 | 0.37021698 | TBU07280.1 |
| FUN_000765-T1 | 0.01457403 | 0.63204734 | TBU07281.1 |
| FUN_000767-T1 | 0.03330346 | 0.79400861 | TBU07282.1 |
| FUN_000768-T1 | 0.00288965 | 0.1684625 | TBU00988.1 |
| FUN_000771-T1 | 0.00223559 | 0 | TBU08903.1 |
| FUN_000772-T1 | 0 | NA | TBU08904.1 |
| FUN_000773-T1 | 0.0033999 | 0.01623475 | TBU08905.1 |
| FUN_000775-T1 | 0.01378721 | 1.32635083 | TBU08906.1 |
| FUN_000776-T1 | 0.00356468 | 0.10739265 | TBU05915.1 |
| FUN_000786-T1 | 0.00052385 | Inf | TBU07601.1 |
| FUN_000787-T1 | 0.00129255 | 0.08551134 | TBU07602.1 |
| FUN_000789-T1 | 0.00151198 | 0.18670331 | TBU07603.1 |
| FUN_000790-T1 | 0.00793264 | 0.09268292 | TBU07604.1 |
| FUN_000791-T1 | 0.003059 | 0.09950978 | TBT98461.1 |
| FUN_000792-T1 | 0.00151595 | 1.71339286 | TBU02357.1 |
| FUN_000793-T1 | 0.00390671 | 1.55322209 | TBU03178.1 |
| FUN_000794-T1 | 0.01271207 | 0.24629945 | TBU03177.1 |
| FUN_000795-T1 | 0.04512931 | 1.76427355 | TBU03176.1 |
| FUN_000801-T1 | 0.00147714 | 0.24898959 | TBU08081.1 |
| FUN_000804-T1 | 0.00106833 | 0 | TBU04869.1 |
| FUN_000806-T1 | 0.00423352 | 0.2570213 | TBU04868.1 |
| FUN_000807-T1 | 0.00014831 | Inf | TBT98237.1 |
| FUN_000814-T1 | 0.00079265 | 0.0223308 | TBU07491.1 |
| FUN_000815-T1 | 0 | NA | TBU07490.1 |
| FUN_000816-T1 | 0.00350537 | 0.557757 | TBU07489.1 |
| FUN_000820-T1 | 0.00907283 | 0.21711249 | TBU04684.1 |
| FUN_000826-T1 | 0.00015145 | 0.59048408 | TBU04933.1 |
| FUN_000827-T1 | 0.00187863 | Inf | TBU04932.1 |
| FUN_000830-T1 | 0.00045116 | 1.15504474 | TBU01337.1 |
| FUN_000832-T1 | 0.00211311 | 0.15733401 | TBU08775.1 |
| FUN_000833-T1 | 0.001197 | 0.01950555 | TBU08774.1 |
| FUN_000834-T1 | 0.00319508 | Inf | TBU08773.1 |
| FUN_000837-T1 | 0.00214025 | 0.31230337 | TBU08772.1 |
| FUN_000839-T1 | 0.00377008 | 0.34113463 | TBT99128.1 |
| FUN_000842-T1 | 0.00629426 | 0.57852066 | TBU04672.1 |
| FUN_000844-T1 | 0.00797026 | 0.69462269 | TBU03575.1 |
| FUN_000848-T1 | 0.00248097 | 0.57888363 | TBU07008.1 |
| FUN_000850-T1 | 0.00468958 | 0.90527214 | TBU07010.1 |
| FUN_000851-T1 | 0.00228167 | 0 | TBU07011.1 |
| FUN_000853-T1 | 0 | NA | TBU03104.1 |
| FUN_000854-T1 | 0.00343509 | 1.14205109 | TBU03105.1 |
| FUN_000863-T1 | 0.00104013 | Inf | TBT98563.1 |
| FUN_000867-T1 | 0.00590498 | 0.51622006 | TBT98096.1 |
| FUN_000868-T1 | 0.00409626 | 2.16784203 | TBU07720.1 |
| FUN_000869-T1 | 0.00434452 | 4.81985294 | TBT98961.1 |
| FUN_000875-T1 | 0.0091684 | 1.26085089 | TBU01484.1 |
| FUN_000878-T1 | 0 | NA | TBU07721.1 |
| FUN_000884-T1 | 0.00298173 | 0.14073784 | TBU03008.1 |
| FUN_000885-T1 | 0.00483159 | 0.27315126 | TBU07479.1 |
| FUN_000886-T1 | 0.00641118 | 0.06748913 | TBU07480.1 |
| FUN_000887-T1 | 0.00326517 | 0.07001126 | TBU07481.1 |
| FUN_000888-T1 | 0.00339277 | 0.00449533 | TBU07482.1 |
| FUN_000889-T1 | 0.00147586 | 0.02881698 | TBU07483.1 |
| FUN_000890-T1 | 0.00531131 | 0.36065198 | TBT99271.1 |
| FUN_000900-T1 | 0.0026048 | 0.09721668 | TBU06821.1 |
| FUN_000901-T1 | 0.00159075 | 0.11303572 | TBU06822.1 |
| FUN_000902-T1 | 0.00481921 | 0.49361822 | TBU06823.1 |
| FUN_000903-T1 | 0.00257844 | 0.433368 | TBU05608.1 |
| FUN_000904-T1 | 0.00661643 | 0.19962691 | TBU03493.1 |
| FUN_000906-T1 | 0.0029269 | 0.18559146 | TBU02299.1 |
| FUN_000908-T1 | 0.00373147 | 14.9558292 | TBU04644.1 |
| FUN_000909-T1 | 0.00258065 | 0.04749726 | TBU08161.1 |
| FUN_000910-T1 | 0.00795905 | 0.10664409 | TBU08162.1 |
| FUN_000911-T1 | 0.00390074 | 0.37254797 | TBU08163.1 |
| FUN_000912-T1 | 0.00276239 | 0.10909525 | TBU08164.1 |
| FUN_000923-T1 | 0.00116877 | 0.36468423 | TBU05015.1 |
| FUN_000924-T1 | 0.00371527 | 0.71597676 | TBU05016.1 |
| FUN_000925-T1 | 0.03113278 | 0.93448451 | TBU05017.1 |
| FUN_000926-T1 | 0.00229724 | 0.05244401 | TBU05018.1 |
| FUN_000931-T1 | 0.00195665 | 0.12084527 | TBT98293.1 |
| FUN_000932-T1 | 0.00432439 | 0.28934314 | TBU03037.1 |
| FUN_000939-T1 | 0.0038142 | 0.27154761 | TBU06503.1 |
| FUN_000940-T1 | 0.00232156 | 0.07875197 | TBU06505.1 |
| FUN_000941-T1 | 0.00136542 | 0.34346173 | TBU06506.1 |
| FUN_000943-T1 | 0.00063458 | Inf | TBU03793.1 |
| FUN_000944-T1 | 0.00178846 | 0.54063577 | TBU02111.1 |
| FUN_000945-T1 | 0.00124434 | 0 | TBU01497.1 |
| FUN_000946-T1 | 0.00398862 | 0.31266603 | TBU01498.1 |
| FUN_000947-T1 | 0 | NA | TBU01499.1 |
| FUN_000949-T1 | 0.00860458 | 0.66947036 | TBU00555.1 |
| FUN_000950-T1 | 0.00467625 | 0.22716924 | TBU01461.1 |
| FUN_000953-T1 | 0.02106044 | 0.41300198 | TBU05523.1 |
| FUN_000954-T1 | 0.02267228 | 0.99029557 | TBU05524.1 |
| FUN_000957-T1 | 0.0010552 | Inf | TBU06192.1 |
| FUN_000958-T1 | 0.00231439 | 0.06062883 | TBU06193.1 |
| FUN_000959-T1 | 0.00277893 | 1.36143978 | TBU06194.1 |
| FUN_000964-T1 | 0.00375414 | 0.47084739 | TBU05023.1 |
| FUN_000965-T1 | 0.00254327 | 0.45456417 | TBU06323.1 |
| FUN_000973-T1 | 0.00121009 | Inf | TBU06479.1 |
| FUN_000975-T1 | 0.0069263 | 0.67781499 | TBU07932.1 |
| FUN_000976-T1 | 0.01445443 | 0.63007181 | TBU06293.1 |
| FUN_000977-T1 | 0.0032694 | 0.34077047 | TBU07933.1 |
| FUN_000978-T1 | 0.00068568 | 0 | TBU07934.1 |
| FUN_000979-T1 | 0.00170251 | Inf | TBU00036.1 |
| FUN_000980-T1 | 0.00038679 | 0.18220223 | TBU00037.1 |
| FUN_000981-T1 | 0.00060864 | 0.12598425 | TBU01735.1 |
| FUN_000984-T1 | 0.00024438 | 0 | TBU01093.1 |
| FUN_000985-T1 | 0.00036692 | 0 | TBT98856.1 |
| FUN_000987-T1 | 0.00190686 | 0.1818239 | TBU07586.1 |
| FUN_000989-T1 | 0.00331751 | 0.68899285 | TBU07583.1 |
| FUN_000990-T1 | 0.00011085 | 0 | TBU01387.1 |
| FUN_000991-T1 | 0.00345781 | 0.2448205 | TBU01388.1 |
| FUN_000992-T1 | 0.00290863 | 0.35406707 | TBU00626.1 |
| FUN_000995-T1 | 0.0035516 | 0.15979853 | TBU00816.1 |
| FUN_000996-T1 | 0.00105536 | 2.31184319 | TBU00815.1 |
| FUN_000998-T1 | 0.00011881 | 0 | TBT99063.1 |
| FUN_000999-T1 | 0 | NA | TBT99062.1 |
| FUN_001000-T1 | 0.00128775 | 0 | TBU01120.1 |
| FUN_001003-T1 | 0.01246738 | 0.19382367 | TBU07091.1 |
| FUN_001015-T1 | 0.00191787 | 1.19229124 | TBT99417.1 |
| FUN_001016-T1 | 0.00136542 | Inf | TBU08760.1 |
| FUN_001017-T1 | 0.00183428 | 0.04926246 | TBU08761.1 |
| FUN_001018-T1 | 0.00183855 | 0.10173651 | TBU08762.1 |
| FUN_001025-T1 | 0.00365064 | 0.40652168 | TBU08291.1 |
| FUN_001026-T1 | 0.00581773 | 0.33871195 | TBU06234.1 |
| FUN_001027-T1 | 0.00285419 | 0.18168376 | TBU06233.1 |
| FUN_001028-T1 | 0.00032916 | 1.08726805 | TBU06232.1 |
| FUN_001029-T1 | 0.017981 | 1.54215932 | TBU06231.1 |
| FUN_001030-T1 | 0.00151562 | 0.01266217 | TBU05259.1 |
| FUN_001031-T1 | 0.00229469 | 0.17429727 | TBU05260.1 |
| FUN_001032-T1 | 0.00316884 | 0.11015307 | TBU05261.1 |
| FUN_001033-T1 | 0.0040342 | 0.13280204 | TBU05262.1 |
| FUN_001035-T1 | 0.02903226 | 0.41522491 | TBU05205.1 |
| FUN_001042-T1 | 0.00095157 | Inf | TBU09239.1 |
| FUN_001045-T1 | 0.00123392 | 0 | TBU06055.1 |
| FUN_001047-T1 | 0.00228144 | 0.42253343 | TBU06054.1 |
| FUN_001048-T1 | 0.00272545 | 0.06358838 | TBU06741.1 |
| FUN_001049-T1 | 0.00494688 | 0.11931736 | TBU02361.1 |
| FUN_001051-T1 | 0.04215054 | 2.21480485 | TBU02381.1 |
| FUN_001059-T1 | 0.02470575 | 5.1561422 | TBU09522.1 |
| FUN_001060-T1 | 0.00880058 | 0.33563924 | TBU09523.1 |
| FUN_001062-T1 | 0.00381718 | 0.04170733 | TBU07085.1 |
| FUN_001063-T1 | 0.00341256 | 0.05485407 | TBU07084.1 |
| FUN_001064-T1 | 0.00470202 | 0.24765598 | TBU07083.1 |
| FUN_001070-T1 | 0.01172807 | 0.49976466 | TBU00876.1 |
| FUN_001071-T1 | 0.01870599 | 1.44764756 | TBU00877.1 |
| FUN_001073-T1 | 0.00710994 | 1.0545214 | TBU03134.1 |
| FUN_001076-T1 | 0.00368057 | 0.37469611 | TBT97725.1 |
| FUN_001078-T1 | 0.00947992 | 0.73346082 | TBU05703.1 |
| FUN_001079-T1 | 0.00651637 | 0.28561256 | TBU05704.1 |
| FUN_001080-T1 | 0.00846334 | 0.52392376 | TBU00581.1 |
| FUN_001081-T1 | 0.0020521 | 0.11371186 | TBU06579.1 |
| FUN_001082-T1 | 0.00084688 | 0.16302036 | TBU06580.1 |
| FUN_001083-T1 | 0.00196759 | 0.07118671 | TBU06581.1 |
| FUN_001084-T1 | 0.00231968 | Inf | TBU06582.1 |
| FUN_001085-T1 | 0.00048359 | 0.31340638 | TBU06583.1 |
| FUN_001086-T1 | 0.00258783 | 0.91515219 | TBT98167.1 |
| FUN_001087-T1 | 0.00175757 | 0.4741503 | TBU09355.1 |
| FUN_001090-T1 | 0.00283154 | 0.2613354 | TBU09357.1 |
| FUN_001091-T1 | 0.00191335 | 2.92603478 | TBU09358.1 |
| FUN_001093-T1 | 0.01091133 | 0.15822009 | TBU08000.1 |
| FUN_001094-T1 | 0.0278994 | 0.55169339 | TBU08001.1 |
| FUN_001095-T1 | 0.0040553 | 0.08037431 | TBU08002.1 |
| FUN_001098-T1 | 0.0041165 | 0.11576982 | TBU06178.1 |
| FUN_001100-T1 | 0.00602391 | 0.07331787 | TBU06177.1 |
| FUN_001101-T1 | 0.01168017 | 0.43259851 | TBU06176.1 |
| FUN_001102-T1 | 0.00330737 | 0.07783931 | TBU06906.1 |
| FUN_001103-T1 | 0.00741662 | 0.5086735 | TBU06907.1 |
| FUN_001108-T1 | 0.00242542 | 3.59638248 | TBU04584.1 |
| FUN_001111-T1 | 0.01226985 | 0.23317492 | TBU02516.1 |
| FUN_001123-T1 | 0.0036443 | 0.15592719 | TBU07036.1 |
| FUN_001125-T1 | 0.00627858 | 1.46975263 | TBU06830.1 |
| FUN_001126-T1 | 0.02374606 | 0.91449089 | TBU06828.1 |
| FUN_001129-T1 | 0.01283187 | 1.03450294 | TBU02014.1 |
| FUN_001132-T1 | 0.01762842 | 0.24177109 | TBT97393.1 |
| FUN_001134-T1 | 0.01424731 | 2.52293409 | TBU04877.1 |
| FUN_001137-T1 | 0.00365997 | 0.9183911 | TBU06278.1 |
| FUN_001138-T1 | 0.00302216 | 0.49086534 | TBU06277.1 |
| FUN_001139-T1 | 0.00069613 | 0 | TBU06276.1 |
| FUN_001140-T1 | 0.00255529 | 0.50326604 | TBU06374.1 |
| FUN_001141-T1 | 0.00110064 | 5.30282764 | TBU06375.1 |
| FUN_001142-T1 | 0.00596689 | 0.27392753 | TBU06376.1 |
| FUN_001143-T1 | 0.01031155 | 0.27535331 | TBU06377.1 |
| FUN_001144-T1 | 0.00253868 | 0 | TBT97810.1 |
| FUN_001145-T1 | 0.00663321 | 0.5425031 | TBT97130.1 |
| FUN_001147-T1 | 0.02076381 | 0.4849791 | TBU00026.1 |
| FUN_001148-T1 | 0.00428144 | 0.60481816 | TBU07019.1 |
| FUN_001149-T1 | 0.00484227 | 0.6225278 | TBT98053.1 |
| FUN_001157-T1 | 0.00828853 | 0.10139745 | TBU07016.1 |
| FUN_001159-T1 | 0.04382772 | 1.32540211 | TBU06000.1 |
| FUN_001162-T1 | 0.02551085 | 1.5697558 | TBT97193.1 |
| FUN_001166-T1 | 0.00143369 | 0.18534434 | TBU08750.1 |
| FUN_001167-T1 | 0.00351156 | 4.08444074 | TBU08749.1 |
| FUN_001168-T1 | 0.00304468 | 0.14519519 | TBU08748.1 |
| FUN_001169-T1 | 0.01313008 | 1.6675948 | TBU08747.1 |
| FUN_001170-T1 | 0.0004368 | 0.83200689 | TBU08746.1 |
| FUN_001171-T1 | 0.00220831 | 0.05541721 | TBU07900.1 |
| FUN_001172-T1 | 0.00395851 | 4.07404291 | TBU07899.1 |
| FUN_001175-T1 | 0.00315014 | 1.02061002 | TBT98592.1 |
| FUN_001179-T1 | 0.00311421 | 0.18126529 | TBU07976.1 |
| FUN_001180-T1 | 7.17E-05 | 0 | TBU07975.1 |
| FUN_001181-T1 | 0.00106531 | 0 | TBU07974.1 |
| FUN_001182-T1 | 0.00097349 | 0 | TBU07973.1 |
| FUN_001183-T1 | 0.00196232 | 0.04927454 | TBU07972.1 |
| FUN_001184-T1 | 0.00296668 | 0.19679519 | TBU07971.1 |
| FUN_001185-T1 | 0.00276729 | 0.04538409 | TBU08314.1 |
| FUN_001186-T1 | 0.01254012 | 0.91194086 | TBU08313.1 |
| FUN_001187-T1 | 0.01513341 | 1.14986456 | TBU08312.1 |
| FUN_001188-T1 | 0.00590641 | 0.70074818 | TBU08311.1 |
| FUN_001189-T1 | 0.00303605 | 0.17047029 | TBU07438.1 |
| FUN_001191-T1 | 0.00182334 | 0.29719757 | TBU07437.1 |
| FUN_001192-T1 | 0.00215666 | 0.00539082 | TBU07436.1 |
| FUN_001195-T1 | 0.00602566 | 0.07601086 | TBU03083.1 |
| FUN_001196-T1 | 0.00389921 | 0.33665628 | TBU03084.1 |
| FUN_001198-T1 | 0.00876003 | 0.37210023 | TBU08156.1 |
| FUN_001206-T1 | 0.00585614 | 0.22615153 | TBU07116.1 |
| FUN_001208-T1 | 0.00637502 | 0.29049965 | TBU03145.1 |
| FUN_001209-T1 | 0.00532492 | 0.25127295 | TBU03144.1 |
| FUN_001212-T1 | 0.00010542 | Inf | TBU01344.1 |
| FUN_001213-T1 | 0.00654411 | 0.11321712 | TBT99578.1 |
| FUN_001214-T1 | 0.00331548 | 0.28991881 | TBU01992.1 |
| FUN_001215-T1 | 0.00387772 | 0.75175843 | TBU01991.1 |
| FUN_001217-T1 | 0.01733464 | 0.66953377 | TBU01764.1 |
| FUN_001218-T1 | 0.00476831 | 0.52371063 | TBU06511.1 |
| FUN_001221-T1 | 0.00385509 | 1.4835506 | TBT99239.1 |
| FUN_001223-T1 | 0.00205519 | 0.01234118 | TBU00110.1 |
| FUN_001225-T1 | 0.00837578 | 0.97720274 | TBU01486.1 |
| FUN_001227-T1 | 0.00480584 | 0.4040517 | TBU09020.1 |
| FUN_001228-T1 | 0.00240176 | 0.44660908 | TBU03924.1 |
| FUN_001230-T1 | 0.00046214 | 0.45616205 | TBT99088.1 |
| FUN_001234-T1 | 0.0034188 | 0.53994548 | TBT99971.1 |
| FUN_001235-T1 | 0.00564204 | 0.70269806 | TBT97816.1 |
| FUN_001236-T1 | 0.00930352 | 0.20650723 | TBT99135.1 |
| FUN_001237-T1 | 0.01119942 | 0.78614274 | TBT99136.1 |
| FUN_001239-T1 | 0.00296534 | 0.25909214 | TBU06221.1 |
| FUN_001240-T1 | 0.00712238 | 0.65583783 | TBU00885.1 |
| FUN_001242-T1 | 0.02147612 | 0.42095463 | TBU06406.1 |
| FUN_001243-T1 | 0.00151987 | 0.09883199 | TBU06407.1 |
| FUN_001244-T1 | 0.00094447 | 0.32223109 | TBU02095.1 |
| FUN_001246-T1 | 0 | NA | TBU02094.1 |
| FUN_001251-T1 | 0.00287974 | 0.02826061 | TBU04361.1 |
| FUN_001252-T1 | 0.00629447 | 0.12799268 | TBU04362.1 |
| FUN_001253-T1 | 0.00273273 | 0.05030215 | TBU02763.1 |
| FUN_001254-T1 | 0.00340392 | 0.3390639 | TBU02762.1 |
| FUN_001255-T1 | 0.00695455 | 0.57726391 | TBU07324.1 |
| FUN_001258-T1 | 0.00351869 | 0.03954162 | TBU07325.1 |
| FUN_001259-T1 | 0.00267936 | 0 | TBU07326.1 |
| FUN_001261-T1 | 0.00338972 | 0.41756425 | TBU05421.1 |
| FUN_001264-T1 | 0.01162159 | 1.43022215 | TBU02166.1 |
| FUN_001266-T1 | 0.0006372 | 0.80121779 | TBT97971.1 |
| FUN_001267-T1 | 0.0030459 | 0.32284141 | TBU09535.1 |
| FUN_001268-T1 | 0.00213928 | 0.08187321 | TBU09534.1 |
| FUN_001270-T1 | 0.00314309 | 0.82355859 | TBU09533.1 |
| FUN_001271-T1 | 0.00427968 | 0.41993169 | TBU09532.1 |
| FUN_001274-T1 | 0.0061808 | 0 | TBU09531.1 |
| FUN_001279-T1 | 0.01575137 | 0.66666199 | TBU05432.1 |
| FUN_001281-T1 | 0.01535509 | 0.39640044 | TBU05433.1 |
| FUN_001289-T1 | 0.00108846 | 0.43871145 | TBU03982.1 |
| FUN_001291-T1 | 0.00397022 | 0.40823249 | TBU03201.1 |
| FUN_001296-T1 | 0.00524017 | 0.07649508 | TBU07288.1 |
| FUN_001301-T1 | 0.00423497 | 0.11621056 | TBT99953.1 |
| FUN_001313-T1 | 0.00584772 | 0.21734312 | TBU05130.1 |
| FUN_001314-T1 | 0.00335104 | 0.05331541 | TBU05129.1 |
| FUN_001315-T1 | 0.00238949 | 0 | TBU05127.1 |
| FUN_001316-T1 | 0.00815972 | 0.38933748 | TBU09606.1 |
| FUN_001317-T1 | 0.00866043 | 0.32919697 | TBU09608.1 |
| FUN_001318-T1 | 0.00662092 | 0.15530236 | TBU09610.1 |
| FUN_001321-T1 | 0.00141294 | 0.24216793 | TBU09611.1 |
| FUN_001327-T1 | 0.00446677 | 0.44683753 | TBU02182.1 |
| FUN_001328-T1 | 0.00571857 | 1.92364448 | TBU06097.1 |
| FUN_001329-T1 | 0.00354905 | 0.08355029 | TBU06098.1 |
| FUN_001331-T1 | 0.00327106 | 0.3957687 | TBU02085.1 |
| FUN_001332-T1 | 0.00466155 | 0.16886384 | TBU03670.1 |
| FUN_001335-T1 | 0.00310695 | 0.16241887 | TBU08331.1 |
| FUN_001339-T1 | 0.00304116 | 0.98791889 | TBT99792.1 |
| FUN_001340-T1 | 0.00499582 | 0.45509446 | TBU01165.1 |
| FUN_001342-T1 | 0.00305041 | 0.17955127 | TBU01546.1 |
| FUN_001344-T1 | 0.00181452 | 0.02195815 | TBT99379.1 |
| FUN_001348-T1 | 0.00060079 | 0 | TBU04744.1 |
| FUN_001358-T1 | 0.00463256 | 0.08316903 | TBT98387.1 |
| FUN_001363-T1 | 0.00959471 | 0.74720738 | TBU04448.1 |
| FUN_001366-T1 | 0.02300254 | 1.88679478 | TBU07198.1 |
| FUN_001367-T1 | 0.00086988 | 0.20348964 | TBU07197.1 |
| FUN_001368-T1 | 0.00627093 | 0.21920379 | TBU07196.1 |
| FUN_001370-T1 | 0.00204813 | 0.16755459 | TBU03993.1 |
| FUN_001371-T1 | 0.00226001 | 0 | TBU03992.1 |
| FUN_001372-T1 | 0.00215864 | 1.10459788 | TBU03991.1 |
| FUN_001373-T1 | 0.0051451 | 0.50078859 | TBU03990.1 |
| FUN_001374-T1 | 0.00202694 | 0.04303483 | TBU03989.1 |
| FUN_001375-T1 | 0.0030945 | 0.20184249 | TBU03988.1 |
| FUN_001376-T1 | 0.00114695 | 0 | TBU00538.1 |
| FUN_001377-T1 | 0.0011176 | 0 | TBU04432.1 |
| FUN_001378-T1 | 0.00452296 | 0.09127574 | TBU04433.1 |
| FUN_001379-T1 | 0.00421084 | 3.32857952 | TBU04434.1 |
| FUN_001380-T1 | 0.00189247 | 0 | TBU04435.1 |
| FUN_001381-T1 | 0.00105116 | 0.78845541 | TBT98747.1 |
| FUN_001382-T1 | 0.00628372 | 0.3703365 | TBU05725.1 |
| FUN_001383-T1 | 0.00557185 | 0.29154069 | TBU05724.1 |
| FUN_001389-T1 | 0.02197707 | 0.82691296 | TBU08221.1 |
| FUN_001394-T1 | 0.00895788 | 0.37269891 | TBU07536.1 |
| FUN_001395-T1 | 0.00293602 | 0.11809558 | TBU07535.1 |
| FUN_001396-T1 | 0.00189188 | 1.23416742 | TBU07534.1 |
| FUN_001400-T1 | 0.00572566 | 0.32039854 | TBU08617.1 |
| FUN_001402-T1 | 0.00730643 | 0.19555166 | TBU01568.1 |
| FUN_001403-T1 | 0.00237837 | 0.0188632 | TBU01567.1 |
| FUN_001406-T1 | 0.0062963 | 4.21246305 | TBU07314.1 |
| FUN_001408-T1 | 0.01009729 | 1.00306367 | TBU04441.1 |
| FUN_001409-T1 | 0.00281837 | 0.15095486 | TBU04442.1 |
| FUN_001411-T1 | 0.00335053 | 4.29186712 | TBT97646.1 |
| FUN_001418-T1 | 0.00033602 | 0 | TBU06930.1 |
| FUN_001421-T1 | 0.00259643 | 0.56299025 | TBU04550.1 |
| FUN_001423-T1 | 0.0039837 | 0.32037657 | TBU04549.1 |
| FUN_001424-T1 | 0.01037647 | 0.29773254 | TBU03465.1 |
| FUN_001425-T1 | 0.0058457 | 0.47009655 | TBU04370.1 |
| FUN_001426-T1 | 0.00959042 | 0.22799358 | TBU04371.1 |
| FUN_001427-T1 | 0.01040182 | 0.37231895 | TBU01059.1 |
| FUN_001429-T1 | 0.02946019 | 1.1192495 | TBU04973.1 |
| FUN_001433-T1 | 0.00739398 | 0.00788339 | TBU04100.1 |
| FUN_001436-T1 | 0.0053887 | 1.35016087 | TBU03655.1 |
| FUN_001438-T1 | 0.0013108 | 0.34300607 | TBU03528.1 |
| FUN_001439-T1 | 0.01682295 | 0.41677343 | TBU02306.1 |
| FUN_001442-T1 | 0.00422638 | 0.11174423 | TBU00744.1 |
| FUN_001443-T1 | 0.00203887 | 0.11938768 | TBU00745.1 |
| FUN_001446-T1 | 0.00952255 | 2.85278959 | TBU09314.1 |
| FUN_001454-T1 | 0.00621705 | 0.57366771 | TBT96707.1 |
| FUN_001456-T1 | 0.00720096 | 0.91464437 | TBU06256.1 |
| FUN_001457-T1 | 0.0029645 | 0.78224695 | TBU08535.1 |
| FUN_001458-T1 | 0.00373869 | 0.82000762 | TBU08536.1 |
| FUN_001460-T1 | 0.00595254 | 0.27998123 | TBU06703.1 |
| FUN_001461-T1 | 0.00221935 | 0.66047055 | TBU06702.1 |
| FUN_001462-T1 | 0.00477897 | 0.28946586 | TBU01174.1 |
| FUN_001464-T1 | 0.00297588 | 1.74773861 | TBU04535.1 |
| FUN_001469-T1 | 0.009823 | 0.69040361 | TBU05360.1 |
| FUN_001470-T1 | 0.00178106 | 0.07287056 | TBT98469.1 |
| FUN_001472-T1 | 0.00397654 | 0.2125302 | TBU00765.1 |
| FUN_001473-T1 | 0.00347803 | 0 | TBU00645.1 |
| FUN_001474-T1 | 0.00417277 | 0.09844486 | TBU08452.1 |
| FUN_001475-T1 | 0.04096685 | 0.71369924 | TBU08451.1 |
| FUN_001476-T1 | 0.00277803 | 7.22291843 | TBU08450.1 |
| FUN_001477-T1 | 0.0081402 | 1.40035082 | TBU08449.1 |
| FUN_001478-T1 | 0.00209223 | 1.09209132 | TBU08448.1 |
| FUN_001480-T1 | 0.00221876 | 0.27395941 | TBU06314.1 |
| FUN_001483-T1 | 0.00203175 | 0 | TBU00454.1 |
| FUN_001484-T1 | 0.00630917 | 0.28498857 | TBU00792.1 |
| FUN_001485-T1 | 0.00763963 | 0.55757638 | TBT98446.1 |
| FUN_001486-T1 | 0.00275652 | 0.18025593 | TBU07693.1 |
| FUN_001488-T1 | 0.0072282 | 0.40381461 | TBU07691.1 |
| FUN_001491-T1 | 0.00215054 | 0 | TBU09209.1 |
| FUN_001492-T1 | 0.01446967 | 0.82857769 | TBU09208.1 |
| FUN_001494-T1 | 0.00156063 | 1.53832065 | TBU09207.1 |
| FUN_001496-T1 | 0.00414646 | 0.4194643 | TBU09205.1 |
| FUN_001499-T1 | 0.01383513 | 0.14373661 | TBU08356.1 |
| FUN_001500-T1 | 0.00248091 | 0.19582931 | TBU08355.1 |
| FUN_001502-T1 | 0.01269916 | 0.51283317 | TBU08354.1 |
| FUN_001503-T1 | 0.00238047 | 0.03286734 | TBU08353.1 |
| FUN_001504-T1 | 0.00286738 | 0 | TBU08352.1 |
| FUN_001505-T1 | 0.00955795 | 0.51406439 | TBU00637.1 |
| FUN_001506-T1 | 0.00307719 | 0.72564717 | TBU00549.1 |
| FUN_001514-T1 | 0.00048327 | 0.27149368 | TBU07520.1 |
| FUN_001516-T1 | 0.00426643 | 0.42447014 | TBU07666.1 |
| FUN_001517-T1 | 0.00381441 | 0.46919155 | TBU07667.1 |
| FUN_001523-T1 | 0.00069056 | 1.26400057 | TBU05974.1 |
| FUN_001525-T1 | 0.00340804 | 0.69449885 | TBU01012.1 |
| FUN_001526-T1 | 0.00954809 | 0.24370259 | TBU02745.1 |
| FUN_001527-T1 | 0.00028674 | Inf | TBU04478.1 |
| FUN_001528-T1 | 0.00346239 | 0.25864419 | TBU04479.1 |
| FUN_001529-T1 | 0.00259857 | 0.47643545 | TBU04480.1 |
| FUN_001537-T1 | 0.00136852 | 0 | TBU09112.1 |
| FUN_001538-T1 | 0.00261399 | 0.17035636 | TBU01708.1 |
| FUN_001543-T1 | 0.00550698 | 0.14834607 | TBU06895.1 |
| FUN_001544-T1 | 0.00646696 | 0.31122942 | TBU06897.1 |
| FUN_001545-T1 | 0.00242772 | 0.14810234 | TBU06898.1 |
| FUN_001550-T1 | 0.00115138 | 0.43920614 | TBU01922.1 |
| FUN_001553-T1 | 0.0188192 | 1.75645161 | TBT97823.1 |
| FUN_001555-T1 | 0.05010272 | 0.59365103 | TBT97050.1 |
| FUN_001557-T1 | 0.00284349 | Inf | TBT96762.1 |
| FUN_001563-T1 | 0.00712411 | 0.52833996 | TBT99182.1 |
| FUN_001567-T1 | 0.0141481 | 0.52235579 | TBU07768.1 |
| FUN_001568-T1 | 0.00549773 | 0.725918 | TBT97658.1 |
| FUN_001569-T1 | 0.00221379 | 0 | TBU00631.1 |
| FUN_001571-T1 | 0.00604111 | 0.21294241 | TBT98577.1 |
| FUN_001572-T1 | 0.00268239 | 0.51905381 | TBT96737.1 |
| FUN_001580-T1 | 0.00264861 | 0.15715065 | TBU00067.1 |
| FUN_001582-T1 | 0.00868244 | 0.32719199 | TBU05684.1 |
| FUN_001583-T1 | 0.00680675 | 1.552812 | TBU05683.1 |
| FUN_001584-T1 | 0.00174312 | Inf | TBU05682.1 |
| FUN_001592-T1 | 0.01068752 | 0.15472534 | TBU09633.1 |
| FUN_001593-T1 | 0.00335856 | 0.92839415 | TBU09634.1 |
| FUN_001595-T1 | 0.01064029 | 0.45400538 | TBU09635.1 |
| FUN_001596-T1 | 0.00484254 | 0.05491804 | TBU09636.1 |
| FUN_001597-T1 | 0.00127334 | 0 | TBU03832.1 |
| FUN_001600-T1 | 0.00378993 | 0.4408634 | TBT98069.1 |
| FUN_001602-T1 | 0.00321734 | 0.7974674 | TBU07261.1 |
| FUN_001607-T1 | 0.00181499 | Inf | TBU00698.1 |
| FUN_001617-T1 | 0.002716 | 0.03426834 | TBU04205.1 |
| FUN_001618-T1 | 0.00410036 | 0.17014027 | TBU04206.1 |
| FUN_001619-T1 | 0.00334569 | 0.03828811 | TBU09283.1 |
| FUN_001620-T1 | 0.00106718 | 0 | TBU03228.1 |
| FUN_001621-T1 | 0.00706797 | 0.01993143 | TBU03229.1 |
| FUN_001622-T1 | 0.00039978 | Inf | TBU00262.1 |
| FUN_001624-T1 | 0.00523123 | 0 | TBU05074.1 |
| FUN_001626-T1 | 0.00674049 | 0.04837509 | TBU05114.1 |
| FUN_001627-T1 | 0.0035325 | 0.1693206 | TBU05113.1 |
| FUN_001628-T1 | 0.00385343 | 0.16801197 | TBU05112.1 |
| FUN_001629-T1 | 0.00111163 | 0.18861947 | TBU00165.1 |
| FUN_001630-T1 | 0.01564702 | 0.29687036 | TBT99433.1 |
| FUN_001632-T1 | 0.00201576 | 0 | TBU09282.1 |
| FUN_001635-T1 | 0.020327 | 0.69942887 | TBU09285.1 |
| FUN_001652-T1 | 0.00144574 | 0.66821346 | TBU07752.1 |
| FUN_001662-T1 | 0.00688383 | 0.3473996 | TBU05581.1 |
| FUN_001670-T1 | 0.00469968 | 0.5214612 | TBU01863.1 |
| FUN_001690-T1 | 0.00191159 | 0.10702703 | TBU01864.1 |
| FUN_001694-T1 | 0.02172952 | 1.87077855 | TBU05928.1 |
| FUN_001696-T1 | 0.00377797 | 0.52965725 | TBU05582.1 |
| FUN_001697-T1 | 0.00124423 | 0.17917715 | TBU05583.1 |
| FUN_001709-T1 | 0.02221128 | 0.76763811 | TBU03900.1 |
| FUN_001725-T1 | 0.00498831 | 1.23197191 | TBU08711.1 |
| FUN_001726-T1 | 0.00323967 | 0.10823879 | TBU00759.1 |
| FUN_001727-T1 | 0.00307663 | 0.13919277 | TBU01695.1 |
| FUN_001728-T1 | 0.00381092 | 0 | TBU01696.1 |
| FUN_001729-T1 | 0.00357617 | 0.16520991 | TBT99816.1 |
| FUN_001730-T1 | 0.04696454 | 0.64165218 | TBT99817.1 |
| FUN_001731-T1 | 0.00246294 | 0.1084539 | TBT98078.1 |
| FUN_001732-T1 | 0.00361271 | 0.07836822 | TBU06616.1 |
| FUN_001733-T1 | 0.01999188 | 0.31591416 | TBU06617.1 |
| FUN_001734-T1 | 0.0094148 | 0.55625836 | TBU02834.1 |
| FUN_001735-T1 | 0.00096024 | 1.01532417 | TBU02835.1 |
| FUN_001738-T1 | 0.00817887 | 3.37535737 | TBU02976.1 |
| FUN_001739-T1 | 0.00339465 | 0.07296487 | TBU02975.1 |
| FUN_001740-T1 | 0.01492101 | 4.48642715 | TBU02974.1 |
| FUN_001741-T1 | 0.00343478 | 0.05954327 | TBU02973.1 |
| FUN_001742-T1 | 0.00294488 | 0.33915358 | TBU05716.1 |
| FUN_001743-T1 | 0.0021109 | 0.39987516 | TBU05713.1 |
| FUN_001744-T1 | 0 | NA | TBT98189.1 |
| FUN_001745-T1 | 0.00068271 | 0 | TBT98172.1 |
| FUN_001748-T1 | 0.04050371 | 1.07813617 | TBT99257.1 |
| FUN_001749-T1 | 0.00545659 | 0.09813892 | TBU06064.1 |
| FUN_001750-T1 | 0.00233326 | 0.23722548 | TBU06065.1 |
| FUN_001757-T1 | 0.01223851 | 1.71582184 | TBU04651.1 |
| FUN_001759-T1 | 0.00189669 | 0.57877631 | TBU04652.1 |
| FUN_001763-T1 | 0.01012545 | 0.30182201 | TBT97007.1 |
| FUN_001769-T1 | 0.00013698 | 0 | TBU03622.1 |
| FUN_001770-T1 | 0.00419079 | 0.21243523 | TBT97253.1 |
| FUN_001772-T1 | 0.00187374 | 1.6936161 | TBU04797.1 |
| FUN_001773-T1 | 0.00996332 | 0.49220348 | TBU01933.1 |
| FUN_001774-T1 | 0.00730265 | 2.710839 | TBU06111.1 |
| FUN_001776-T1 | 0.00744627 | 0.32685174 | TBT98481.1 |
| FUN_001781-T1 | 0.00099256 | 0 | TBU03826.1 |
| FUN_001783-T1 | 0.00872613 | 0.22574528 | TBU06884.1 |
| FUN_001784-T1 | 0.01096353 | 0.57532614 | TBU06885.1 |
| FUN_001786-T1 | 0.00543455 | 0.30349985 | TBT99975.1 |
| FUN_001787-T1 | 0.0031715 | 1.95682399 | TBT96785.1 |
| FUN_001788-T1 | 0.00061444 | 0 | TBU04810.1 |
| FUN_001791-T1 | 0.00804343 | 0.77760272 | TBU06689.1 |
| FUN_001792-T1 | 0.02293907 | 1.95896355 | TBU06690.1 |
| FUN_001793-T1 | 0.00266206 | 0.05361688 | TBU06691.1 |
| FUN_001794-T1 | 0.00338776 | 0.10567668 | TBT97346.1 |
| FUN_001796-T1 | 0.02150538 | 0.43814433 | TBT97336.1 |
| FUN_001797-T1 | 0 | NA | TBT98934.1 |
| FUN_001798-T1 | 0.00762375 | 1.75745613 | TBT98937.1 |
| FUN_001801-T1 | 0.04261616 | 0.95194357 | TBT98595.1 |
| FUN_001805-T1 | 0.00244273 | 0.09488014 | TBT99996.1 |
| FUN_001806-T1 | 0.00443905 | 0.47982152 | TBU03435.1 |
| FUN_001808-T1 | 0.01447417 | 1.1627644 | TBU07820.1 |
| FUN_001810-T1 | 0.00389462 | 0.15638712 | TBU07819.1 |
| FUN_001820-T1 | 0.00212399 | 0.38547456 | TBU07151.1 |
| FUN_001823-T1 | 0.00337436 | 0.64231897 | TBU03635.1 |
| FUN_001828-T1 | 0.00142653 | 0.49389845 | TBU07103.1 |
| FUN_001832-T1 | 0.00162738 | 0.16247073 | TBU02785.1 |
| FUN_001833-T1 | 0.00255113 | 4.03222189 | TBT98796.1 |
| FUN_001834-T1 | 0 | NA | TBU00039.1 |
| FUN_001835-T1 | 0.00182268 | 0.1465416 | TBT99941.1 |
| FUN_001837-T1 | 0.0054015 | 0.22762059 | TBT98953.1 |
| FUN_001838-T1 | 0.00482017 | 0 | TBU01572.1 |
| FUN_001840-T1 | 0.00218716 | 7.3493264 | TBU01571.1 |
| FUN_001841-T1 | 0.00160072 | 0.03664025 | TBU03267.1 |
| FUN_001842-T1 | 0.00068106 | 0.21588632 | TBU03268.1 |
| FUN_001846-T1 | 0.00130336 | Inf | TBU06991.1 |
| FUN_001866-T1 | 0.0039444 | 0.06972991 | TBU02914.1 |
| FUN_001867-T1 | 0.00117984 | 0.11630585 | TBU06201.1 |
| FUN_001868-T1 | 0 | NA | TBU01221.1 |
| FUN_001870-T1 | 0.00165882 | Inf | TBU07156.1 |
| FUN_001872-T1 | 0.00857686 | 1.40867048 | TBU07158.1 |
| FUN_001873-T1 | 0 | NA | TBU06008.1 |
| FUN_001879-T1 | 0.00136542 | 0.02345046 | TBU09064.1 |
| FUN_001881-T1 | 0.0022811 | 0.12121129 | TBU09065.1 |
| FUN_001882-T1 | 0.00309299 | 0.15529131 | TBU00345.1 |
| FUN_001885-T1 | 0.00122629 | 0.10530629 | TBU04340.1 |
| FUN_001886-T1 | 0.00468914 | 0.38741913 | TBU05961.1 |
| FUN_001887-T1 | 0.00049438 | 0.15350786 | TBT98682.1 |
| FUN_001888-T1 | 0.00122515 | 0.07099944 | TBU01255.1 |
| FUN_001889-T1 | 0.00077016 | 0 | TBT98420.1 |
| FUN_001890-T1 | 0.00126377 | 0 | TBU01908.1 |
| FUN_001891-T1 | 0.00074428 | 0.50214533 | TBU06922.1 |
| FUN_001892-T1 | 0.00277429 | 0.3266943 | TBU07167.1 |
| FUN_001896-T1 | 0.01206368 | 0.70516451 | TBT98541.1 |
| FUN_001898-T1 | 0.00576974 | 2.93833333 | TBU07811.1 |
| FUN_001899-T1 | 0.01010258 | 0.95492881 | TBU07812.1 |
| FUN_001901-T1 | 0.00563451 | 0.15080064 | TBU07813.1 |
| FUN_001902-T1 | 0.00320962 | 0.22659739 | TBU07637.1 |
| FUN_001903-T1 | 0.00311361 | 0.76700756 | TBU07636.1 |
| FUN_001904-T1 | 0.00247147 | 0.10319353 | TBU07635.1 |
| FUN_001905-T1 | 0.00545925 | 0.40507554 | TBU07634.1 |
| FUN_001906-T1 | 0.00604694 | 0.29901004 | TBU07633.1 |
| FUN_001907-T1 | 0.00371103 | 0.08421553 | TBU07631.1 |
| FUN_001924-T1 | 0.02841609 | 1.55070085 | TBU04678.1 |
| FUN_001931-T1 | 0.00907668 | 0.3806374 | TBU06189.1 |
| FUN_001933-T1 | 0.00495526 | 0.05919762 | TBU04736.1 |
| FUN_001934-T1 | 0.00164299 | 0.32327865 | TBU04735.1 |
| FUN_001935-T1 | 0.00151082 | 0.57866503 | TBU04734.1 |
| FUN_001938-T1 | 0.00166251 | 0.0172994 | TBU07187.1 |
| FUN_001939-T1 | 0.00283274 | 0.42629349 | TBU07186.1 |
| FUN_001940-T1 | 0.00283894 | 0.19584003 | TBU07185.1 |
| FUN_001942-T1 | 0.00215396 | 0.09138499 | TBU07184.1 |
| FUN_001943-T1 | 0.00215054 | 0.53989478 | TBT98399.1 |
| FUN_001944-T1 | 0.00221744 | 0.65585942 | TBU01504.1 |
| FUN_001945-T1 | 0.00278927 | 0.15071459 | TBU04247.1 |
| FUN_001946-T1 | 0.00985266 | 0.48620946 | TBU04248.1 |
| FUN_001948-T1 | 0.00206312 | 0.04335511 | TBU02550.1 |
| FUN_001950-T1 | 0.00812425 | 0.10161833 | TBU02220.1 |
| FUN_001954-T1 | 0.00444985 | 0.32961245 | TBU04090.1 |
| FUN_001955-T1 | 0.00479545 | 0.40099197 | TBU04089.1 |
| FUN_001956-T1 | 0.00083955 | 0 | TBU04088.1 |
| FUN_001958-T1 | 0.02327641 | 1.07429719 | TBU03960.1 |
| FUN_001961-T1 | 0.00184332 | 0.98932777 | TBU01808.1 |
| FUN_001962-T1 | 0.00926072 | 0.1753659 | TBU01807.1 |
| FUN_001963-T1 | 0.00717605 | 0.32017379 | TBU02502.1 |
| FUN_001964-T1 | 0.00086022 | Inf | TBU03485.1 |
| FUN_001965-T1 | 0.00711514 | 0.41933479 | TBU03486.1 |
| FUN_001966-T1 | 0.00603218 | 0.04094035 | TBU03487.1 |
| FUN_001967-T1 | 0.00327011 | 1.4119051 | TBU03488.1 |
| FUN_001968-T1 | 0.00610835 | 0.72187729 | TBU05237.1 |
| FUN_001969-T1 | 0.0027761 | 0.47054954 | TBU04819.1 |
| FUN_001970-T1 | 0.00662929 | 0.26257321 | TBU04820.1 |
| FUN_001971-T1 | 0.00775557 | 0.88045529 | TBT99237.1 |
| FUN_001973-T1 | 0.00210103 | 0.14494597 | TBU05670.1 |
| FUN_001974-T1 | 0.00617328 | 0.49728274 | TBU05669.1 |
| FUN_001975-T1 | 0.00558645 | 0.60323197 | TBU06360.1 |
| FUN_001976-T1 | 0.00722267 | 0.17804542 | TBU06359.1 |
| FUN_001978-T1 | 0.0117525 | 0.47300584 | TBU00146.1 |
| FUN_001980-T1 | 0.00698017 | 0.07998056 | TBT97799.1 |
| FUN_001985-T1 | 0.00332077 | 0.08914517 | TBU07297.1 |
| FUN_001986-T1 | 0.00166225 | Inf | TBU07296.1 |
| FUN_001988-T1 | 0.00392957 | 0.43018124 | TBU00772.1 |
| FUN_001991-T1 | 0.03010753 | 0.70668456 | TBU09345.1 |
| FUN_001995-T1 | 0.00379998 | 0.43571357 | TBT98863.1 |
| FUN_001996-T1 | 0.00116882 | 0 | TBU00529.1 |
| FUN_001997-T1 | 0 | NA | TBU00530.1 |
| FUN_002000-T1 | 0.00110671 | Inf | TBU06793.1 |
| FUN_002001-T1 | 0.00204066 | 0.01342686 | TBU03233.1 |
| FUN_002003-T1 | 0.0079577 | 2.11702413 | TBU05447.1 |
| FUN_002005-T1 | 0.00069936 | 0 | TBU03568.1 |
| FUN_002006-T1 | 0.00481853 | 13.5962468 | TBU03569.1 |
| FUN_002007-T1 | 0.00401714 | 0.54742833 | TBU00658.1 |
| FUN_002008-T1 | 0 | NA | TBT97714.1 |
| FUN_002009-T1 | 0.00090642 | 0 | TBT98642.1 |
| FUN_002010-T1 | 0.00146341 | 2.91650542 | TBT98232.1 |
| FUN_002011-T1 | 0.00025097 | 0.05196404 | TBU04009.1 |
| FUN_002013-T1 | 0.00762745 | 0.35276641 | TBU04010.1 |
| FUN_002015-T1 | 0.00133901 | 0 | TBU08288.1 |
| FUN_002017-T1 | 0.00265538 | 1.05328399 | TBU08287.1 |
| FUN_002019-T1 | 0.00192369 | 0.14917716 | TBU08286.1 |
| FUN_002020-T1 | 0.0027803 | 0.40944404 | TBU08285.1 |
| FUN_002021-T1 | 0.0085044 | 0.2416805 | TBU06558.1 |
| FUN_002036-T1 | 0.00482583 | 0.06054946 | TBU09745.1 |
| FUN_002042-T1 | 0.06110278 | 0.92969951 | TBU04183.1 |
| FUN_002047-T1 | 0.0037691 | 0.20109103 | TBU00841.1 |
| FUN_002053-T1 | 0.00143936 | 0.31070154 | TBU05169.1 |
| FUN_002058-T1 | 0.00210188 | 0.40149161 | TBU02323.1 |
| FUN_002059-T1 | 0.00585787 | 0.46714034 | TBU02324.1 |
| FUN_002060-T1 | 0.00047438 | 0 | TBU02860.1 |
| FUN_002061-T1 | 0.01760132 | 0.8649861 | TBU05491.1 |
| FUN_002065-T1 | 0.0054792 | 0.27519524 | TBU07859.1 |
| FUN_002072-T1 | 0.00173032 | 0 | TBT98505.1 |
| FUN_002073-T1 | 0.00805763 | 2.23627336 | TBU06453.1 |
| FUN_002074-T1 | 0 | NA | TBU06452.1 |
| FUN_002077-T1 | 0.00537157 | 0.55643515 | TBU06451.1 |
| FUN_002081-T1 | 0.00432653 | 1.02851319 | TBU02347.1 |
| FUN_002086-T1 | 0.00619571 | 0.07669027 | TBU04125.1 |
| FUN_002088-T1 | 0.00408375 | 0.14172583 | TBU06867.1 |
| FUN_002089-T1 | 0.00320316 | 0.07200956 | TBU06866.1 |
| FUN_002094-T1 | 0.01457587 | 0.58963633 | TBT98091.1 |
| FUN_002101-T1 | 0.00665478 | 0.5568594 | TBT98627.1 |
| FUN_002108-T1 | 0.00680808 | 0.20012455 | TBU06060.1 |
| FUN_002115-T1 | 0.01081989 | 1.18415712 | TBT97300.1 |
| FUN_002117-T1 | 0.00964803 | 0.63420295 | TBU00443.1 |
| FUN_002119-T1 | 0.00911418 | 0.5411324 | TBU03473.1 |
| FUN_002121-T1 | 0.00910206 | 0.08236859 | TBU00828.1 |
| FUN_002123-T1 | 0.01828262 | 0.30516507 | TBT99719.1 |
| FUN_002126-T1 | 0.00296993 | 0.06767555 | TBU07424.1 |
| FUN_002137-T1 | 0.00422183 | 0 | TBU08839.1 |
| FUN_002138-T1 | 0.00272401 | 0.19444657 | TBU08840.1 |
| FUN_002139-T1 | 0.00637097 | 0.58676878 | TBU08841.1 |
| FUN_002140-T1 | 0.00994842 | 0.49905032 | TBU08842.1 |
| FUN_002142-T1 | 0.0047559 | 0.32383712 | TBU08843.1 |
| FUN_002143-T1 | 0.00796495 | 15.2084875 | TBU08844.1 |
| FUN_002144-T1 | 0.00334528 | 0.13442367 | TBU08345.1 |
| FUN_002145-T1 | 0.02622101 | 0.98183614 | TBU08344.1 |
| FUN_002146-T1 | 0.0028499 | 0.97064076 | TBU08343.1 |
| FUN_002147-T1 | 0.00654235 | 0.0704077 | TBU08342.1 |
| FUN_002148-T1 | 0.00677608 | 0.5765258 | TBU08341.1 |
| FUN_002149-T1 | 0.00107196 | 0.0164451 | TBU08340.1 |
| FUN_002150-T1 | 0.00886466 | 0.90269976 | TBU08339.1 |
| FUN_002151-T1 | 0.00641223 | 0.22895347 | TBU08338.1 |
| FUN_002152-T1 | 0.00939336 | 0.45494857 | TBT97947.1 |
| FUN_002154-T1 | 0.00097703 | Inf | TBU04531.1 |
| FUN_002155-T1 | 0.01864817 | 1.30810628 | TBU04530.1 |
| FUN_002162-T1 | 0.00775405 | 4.82463619 | TBU06802.1 |
| FUN_002172-T1 | 0.0061563 | 0.51510344 | TBU07577.1 |
| FUN_002173-T1 | 0.00254747 | 0.20161208 | TBU07576.1 |
| FUN_002174-T1 | 0.00138509 | 0.12255588 | TBU07575.1 |
| FUN_002175-T1 | 0.013154 | 1.02079743 | TBU07574.1 |
| FUN_002177-T1 | 0.00929428 | 0.39373598 | TBU09196.1 |
| FUN_002178-T1 | 0.00528686 | 0.49094058 | TBU09197.1 |
| FUN_002179-T1 | 0.00245776 | 0.1848368 | TBU09198.1 |
| FUN_002180-T1 | 0.00300773 | 0.16877739 | TBU09199.1 |
| FUN_002181-T1 | 0.00584677 | 0.20339269 | TBU09200.1 |
| FUN_002182-T1 | 0.00568882 | 0.34552707 | TBU09201.1 |
| FUN_002183-T1 | 0.00957077 | 0.60173746 | TBU09202.1 |
| FUN_002191-T1 | 0.01076212 | 2.4673237 | TBU07645.1 |
| FUN_002194-T1 | 0.00695565 | 0.22368859 | TBU06979.1 |
| FUN_002195-T1 | 0.00508902 | 0.20294605 | TBU06980.1 |
| FUN_002196-T1 | 0.00236559 | Inf | TBU06981.1 |
| FUN_002197-T1 | 0.00459472 | 0.18804182 | TBU06982.1 |
| FUN_002198-T1 | 0.01487795 | 2.26198488 | TBU06983.1 |
| FUN_002205-T1 | 0.01152953 | 0.6918436 | TBU01306.1 |
| FUN_002207-T1 | 0.00258065 | 0.84987169 | TBU05173.1 |
| FUN_002208-T1 | 0.00174268 | 0 | TBU05174.1 |
| FUN_002209-T1 | 0.00405053 | 0.14511989 | TBU01408.1 |
| FUN_002213-T1 | 0.0023291 | Inf | TBU03589.1 |
| FUN_002214-T1 | 0.00230803 | 0.04365792 | TBU02573.1 |
| FUN_002215-T1 | 0 | NA | TBU02572.1 |
| FUN_002217-T1 | 0.0046584 | 0.10582639 | TBU06500.1 |
| FUN_002220-T1 | 0.00021944 | 0 | TBU06498.1 |
| FUN_002221-T1 | 0.00368494 | 0.28130529 | TBU05392.1 |
| FUN_002223-T1 | 0.00169485 | 0.10782279 | TBT99603.1 |
| FUN_002224-T1 | 0.00294594 | 0.31622892 | TBT98920.1 |
| FUN_002236-T1 | 0.00289993 | 0.1576408 | TBU03153.1 |
| FUN_002237-T1 | 0.00181969 | 0.36176694 | TBU07110.1 |
| FUN_002239-T1 | 0.00328254 | 0.61013957 | TBU07109.1 |
| FUN_002240-T1 | 0.00133811 | 0.14577115 | TBU07108.1 |
| FUN_002241-T1 | 0.001899 | 0.61756372 | TBU07107.1 |
| FUN_002244-T1 | 0.00375668 | 0.31959147 | TBU05786.1 |
| FUN_002245-T1 | 0.0018124 | 1.63480826 | TBU05787.1 |
| FUN_002246-T1 | 0.00588496 | 0.25435448 | TBU05788.1 |
| FUN_002247-T1 | 0.00225606 | 0.1206567 | TBU05789.1 |
| FUN_002252-T1 | 0.0053099 | 0.09011289 | TBU06134.1 |
| FUN_002253-T1 | 0.00614304 | 1.33612925 | TBU06135.1 |
| FUN_002254-T1 | 0.00463178 | 0.21242127 | TBU06136.1 |
| FUN_002257-T1 | 0.00419065 | 0.2250073 | TBU06934.1 |
| FUN_002260-T1 | 0.01109158 | 0.20822683 | TBT98187.1 |
| FUN_002272-T1 | 0.00371929 | 0.3353901 | TBU07331.1 |
| FUN_002274-T1 | 0.00043155 | 0.05091664 | TBU01901.1 |
| FUN_002275-T1 | 0.00440258 | 0.72759356 | TBU01900.1 |
| FUN_002280-T1 | 0.00703669 | 0.28963249 | TBU08922.1 |
| FUN_002283-T1 | 0.00746916 | 0.24517136 | TBU08923.1 |
| FUN_002284-T1 | 0.0002217 | Inf | TBU08924.1 |
| FUN_002285-T1 | 0.00217893 | 0.070991 | TBU02494.1 |
| FUN_002292-T1 | 0.0031645 | 1.46813016 | TBU08878.1 |
| FUN_002293-T1 | 0.00919082 | 0.28981672 | TBU08879.1 |
| FUN_002302-T1 | 0.00161575 | 0.23268129 | TBU01628.1 |
| FUN_002303-T1 | 0.00101234 | 0.12690571 | TBU03972.1 |
| FUN_002304-T1 | 0.00128233 | 0.3071144 | TBU03973.1 |
| FUN_002305-T1 | 0.02372619 | 0.65910704 | TBU02871.1 |
| FUN_002307-T1 | 0.00168516 | 0.12724608 | TBU01368.1 |
| FUN_002308-T1 | 0.00275986 | 0.11380847 | TBT98764.1 |
| FUN_002310-T1 | 0.00113464 | 0.03035588 | TBU06338.1 |
| FUN_002312-T1 | 0.00367055 | 0.2012746 | TBU06337.1 |
| FUN_002313-T1 | 0.00110995 | 0.06028287 | TBU06336.1 |
| FUN_002314-T1 | 0.00460023 | 0.4519723 | TBU06334.1 |
| FUN_002315-T1 | 0.00329508 | 2.59744904 | TBU00603.1 |
| FUN_002316-T1 | 0.00593638 | 0.46859034 | TBU00602.1 |
| FUN_002317-T1 | 0.00491655 | 0.21855745 | TBT97127.1 |
| FUN_002318-T1 | 0.0031209 | 0.49656046 | TBT99293.1 |
| FUN_002319-T1 | 0.00432947 | 0.06592636 | TBT99292.1 |
| FUN_002320-T1 | 0.0033258 | 0.12099848 | TBU03839.1 |
| FUN_002323-T1 | 0.00533106 | 0.39974412 | TBT98464.1 |
| FUN_002324-T1 | 0.0053514 | 0.18102155 | TBT97045.1 |
| FUN_002326-T1 | 0.00754072 | 0.99835768 | TBU04958.1 |
| FUN_002327-T1 | 0.00353798 | 0.02288009 | TBU04957.1 |
| FUN_002328-T1 | 0.00512803 | 0.72676928 | TBU04956.1 |
| FUN_002334-T1 | 0.00168556 | Inf | TBU03863.1 |
| FUN_002338-T1 | 0.00681821 | 1.16732205 | TBU03865.1 |
| FUN_002339-T1 | 0.01860812 | 0.76563383 | TBU03864.1 |
| FUN_002341-T1 | 0.00131271 | 0.15951525 | TBU05569.1 |
| FUN_002342-T1 | 0.00410145 | 0.25293983 | TBU05568.1 |
| FUN_002343-T1 | 0.005582 | 0.14385822 | TBU05567.1 |
| FUN_002344-T1 | 0.0038048 | 0.20920923 | TBU05566.1 |
| FUN_002348-T1 | 0.0032497 | 0.0642139 | TBU07024.1 |
| FUN_002349-T1 | 0.00184332 | Inf | TBU07023.1 |
| FUN_002358-T1 | 0.0096126 | 0.37851073 | TBT99050.1 |
| FUN_002359-T1 | 0.00356394 | 0.23179343 | TBU02139.1 |
| FUN_002360-T1 | 0.00391781 | 0.08501458 | TBT99612.1 |
| FUN_002361-T1 | 0.00999305 | 0.54788267 | TBT99613.1 |
| FUN_002362-T1 | 0.00530924 | 0.37259595 | TBU07660.1 |
| FUN_002364-T1 | 0.02098648 | 1.20099913 | TBU07659.1 |
| FUN_002365-T1 | 0.00364112 | 0.08337512 | TBU07403.1 |
| FUN_002366-T1 | 0.00141577 | 1.18497486 | TBU07404.1 |
| FUN_002369-T1 | 0.0038527 | 0.28418528 | TBU07798.1 |
| FUN_002372-T1 | 0.00211708 | 0.12232381 | TBU02386.1 |
| FUN_002373-T1 | 0.00541741 | 0.31269141 | TBU02387.1 |
| FUN_002376-T1 | 0.00095024 | 0.08585165 | TBU06575.1 |
| FUN_002387-T1 | 0.01274539 | 0.50313253 | TBU09755.1 |
| FUN_002388-T1 | 0.00315185 | 0.48388811 | TBU09756.1 |
| FUN_002390-T1 | 0.01957801 | 0.49750806 | TBU05836.1 |
| FUN_002391-T1 | 0.00678876 | 2.01881712 | TBU05837.1 |
| FUN_002393-T1 | 0.00162485 | 0 | TBU05838.1 |
| FUN_002394-T1 | 0.00277778 | 0.44667162 | TBU06606.1 |
| FUN_002396-T1 | 0.01649784 | 0.04330026 | TBT96883.1 |
| FUN_002397-T1 | 0.00514496 | 0.03770908 | TBU03165.1 |
| FUN_002399-T1 | 0.01178121 | 0.09496355 | TBT97743.1 |
| FUN_002401-T1 | 0.00541278 | 0.22944015 | TBU00124.1 |
| FUN_002403-T1 | 0.00673639 | 0.28528272 | TBU00677.1 |
| FUN_002406-T1 | 0.00649324 | 0.24277765 | TBT99006.1 |
| FUN_002408-T1 | 0.02069576 | 0.50720331 | TBU05811.1 |
| FUN_002414-T1 | 0.00092801 | 0.40318228 | TBU08983.1 |
| FUN_002419-T1 | 0.00786901 | 0.09890278 | TBU06751.1 |
| FUN_002420-T1 | 0 | NA | TBU06750.1 |
| FUN_002421-T1 | 0.0020893 | 0.49722648 | TBU06749.1 |
| FUN_002422-T1 | 0.00243467 | 0 | TBU06748.1 |
| FUN_002423-T1 | 0.00060864 | Inf | TBU06747.1 |
| FUN_002449-T1 | 0.00022095 | 0.13372098 | TBU01324.1 |
| FUN_002450-T1 | 0.0068271 | 3.02718534 | TBU00375.1 |
| FUN_002452-T1 | 0.00237162 | 2.29554833 | TBU07344.1 |
| FUN_002453-T1 | 0.001136 | 0.28677583 | TBU07345.1 |
| FUN_002454-T1 | 0.00109188 | 0 | TBU07346.1 |
| FUN_002455-T1 | 0.00104596 | 0.08435493 | TBU07347.1 |
| FUN_002459-T1 | 0 | NA | TBU09095.1 |
| FUN_002462-T1 | 0.00039121 | 0.44749714 | TBT99447.1 |
| FUN_002463-T1 | 0.00072389 | 0.41331121 | TBU04456.1 |
| FUN_002464-T1 | 0.00043889 | 0.44081414 | TBU04455.1 |
| FUN_002466-T1 | 0.00100439 | 0.21916901 | TBT98004.1 |
| FUN_002484-T1 | 0.00067978 | 0.26930171 | TBU06481.1 |
| FUN_002487-T1 | 0.00015198 | Inf | TBU08361.1 |
| FUN_002489-T1 | 0.00065068 | 0.68261338 | TBU08363.1 |
| FUN_002492-T1 | 0.00117636 | 0.55886364 | TBU08366.1 |
| FUN_002497-T1 | 0.00242561 | 0.31490957 | TBU06159.1 |
| FUN_002499-T1 | 0.00054981 | 0 | TBU09451.1 |
| FUN_002506-T1 | 0.00149834 | 0.23552072 | TBU04833.1 |
| FUN_002507-T1 | 0.00046891 | 0 | TBU04832.1 |
| FUN_002508-T1 | 0.00102876 | 0.20025587 | TBU04831.1 |
| FUN_002509-T1 | 0.00055677 | 0.13058106 | TBU03643.1 |
| FUN_002510-T1 | 0.00296979 | 1.09173738 | TBU03642.1 |
| FUN_002515-T1 | 0.00148271 | 0.06323129 | TBU08956.1 |
| FUN_002516-T1 | 0.00433028 | 0.04287326 | TBU08957.1 |
| FUN_002517-T1 | 0.00389355 | 0.09594299 | TBU08958.1 |
| FUN_002518-T1 | 0.00489672 | 0.49647083 | TBU08959.1 |
| FUN_002522-T1 | 0.00491948 | 0.20931972 | TBU02781.1 |
| FUN_002524-T1 | 0.00364811 | 0 | TBU02782.1 |
| FUN_002525-T1 | 0.00304175 | 0 | TBU02783.1 |
| FUN_002529-T1 | 0.01689742 | 0.22450809 | TBU05272.1 |
| FUN_002530-T1 | 0.00267405 | 0.17261685 | TBU05271.1 |
| FUN_002531-T1 | 0.00041816 | 1.53008209 | TBU05270.1 |
| FUN_002534-T1 | 0.00266159 | 1.34139431 | TBU08088.1 |
| FUN_002535-T1 | 0.00425787 | 0.08380832 | TBU08089.1 |
| FUN_002536-T1 | 0.00122971 | 0.18579681 | TBU08090.1 |
| FUN_002537-T1 | 0.00046293 | 0 | TBU08091.1 |
| FUN_002540-T1 | 0.0052951 | 0.83989601 | TBU01290.1 |
| FUN_002541-T1 | 0.03520674 | 0.85500881 | TBT97436.1 |
| FUN_002542-T1 | 0.00083445 | 0.5979398 | TBT99912.1 |
| FUN_002543-T1 | 0.00523478 | 2.97589035 | TBT99913.1 |
| FUN_002544-T1 | 0.00137579 | 0.27177282 | TBU00992.1 |
| FUN_002546-T1 | 0.00747985 | 0.29021975 | TBU04714.1 |
| FUN_002548-T1 | 0.00264787 | 0.03986988 | TBU06910.1 |
| FUN_002549-T1 | 0.00459057 | 0.09511556 | TBU06911.1 |
| FUN_002550-T1 | 0.00410753 | 0.12554711 | TBU06912.1 |
| FUN_002554-T1 | 0.00329464 | 0.02141087 | TBU05862.1 |
| FUN_002555-T1 | 0.00191159 | Inf | TBU05863.1 |
| FUN_002556-T1 | 0.0021525 | 0.02090025 | TBU05864.1 |
| FUN_002557-T1 | 0.00310304 | 0.06138668 | TBU05105.1 |
| FUN_002558-T1 | 0.00904954 | 0.37346638 | TBU05104.1 |
| FUN_002559-T1 | 0.00448207 | 0.16813733 | TBU05384.1 |
| FUN_002561-T1 | 0.00220771 | 0.13742522 | TBU05383.1 |
| FUN_002562-T1 | 0.00121864 | 0.1220909 | TBU05382.1 |
| FUN_002563-T1 | 0.00508309 | 0.60183043 | TBU07275.1 |
| FUN_002564-T1 | 0.00342888 | 0.35151563 | TBU07274.1 |
| FUN_002565-T1 | 0.00371536 | 0.08803179 | TBU07273.1 |
| FUN_002566-T1 | 0.003838 | 0.15581858 | TBU07272.1 |
| FUN_002567-T1 | 0.00010003 | Inf | TBU07271.1 |
| FUN_002569-T1 | 0.00166817 | 0 | TBU03319.1 |
| FUN_002570-T1 | 0.00718783 | 0.91624095 | TBU07044.1 |
| FUN_002571-T1 | 0.00523884 | 0.15230465 | TBU07043.1 |
| FUN_002572-T1 | 0.00586081 | 0.06831994 | TBU07042.1 |
| FUN_002573-T1 | 0.00136746 | 0.01067287 | TBU07041.1 |
| FUN_002574-T1 | 0.00516584 | 0.18740549 | TBU07040.1 |
| FUN_002575-T1 | 0.0049656 | 0.0881459 | TBU07039.1 |
| FUN_002583-T1 | 0.00191693 | 0.51438885 | TBU05611.1 |
| FUN_002584-T1 | 0.00524108 | 0.26862195 | TBU05612.1 |
| FUN_002585-T1 | 0.00178687 | 0.10005357 | TBU01109.1 |
| FUN_002586-T1 | 0.00214925 | 0.08885512 | TBT98611.1 |
| FUN_002588-T1 | 0.00481651 | 0.19757072 | TBU07442.1 |
| FUN_002589-T1 | 0.00474221 | 0.53405445 | TBU07443.1 |
| FUN_002593-T1 | 0.00051613 | 0 | TBU05990.1 |
| FUN_002594-T1 | 0.00334443 | 0.08548621 | TBU05991.1 |
| FUN_002597-T1 | 0.00455483 | 0.08627321 | TBU07000.1 |
| FUN_002598-T1 | 0.00221342 | 0.01405041 | TBU06999.1 |
| FUN_002603-T1 | 0.00368379 | 0.11810777 | TBU07248.1 |
| FUN_002604-T1 | 0.0001593 | Inf | TBU07247.1 |
| FUN_002606-T1 | 0.00320226 | 0.47641562 | TBU04072.1 |
| FUN_002608-T1 | 0.00554612 | 7.434903 | TBU08815.1 |
| FUN_002609-T1 | 0.01485826 | 1.12550631 | TBU08816.1 |
| FUN_002610-T1 | 0.00158133 | 0.0371804 | TBU08818.1 |
| FUN_002611-T1 | 0.00619954 | 0.20648255 | TBU08819.1 |
| FUN_002613-T1 | 0.01150314 | 0.35409029 | TBU08820.1 |
| FUN_002614-T1 | 0.00655164 | 0.28453965 | TBU08821.1 |
| FUN_002615-T1 | 0.00514259 | 0.10569173 | TBU08822.1 |
| FUN_002618-T1 | 0.00397263 | 0.26894673 | TBU07773.1 |
| FUN_002623-T1 | 0.00240273 | 0.11377564 | TBU05658.1 |
| FUN_002624-T1 | 0.00238547 | 0.73976383 | TBU00102.1 |
| FUN_002625-T1 | 0.00261612 | 0.01969862 | TBU08833.1 |
| FUN_002626-T1 | 0.00277636 | 0.61213723 | TBU08834.1 |
| FUN_002632-T1 | 0.00509095 | 0.17913904 | TBU03079.1 |
| FUN_002635-T1 | 0.00323486 | 0.01257794 | TBU03416.1 |
| FUN_002638-T1 | 0.00494112 | 0.24924248 | TBU07848.1 |
| FUN_002639-T1 | 0.0049288 | 0.19220076 | TBU07847.1 |
| FUN_002640-T1 | 0.00349826 | 0.12447203 | TBU01396.1 |
| FUN_002641-T1 | 0.00210868 | 0.09211413 | TBU00691.1 |
| FUN_002642-T1 | 0.00452542 | 0.28166804 | TBU08657.1 |
| FUN_002643-T1 | 0.00188373 | 0.04466862 | TBU08656.1 |
| FUN_002646-T1 | 0.00326428 | 0.13623253 | TBU08655.1 |
| FUN_002647-T1 | 0.00296759 | 0.04577045 | TBU08654.1 |
| FUN_002649-T1 | 0.00226229 | 0.0874765 | TBU08653.1 |
| FUN_002650-T1 | 0 | NA | TBU08652.1 |
| FUN_002651-T1 | 0.00691003 | 0.1085122 | TBU02799.1 |
| FUN_002654-T1 | 0.00215054 | Inf | TBU02188.1 |
| FUN_002656-T1 | 0.00338004 | 0 | TBU03013.1 |
| FUN_002657-T1 | 0.00495351 | 0.02521836 | TBU03014.1 |
| FUN_002658-T1 | 0.00398546 | 0.25376885 | TBU06833.1 |
| FUN_002659-T1 | 0.00432146 | 0.57272378 | TBU06834.1 |
| FUN_002660-T1 | 0.0007611 | 0.03582254 | TBT97234.1 |
| FUN_002661-T1 | 0.00189247 | 0.24563741 | TBU07508.1 |
| FUN_002662-T1 | 0.00153301 | 0.10469322 | TBU07507.1 |
| FUN_002664-T1 | 0.00500397 | 0.19656889 | TBU07506.1 |
| FUN_002669-T1 | 0.00347267 | 0.06864098 | TBU09420.1 |
| FUN_002671-T1 | 0.0023075 | 0.16810272 | TBU09422.1 |
| FUN_002672-T1 | 0.00359065 | 0.08846468 | TBU09423.1 |
| FUN_002673-T1 | 0.00628589 | 0.84433911 | TBU09424.1 |
| FUN_002674-T1 | 0.00034409 | 0 | TBU09425.1 |
| FUN_002676-T1 | 0.00123141 | 0.03966393 | TBU09426.1 |
| FUN_002677-T1 | 0.00573913 | 0.37157837 | TBT99509.1 |
| FUN_002678-T1 | 0.00435304 | 0.1020309 | TBU00024.1 |
| FUN_002680-T1 | 0.00583632 | 0.74302702 | TBU04140.1 |
| FUN_002684-T1 | 0.00536703 | 0.16344263 | TBU06081.1 |
| FUN_002685-T1 | 0.0037779 | 0.26200924 | TBU06082.1 |
| FUN_002687-T1 | 0.00425388 | 0.18394668 | TBU06083.1 |
| FUN_002688-T1 | 0.0028607 | 0.23395989 | TBU06786.1 |
| FUN_002689-T1 | 0.00187305 | 0 | TBU06787.1 |
| FUN_002690-T1 | 0.00503017 | 0.07856023 | TBU06788.1 |
| FUN_002691-T1 | 0.00190298 | 0.2222997 | TBT99172.1 |
| FUN_002693-T1 | 0.00135292 | 0 | TBT99715.1 |
| FUN_002695-T1 | 0.0039786 | 0.15455271 | TBU06171.1 |
| FUN_002697-T1 | 0.00471499 | 0.10724676 | TBU06169.1 |
| FUN_002698-T1 | 0.002 | 0.04583359 | TBU03355.1 |
| FUN_002699-T1 | 0.00507766 | 0.41383884 | TBU03354.1 |
| FUN_002701-T1 | 0.00271524 | 0 | TBU01956.1 |
| FUN_002702-T1 | 0.0178118 | 0.77058091 | TBU01957.1 |
| FUN_002703-T1 | 0.00455862 | 0 | TBU09547.1 |
| FUN_002704-T1 | 0.0039404 | 0 | TBU09546.1 |
| FUN_002705-T1 | 0.00909001 | 0.38464858 | TBU09545.1 |
| FUN_002706-T1 | 0.01216374 | 0.28134394 | TBU09544.1 |
| FUN_002707-T1 | 0.00573735 | 0.1659226 | TBU09543.1 |
| FUN_002708-T1 | 0.00739049 | 0.2758692 | TBU09542.1 |
| FUN_002709-T1 | 0.00170837 | 0.01700793 | TBU09541.1 |
| FUN_002710-T1 | 0.00555954 | 0.04596008 | TBU09540.1 |
| FUN_002711-T1 | 0.00794839 | 0.65301004 | TBU09539.1 |
| FUN_002716-T1 | 0.02314195 | 1.61069011 | TBT97214.1 |
| FUN_002723-T1 | 0.00730029 | 0.55819623 | TBU07028.1 |
| FUN_002724-T1 | 0.00953675 | 1.08214399 | TBU07027.1 |
| FUN_002727-T1 | 0.00403043 | 0.03512877 | TBU00451.1 |
| FUN_002728-T1 | 0.00827637 | 0.49754459 | TBU00452.1 |
| FUN_002729-T1 | 0.00500229 | 0.13300064 | TBU03112.1 |
| FUN_002739-T1 | 0.00399557 | 0.30296095 | TBU09135.1 |
| FUN_002740-T1 | 0.00508048 | 0.59607125 | TBU09134.1 |
| FUN_002742-T1 | 0.01037966 | 0.5263834 | TBU09133.1 |
| FUN_002746-T1 | 0.0074541 | 1.24510049 | TBU09473.1 |
| FUN_002749-T1 | 0.00901098 | 0.76709114 | TBU09472.1 |
| FUN_002750-T1 | 0.00367474 | 0.21611733 | TBU09x471.1 |
| FUN_002751-T1 | 0.00356226 | 0.4085377 | TBT99095.1 |
| FUN_002756-T1 | 0.00538263 | 0.22244547 | TBU07779.1 |
| FUN_002758-T1 | 0.00614635 | 0.16548116 | TBT98965.1 |
| FUN_002761-T1 | 0.00490261 | 0.41554349 | TBU01054.1 |
| FUN_002763-T1 | 0.0034178 | 0.08113574 | TBU01716.1 |
| FUN_002764-T1 | 0.00150538 | 0.34189976 | TBT99928.1 |
| FUN_002777-T1 | 0.0003471 | Inf | TBU00045.1 |
| FUN_002780-T1 | 0.00296404 | 0.07997936 | TBU02417.1 |
| FUN_002783-T1 | 0.00445707 | 0.12563471 | TBU05303.1 |
| FUN_002784-T1 | 0.00183092 | 0.31605715 | TBU05302.1 |
| FUN_002785-T1 | 0.00264023 | 0.31166816 | TBU01042.1 |
| FUN_002786-T1 | 0.00206791 | 0.07540387 | TBT98427.1 |
| FUN_002787-T1 | 0.00042405 | 0.18573056 | TBT99366.1 |
| FUN_002788-T1 | 0.00098865 | 0.12443274 | TBU07980.1 |
| FUN_002789-T1 | 0.00151803 | 0.16032971 | TBU07981.1 |
| FUN_002790-T1 | 0.00869123 | 0.75097235 | TBU07982.1 |
| FUN_002792-T1 | 0.00174661 | 0.02995952 | TBT99520.1 |
| FUN_002793-T1 | 0.00471887 | 0.26715215 | TBU01614.1 |
| FUN_002795-T1 | 0.00142735 | 0.36648055 | TBU09042.1 |
| FUN_002796-T1 | 0.00334698 | 0.1184447 | TBU09040.1 |
| FUN_002798-T1 | 0.00404221 | 0.03832461 | TBU09039.1 |
| FUN_002800-T1 | 0.00014831 | 0 | TBU09038.1 |
| FUN_002801-T1 | 0.01290323 | 0.30067778 | TBU09037.1 |
| FUN_002803-T1 | 0.0042938 | 0.14714215 | TBT99220.1 |
| FUN_002806-T1 | 0.00200305 | 1.10442765 | TBU01647.1 |
| FUN_002811-T1 | 0.00355136 | 1.42449188 | TBU06069.1 |
| FUN_002812-T1 | 0.03287771 | 1.02234479 | TBU06068.1 |
| FUN_002813-T1 | 0.00021084 | 0 | TBU05847.1 |
| FUN_002815-T1 | 0.00042076 | Inf | TBU07738.1 |
| FUN_002816-T1 | 0.00277821 | 0.19088766 | TBU07737.1 |
| FUN_002817-T1 | 0.00215054 | 0.2695525 | TBU07736.1 |
| FUN_002818-T1 | 0.00138744 | 0.02889064 | TBU07735.1 |
| FUN_002819-T1 | 0.002096 | 0.1397873 | TBU07734.1 |
| FUN_002822-T1 | 0.00161441 | 0.01564679 | TBU07415.1 |
| FUN_002823-T1 | 0.00385751 | 0.37602649 | TBU07414.1 |
| FUN_002824-T1 | 0.00158935 | 0.03991255 | TBU07413.1 |
| FUN_002826-T1 | 0.01062394 | 0.54896669 | TBU02334.1 |
| FUN_002828-T1 | 0 | NA | TBU05985.1 |
| FUN_002830-T1 | 0.00030423 | 0 | TBU04398.1 |
| FUN_002832-T1 | 0.00545767 | 0.61264344 | TBU03724.1 |
| FUN_002833-T1 | 0.01469076 | 0.62487991 | TBU03725.1 |
| FUN_002835-T1 | 0.00339895 | 0.18310846 | TBU03764.1 |
| FUN_002836-T1 | 0.00021944 | Inf | TBU05695.1 |
| FUN_002837-T1 | 0.00257658 | 0.26467899 | TBU05696.1 |
| FUN_002841-T1 | 0.00343978 | 0.11312958 | TBU07866.1 |
| FUN_002844-T1 | 0.00625184 | 0.83826056 | TBU00366.1 |
| FUN_002846-T1 | 0.0022903 | 0.23492502 | TBU02666.1 |
| FUN_002847-T1 | 0.00645161 | 0.28978267 | TBU02665.1 |
| FUN_002848-T1 | 0.00122358 | Inf | TBT99242.1 |
| FUN_002850-T1 | 0.00109635 | 0 | TBU05859.1 |
| FUN_002851-T1 | 0.00215707 | 0.0959592 | TBU05858.1 |
| FUN_002853-T1 | 0.00127045 | 0.07819624 | TBU02819.1 |
| FUN_002855-T1 | 0.00863449 | 0.39568614 | TBU03809.1 |
| FUN_002856-T1 | 0.00113936 | 0.25051421 | TBU03810.1 |
| FUN_002857-T1 | 0.00114695 | 1.49820144 | TBU01217.1 |
| FUN_002858-T1 | 7.27E-05 | 0 | TBU01216.1 |
| FUN_002865-T1 | 7.94E-05 | Inf | TBU04777.1 |
| FUN_002866-T1 | 0.00158094 | 0 | TBU04776.1 |
| FUN_002869-T1 | 0.00784492 | 0.58404846 | TBU09032.1 |
| FUN_002870-T1 | 0.00130902 | 0.07094706 | TBU09031.1 |
| FUN_002872-T1 | 0.00494948 | 0.11766511 | TBU09125.1 |
| FUN_002873-T1 | 0.0031097 | 0.53171521 | TBU09126.1 |
| FUN_002874-T1 | 0.0090606 | 1.01311806 | TBU09127.1 |
| FUN_002878-T1 | 0.00212766 | 0.04746893 | TBT97031.1 |
| FUN_002879-T1 | 0.01285526 | 0.51710998 | TBU05771.1 |
| FUN_002880-T1 | 0.00283043 | 0.14579285 | TBU05770.1 |
| FUN_002881-T1 | 0.00270196 | 1.31311434 | TBU05769.1 |
| FUN_002882-T1 | 0.01982691 | 0.59469634 | TBU05767.1 |
| FUN_002888-T1 | 0.01066205 | 0.58482779 | TBT98015.1 |
| FUN_002890-T1 | 0.0055359 | 0.88407042 | TBT99700.1 |
| FUN_002901-T1 | 0.00746182 | 0.60766544 | TBT99731.1 |
| FUN_002906-T1 | 0.00585539 | 0.8409947 | TBU07047.1 |
| FUN_002907-T1 | 0.00283154 | 0.69289062 | TBU02374.1 |
| FUN_002909-T1 | 0.00346524 | 0.03305775 | TBT99891.1 |
| FUN_002910-T1 | 0.03537058 | 0.66574251 | TBU01558.1 |
| FUN_002912-T1 | 0.00404671 | 0.13401386 | TBU02224.1 |
| FUN_002914-T1 | 0.01051422 | 0.3917009 | TBT97921.1 |
| FUN_002915-T1 | 0.00968773 | 0.95822097 | TBT99124.1 |
| FUN_002924-T1 | 0.01099066 | 0.46076638 | TBT97661.1 |
| FUN_002925-T1 | 0.00101801 | 1.77507777 | TBU08243.1 |
| FUN_002934-T1 | 0.00355768 | Inf | TBU09769.1 |
| FUN_002939-T1 | 0.00781186 | 0.46496492 | TBU09767.1 |
| FUN_002941-T1 | 0.00204258 | 0.83033557 | TBU09765.1 |
| FUN_002942-T1 | 0.00589454 | 0.5520989 | TBU02539.1 |
| FUN_002943-T1 | 0.00369469 | 0.28869629 | TBU00198.1 |
| FUN_002944-T1 | 0.00342212 | 0.17324496 | TBU00197.1 |
| FUN_002945-T1 | 0.00168505 | 0.05266089 | TBU00282.1 |
| FUN_002946-T1 | 0.00385115 | 0.23304661 | TBU01465.1 |
| FUN_002948-T1 | 0.00205545 | 0.3747615 | TBT98734.1 |
| FUN_002949-T1 | 0.00636478 | 0.36001244 | TBT97924.1 |
| FUN_002950-T1 | 0.00494943 | 0.0418873 | TBU03814.1 |
| FUN_002955-T1 | 0.01405139 | 0.8735422 | TBU01184.1 |
| FUN_002956-T1 | 0.00465785 | 0.18007072 | TBU01638.1 |
| FUN_002957-T1 | 0.00125746 | 0.15587518 | TBU05953.1 |
| FUN_002958-T1 | 0.00516771 | 0.42073958 | TBU05954.1 |
| FUN_002959-T1 | 0.00356945 | 0.07963511 | TBU05956.1 |
| FUN_002960-T1 | 0.00636757 | 0.27085046 | TBU00977.1 |
| FUN_002961-T1 | 0.00790765 | 0.43344703 | TBU00976.1 |
| FUN_002962-T1 | 0.00501459 | 1.00121386 | TBU06722.1 |
| FUN_002965-T1 | 0.00351785 | 0.26459467 | TBT99541.1 |
| FUN_002969-T1 | 0.00454109 | 0.80068172 | TBU07920.1 |
| FUN_002972-T1 | 0.02442911 | 0.90988577 | TBT99763.1 |
| FUN_002982-T1 | 0.00021505 | Inf | TBU01797.1 |
| FUN_002984-T1 | 0 | NA | TBU06003.1 |
| FUN_002985-T1 | 0.00571552 | 0.45297215 | TBU06002.1 |
| FUN_002986-T1 | 0.0005191 | 0.53079113 | TBU06001.1 |
| FUN_002987-T1 | 0.00230686 | 0.00315085 | TBU01660.1 |
| FUN_002988-T1 | 0.00777463 | 0.56210182 | TBU01659.1 |
| FUN_002989-T1 | 0.00452745 | 0.14597197 | TBT96766.1 |
| FUN_002990-T1 | 0.00601569 | 0.4845049 | TBU06094.1 |
| FUN_002994-T1 | 0.00348368 | 0.09423244 | TBU09143.1 |
| FUN_002998-T1 | 0.00106945 | 0.01621886 | TBU02423.1 |
| FUN_002999-T1 | 0.00071324 | 0 | TBU02987.1 |
| FUN_003007-T1 | 0.00842045 | 0.40572543 | TBU09146.1 |
| FUN_003009-T1 | 0.00178611 | 0.4927096 | TBU02090.1 |
| FUN_003010-T1 | 0.00114797 | 0.3430202 | TBU00839.1 |
| FUN_003013-T1 | 0.00114444 | 0.12618835 | TBU05541.1 |
| FUN_003021-T1 | 0.00829552 | 0.35162853 | TBU02709.1 |
| FUN_003022-T1 | 0.00245249 | 0.00801742 | TBU03090.1 |
| FUN_003023-T1 | 0.00391705 | 0.01714025 | TBU03091.1 |
| FUN_003025-T1 | 0.00275285 | 0.28074214 | TBU02244.1 |
| FUN_003027-T1 | 0.00417784 | 0.08497294 | TBU03366.1 |
| FUN_003028-T1 | 0.00195287 | 0.49470204 | TBU03367.1 |
| FUN_003034-T1 | 0.01515457 | 1.00245051 | TBU01159.1 |
| FUN_003043-T1 | 0.00502918 | 0.12138157 | TBU08142.1 |
| FUN_003044-T1 | 0.00168055 | 0.51885683 | TBU08143.1 |
| FUN_003045-T1 | 0.00529425 | 0.03621882 | TBU08144.1 |
| FUN_003046-T1 | 0.00464864 | 0.19629083 | TBU08145.1 |
| FUN_003051-T1 | 0.0031873 | 0.17950276 | TBU05923.1 |
| FUN_003052-T1 | 0.00372931 | 0.23395932 | TBU05922.1 |
| FUN_003053-T1 | 0.01280857 | 0.40917557 | TBU05873.1 |
| FUN_003054-T1 | 0.00437514 | 0.23168534 | TBU05872.1 |
| FUN_003059-T1 | 0.00563451 | 0.3159988 | TBU00242.1 |
| FUN_003060-T1 | 0.00468144 | 0.46300563 | TBU03192.1 |
| FUN_003061-T1 | 0.00441461 | 0.38776515 | TBU03139.1 |
| FUN_003062-T1 | 0.0033308 | 0.10813456 | TBU03254.1 |
| FUN_003063-T1 | 0.00213992 | 0.04473781 | TBU00523.1 |
| FUN_003066-T1 | 0.00579958 | 0.48727403 | TBU08503.1 |
| FUN_003067-T1 | 0.00236089 | 0.12180037 | TBU08504.1 |
| FUN_003069-T1 | 0.0021279 | Inf | TBU08505.1 |
| FUN_003080-T1 | 0.0014977 | Inf | TBU08279.1 |
| FUN_003081-T1 | 0.00562832 | 0.36062827 | TBU08280.1 |
| FUN_003112-T1 | 0.00459185 | 0.07053914 | TBU01322.1 |
| FUN_003113-T1 | 0.00248811 | 0.16860822 | TBU01231.1 |
| FUN_003114-T1 | 0.00289283 | 0.19129976 | TBU02405.1 |
| FUN_003116-T1 | 0.00654225 | 0.22159827 | TBU06956.1 |
| FUN_003117-T1 | 0.00697236 | 0.61135268 | TBU06955.1 |
| FUN_003118-T1 | 0.004693 | 0.13525731 | TBU06954.1 |
| FUN_003133-T1 | 0.00185537 | 0.02575299 | TBU05483.1 |
| FUN_003140-T1 | 0.00588112 | 0.3574157 | TBU09485.1 |
| FUN_003141-T1 | 0.00203555 | 0.21673828 | TBU09486.1 |
| FUN_003142-T1 | 0.00299605 | 0.05002814 | TBU09487.1 |
| FUN_003147-T1 | 0.00176546 | 0 | TBT99809.1 |
| FUN_003148-T1 | 0.01258496 | 0.5969176 | TBT99810.1 |
| FUN_003149-T1 | 0.00525143 | 0.28826586 | TBU02758.1 |
| FUN_003150-T1 | 0.00371985 | 0.21943574 | TBU05502.1 |
| FUN_003151-T1 | 0.00594792 | 0.61847505 | TBU05503.1 |
| FUN_003153-T1 | 0 | NA | TBU06295.1 |
| FUN_003154-T1 | 0.00182581 | 0 | TBU06294.1 |
| FUN_003155-T1 | 0.00890041 | 0.4057844 | TBU06292.1 |
| FUN_003157-T1 | 0.00522516 | 0.07437238 | TBU04416.1 |
| FUN_003158-T1 | 0.0019947 | Inf | TBU04415.1 |
| FUN_003161-T1 | 0.00507627 | 0.20137952 | TBU03262.1 |
| FUN_003166-T1 | 0.0040223 | 0.28719904 | TBU01585.1 |
| FUN_003167-T1 | 0.00318005 | 0.14021658 | TBU01586.1 |
| FUN_003168-T1 | 0.0057433 | 0.19236505 | TBU01587.1 |
| FUN_003170-T1 | 0.00388465 | 0.06077852 | TBU03787.1 |
| FUN_003171-T1 | 0.00449595 | 0.28916346 | TBU03788.1 |
| FUN_003174-T1 | 0.00542053 | 0.15766488 | TBU05316.1 |
| FUN_003175-T1 | 0.0030396 | 0.06264143 | TBU05878.1 |
| FUN_003176-T1 | 0.00763 | 0.62664474 | TBU05877.1 |
| FUN_003177-T1 | 0.00725157 | 0.83956896 | TBU06387.1 |
| FUN_003186-T1 | 0.00183812 | 0.27655244 | TBU05315.1 |
| FUN_003187-T1 | 0.00687368 | 0.25996024 | TBU05879.1 |
| FUN_003189-T1 | 0.0030583 | 0.06794869 | TBU00605.1 |
| FUN_003190-T1 | 0.03530029 | 0.74202889 | TBU04390.1 |
| FUN_003192-T1 | 0.00321345 | 0.43603986 | TBU04391.1 |
| FUN_003194-T1 | 0.00581531 | 0.24155479 | TBU02331.1 |
| FUN_003196-T1 | 0.00745142 | 0.16491527 | TBU08130.1 |
| FUN_003197-T1 | 0.0007965 | 0 | TBU08129.1 |
| FUN_003198-T1 | 0.00563432 | 0.18236889 | TBU08128.1 |
| FUN_003199-T1 | 0.00133664 | 0.02584068 | TBU08127.1 |
| FUN_003205-T1 | 0.00276567 | 0.01978983 | TBU03858.1 |
| FUN_003206-T1 | 0.00278718 | 0.14619894 | TBU03859.1 |
| FUN_003207-T1 | 0.00394853 | 0.64461234 | TBT98303.1 |
| FUN_003212-T1 | 0.00185252 | 0 | TBU00449.1 |
| FUN_003213-T1 | 0.00138132 | 0 | TBU00448.1 |
| FUN_003215-T1 | 0.00291795 | 0.47152077 | TBU05931.1 |
| FUN_003218-T1 | 0.01093457 | 0.27419032 | TBT96811.1 |
| FUN_003219-T1 | 0.01138234 | 1.0835635 | TBT99593.1 |
| FUN_003221-T1 | 0.00885398 | 0.29543421 | TBU03882.1 |
| FUN_003222-T1 | 0.01018511 | 0.29468203 | TBU06591.1 |
| FUN_003223-T1 | 0.00488759 | 0.41114658 | TBU08515.1 |
| FUN_003224-T1 | 0.01303152 | 0.45423318 | TBU08514.1 |
| FUN_003228-T1 | 0.0035305 | 0.30343059 | TBU08513.1 |
| FUN_003229-T1 | 0.00444371 | 0.76793013 | TBU08512.1 |
| FUN_003233-T1 | 0.0040286 | 0.19633575 | TBU01234.1 |
| FUN_003234-T1 | 0.00915713 | 0.38763004 | TBU02875.1 |
| FUN_003235-T1 | 0.00797617 | 0.16101396 | TBU02876.1 |
| FUN_003236-T1 | 0.00682064 | 0.15602143 | TBU02877.1 |
| FUN_003237-T1 | 0.00257261 | 0.33304012 | TBU01167.1 |
| FUN_003238-T1 | 0.00537027 | 0.51230429 | TBU01168.1 |
| FUN_003239-T1 | 0.0026901 | 1.49095443 | TBT96899.1 |
| FUN_003240-T1 | 0.00253058 | 0.06408587 | TBU09100.1 |
| FUN_003243-T1 | 0.00850743 | 0.50920934 | TBU09101.1 |
| FUN_003247-T1 | 0.03257146 | 0.72728904 | TBU09102.1 |
| FUN_003248-T1 | 0.00337888 | 0.50624413 | TBU09103.1 |
| FUN_003250-T1 | 0.00445832 | 0.07769257 | TBU09104.1 |
| FUN_003251-T1 | 0.00211728 | 0.06878169 | TBU09105.1 |
| FUN_003252-T1 | 0.02589069 | 0.53060443 | TBT97978.1 |
| FUN_003254-T1 | 0.00386331 | 0.61811685 | TBU00361.1 |
| FUN_003257-T1 | 0.01624468 | 0.43659481 | TBU00683.1 |
| FUN_003259-T1 | 0.00251956 | 0.16071359 | TBU02506.1 |
| FUN_003260-T1 | 0.00152184 | 0.25007609 | TBU02813.1 |
| FUN_003261-T1 | 0.00171499 | 0 | TBU02812.1 |
| FUN_003262-T1 | 0.00212601 | 0.77583078 | TBU02811.1 |
| FUN_003264-T1 | 0.00879429 | 0.11536847 | TBU09025.1 |
| FUN_003265-T1 | 0.00286228 | 0.39851171 | TBU09024.1 |
| FUN_003266-T1 | 0.00137517 | 0.10719428 | TBU09023.1 |
| FUN_003267-T1 | 0.00198315 | 0.03353238 | TBU09022.1 |
| FUN_003268-T1 | 0.00315532 | 0.36032078 | TBU09021.1 |
| FUN_003275-T1 | 0.00889957 | 0.20098998 | TBU03422.1 |
| FUN_003280-T1 | 0.00676599 | 0.06616649 | TBU03159.1 |
| FUN_003282-T1 | 0.00854333 | 0.78503422 | TBT99966.1 |
| FUN_003299-T1 | 0.00449118 | 0.72144614 | TBT99568.1 |
| FUN_003300-T1 | 0.00594527 | 0.13754949 | TBU08437.1 |
| FUN_003302-T1 | 0.00120612 | Inf | TBU08438.1 |
| FUN_003308-T1 | 0.0037858 | 1.7276159 | TBU01690.1 |
| FUN_003309-T1 | 0.00045756 | 1.00320464 | TBU01688.1 |
| FUN_003310-T1 | 0.00277901 | 0.27113276 | TBU00083.1 |
| FUN_003316-T1 | 0.00157124 | 0.2437728 | TBU09326.1 |
| FUN_003323-T1 | 0.0072823 | 0.50615861 | TBU06603.1 |
| FUN_003324-T1 | 0.01388417 | 0.53239329 | TBU05181.1 |
| FUN_003326-T1 | 0.00519526 | 0.07304032 | TBU05182.1 |
| FUN_003333-T1 | 0.00544738 | 0.09088833 | TBU08808.1 |
| FUN_003334-T1 | 0.00297995 | 0.04653212 | TBU08807.1 |
| FUN_003335-T1 | 0.00901771 | 0.49545577 | TBU08806.1 |
| FUN_003336-T1 | 0.01872667 | 0.5631471 | TBU08805.1 |
| FUN_003340-T1 | 0.00200717 | 0.16799316 | TBU07825.1 |
| FUN_003342-T1 | 0.00127838 | 0 | TBU07826.1 |
| FUN_003343-T1 | 0.00580187 | 0.09871031 | TBU07827.1 |
| FUN_003344-T1 | 0.01690358 | 0.38372862 | TBU06205.1 |
| FUN_003350-T1 | 0.01598931 | 0.72301676 | TBU08136.1 |
| FUN_003351-T1 | 0.0158575 | 1.06476516 | TBU08137.1 |
| FUN_003354-T1 | 0.00265476 | 0.1971794 | TBU08235.1 |
| FUN_003355-T1 | 0.00480246 | 0.07437653 | TBU08236.1 |
| FUN_003357-T1 | 0.00428882 | 0.15742483 | TBU08237.1 |
| FUN_003358-T1 | 0.00627001 | Inf | TBU08238.1 |
| FUN_003360-T1 | 0.00615197 | 0.34699802 | TBU08239.1 |
| FUN_003361-T1 | 0.01171848 | 1.56895944 | TBU09617.1 |
| FUN_003363-T1 | 0.00728164 | 0.37767653 | TBU09618.1 |
| FUN_003370-T1 | 0.01351387 | 0.26629473 | TBU09621.1 |
| FUN_003371-T1 | 0.01326201 | 0.608396 | TBU09622.1 |
| FUN_003372-T1 | 0.00985019 | 0.99818858 | TBT99527.1 |
| FUN_003373-T1 | 0.00215859 | 0.08287282 | TBU02316.1 |
| FUN_003375-T1 | 0.00805033 | 0.56784359 | TBU04630.1 |
| FUN_003376-T1 | 0.00426815 | 0.49527564 | TBU04631.1 |
| FUN_003377-T1 | 0.00227957 | 0.23466139 | TBU05463.1 |
| FUN_003378-T1 | 0.00238704 | 0.12482281 | TBU05462.1 |
| FUN_003379-T1 | 0.00391924 | 1.01157668 | TBU05461.1 |
| FUN_003381-T1 | 0.00180513 | 0.12421719 | TBU05460.1 |
| FUN_003382-T1 | 0.00145483 | 0.13033732 | TBU05760.1 |
| FUN_003383-T1 | 0.00849679 | 0.30452185 | TBU05761.1 |
| FUN_003386-T1 | 0.00657566 | 0.46256333 | TBU02652.1 |
| FUN_003387-T1 | 0.00372181 | 0.16586845 | TBU02653.1 |
| FUN_003388-T1 | 0.00335119 | 0.33278499 | TBU05244.1 |
| FUN_003389-T1 | 0.00171779 | 0.23798561 | TBU05245.1 |
| FUN_003390-T1 | 0.00097713 | 0.05327852 | TBU05246.1 |
| FUN_003391-T1 | 0.00224974 | 0.09424274 | TBU05247.1 |
| FUN_003392-T1 | 0.0027402 | 6.79579639 | TBU02061.1 |
| FUN_003394-T1 | 0.00719713 | 0.94839584 | TBU07390.1 |
| FUN_003395-T1 | 0.00251982 | Inf | TBU07389.1 |
| FUN_003396-T1 | 0.00026226 | 0 | TBU07388.1 |
| FUN_003397-T1 | 0.00729496 | 0.40196893 | TBU07387.1 |
| FUN_003398-T1 | 0.00734237 | 0.32876829 | TBU01417.1 |
| FUN_003399-T1 | 0.0154187 | 0.95182131 | TBU01418.1 |
| FUN_003401-T1 | 0.00348865 | 0.45598131 | TBU02254.1 |
| FUN_003402-T1 | 0.00433158 | 0.43683248 | TBT97844.1 |
| FUN_003405-T1 | 0.00257616 | 0.07496897 | TBU00963.1 |
| FUN_003407-T1 | 0.0020962 | 0.16449198 | TBU06643.1 |
| FUN_003408-T1 | 0.00397445 | 0.30735427 | TBU06644.1 |
| FUN_003409-T1 | 0.00414704 | 0.0253882 | TBU06645.1 |
| FUN_003410-T1 | 0.00753213 | 0.41045879 | TBU00245.1 |
| FUN_003411-T1 | 0.00236871 | 0.609921 | TBT97883.1 |
| FUN_003416-T1 | 0.0054741 | 0.12303933 | TBU01631.1 |
| FUN_003418-T1 | 0.00465879 | 0.46257955 | TBU01930.1 |
| FUN_003420-T1 | 0.01077957 | 0.2185996 | TBT98200.1 |
| FUN_003422-T1 | 0.00757213 | 0.90948985 | TBU01580.1 |
| FUN_003428-T1 | 0.00798771 | 0.12143097 | TBT97555.1 |
| FUN_003429-T1 | 0.0043329 | 0.09978262 | TBU08990.1 |
| FUN_003430-T1 | 0.00386141 | 0.03342554 | TBU08991.1 |
| FUN_003431-T1 | 0.00291308 | 0.064527 | TBU08992.1 |
| FUN_003432-T1 | 0.00081925 | 0.33141953 | TBU08993.1 |
| FUN_003433-T1 | 0.00715588 | 0.08003923 | TBU08994.1 |
| FUN_003434-T1 | 0.00496286 | 0.16496095 | TBU08995.1 |
| FUN_003435-T1 | 0.00358423 | 0.01216799 | TBU06626.1 |
| FUN_003437-T1 | 0.01313849 | 2.42240463 | TBU01354.1 |
| FUN_003444-T1 | 0.00496534 | 0.28339132 | TBU06411.1 |
| FUN_003445-T1 | 0.00736356 | 0.34220124 | TBU05804.1 |
| FUN_003447-T1 | 0.01077206 | 0.58415715 | TBU05803.1 |
| FUN_003448-T1 | 0.00216734 | 0.2209279 | TBU05802.1 |
| FUN_003449-T1 | 0.00542755 | 0.16085712 | TBU00611.1 |
| FUN_003451-T1 | 0.02197235 | 1.01664783 | TBU02485.1 |
| FUN_003452-T1 | 0.00328676 | 0.16049299 | TBU02909.1 |
| FUN_003453-T1 | 0.00453348 | 0.1279027 | TBU02908.1 |
| FUN_003456-T1 | 0.00010889 | 0 | TBU03430.1 |
| FUN_003457-T1 | 0.00227814 | 10.9919219 | TBU02885.1 |
| FUN_003458-T1 | 0.00061649 | 0.52025685 | TBU06153.1 |
| FUN_003459-T1 | 0.0021525 | 0.387395 | TBU06152.1 |
| FUN_003461-T1 | 0.00773218 | 0.5776407 | TBU06843.1 |
| FUN_003463-T1 | 0.00039469 | 0 | TBU08265.1 |
| FUN_003465-T1 | 0.01026351 | 1.18493347 | TBU05827.1 |
| FUN_003466-T1 | 0.00228402 | 0.07562625 | TBU05828.1 |
| FUN_003467-T1 | 0.00291837 | 0.34312074 | TBU05829.1 |
| FUN_003468-T1 | 0.00443221 | 0.3790226 | TBU00712.1 |
| FUN_003471-T1 | 0.00647752 | 0.91985057 | TBU04112.1 |
| FUN_003476-T1 | 0.00073314 | 0.31827487 | TBU00729.1 |
| FUN_003477-T1 | 0.00273237 | 0.28865519 | TBU00730.1 |
| FUN_003478-T1 | 0.001666 | 0.03427945 | TBU03061.1 |
| FUN_003479-T1 | 0.00649776 | 0.269706 | TBU08173.1 |
| FUN_003480-T1 | 0.00598597 | 0.21170781 | TBU08172.1 |
| FUN_003482-T1 | 0.00076205 | 0 | TBU08171.1 |
| FUN_003483-T1 | 0.00326814 | 0.56910311 | TBU06668.1 |
| FUN_003484-T1 | 0.00296388 | 0.6238384 | TBU06667.1 |
| FUN_003487-T1 | 0.00152355 | 0.02227022 | TBU08255.1 |
| FUN_003490-T1 | 0.00299173 | 0.07223248 | TBU08254.1 |
| FUN_003491-T1 | 0.00172915 | 0.10571915 | TBU08253.1 |
| FUN_003492-T1 | 0.0054081 | 0.32822574 | TBU08252.1 |
| FUN_003493-T1 | 0.00460255 | 0.04050794 | TBT96867.1 |
| FUN_003494-T1 | 0.00019088 | 0.1243677 | TBU08731.1 |
| FUN_003495-T1 | 0.00211937 | 0.02833572 | TBU08730.1 |
| FUN_003496-T1 | 0.00147465 | 0 | TBU08729.1 |
| FUN_003497-T1 | 0.00516539 | 1.17827738 | TBU08727.1 |
| FUN_003498-T1 | 0.00737708 | 0.4812731 | TBU08726.1 |
| FUN_003499-T1 | 0.01128585 | 0.50597692 | TBU08725.1 |
| FUN_003500-T1 | 0.00471762 | 0.32711068 | TBU01481.1 |
| FUN_003502-T1 | 0.00313812 | 0.79202778 | TBU05906.1 |
| FUN_003503-T1 | 0.00875835 | 0.28615982 | TBU05618.1 |
| FUN_003504-T1 | 0.01247537 | 1.45132016 | TBT98424.1 |
| FUN_003505-T1 | 0.00442538 | 0.17308622 | TBU04884.1 |
| FUN_003513-T1 | 0.00668227 | 0.45876523 | TBU07802.1 |
| FUN_003515-T1 | 0.00707639 | 1.03366544 | TBU07803.1 |
| FUN_003516-T1 | 0.00494645 | 0.14610457 | TBU07805.1 |
| FUN_003517-T1 | 0.00173293 | 0.16816097 | TBU00496.1 |
| FUN_003518-T1 | 0.00257312 | 1.30926306 | TBU01873.1 |
| FUN_003519-T1 | 0.01923902 | 1.09235518 | TBU08895.1 |
| FUN_003520-T1 | 0.00187386 | 0.44603701 | TBU08894.1 |
| FUN_003521-T1 | 0.0179599 | 0.68971234 | TBU08893.1 |
| FUN_003522-T1 | 0.00318026 | 0.60348375 | TBU08892.1 |
| FUN_003527-T1 | 0.00135355 | 0.19397917 | TBU08546.1 |
| FUN_003530-T1 | 0.003181 | 0.20828567 | TBU08545.1 |
| FUN_003531-T1 | 0.0027402 | 0 | TBU08544.1 |
| FUN_003532-T1 | 0.0123563 | 0.56124783 | TBU08543.1 |
| FUN_003534-T1 | 0.00477087 | 0.38333863 | TBU02099.1 |
| FUN_003535-T1 | 0.00403774 | 1.98740458 | TBU02104.1 |
| FUN_003536-T1 | 0.00119785 | 5.97114991 | TBU02103.1 |
| FUN_003538-T1 | 0.00114695 | 0.82324219 | TBU02004.1 |
| FUN_003540-T1 | 0.00471848 | 0.40978572 | TBU07710.1 |
| FUN_003554-T1 | 0 | NA | TBT99158.1 |
| FUN_003560-T1 | 0.00462787 | 1.43684252 | TBT99985.1 |
| FUN_003573-T1 | 0.00545876 | 0.05518088 | TBU02608.1 |
| FUN_003574-T1 | 0.00127498 | 0 | TBU02609.1 |
| FUN_003575-T1 | 0.00252814 | 0.15898803 | TBT98064.1 |
| FUN_003577-T1 | 0.00304489 | 0.09754206 | TBU04047.1 |
| FUN_003584-T1 | 0.01014663 | 0.28990059 | TBU04861.1 |
| FUN_003605-T1 | 0.00824373 | 0.35614481 | TBU07711.1 |
| FUN_003613-T1 | 0.00191403 | 0.5805561 | TBU06674.1 |
| FUN_003615-T1 | 0.00173281 | 0.40597796 | TBU06673.1 |
| FUN_003616-T1 | 0.00479098 | 1.25147534 | TBU06672.1 |
| FUN_003617-T1 | 0.00274665 | 0.05122081 | TBT97628.1 |
| FUN_003618-T1 | 0.00307046 | 0.08111673 | TBU03586.1 |
| FUN_003619-T1 | 0.00577214 | 0.28458762 | TBU03585.1 |
| FUN_003620-T1 | 0.00403337 | 0.11260078 | TBU03584.1 |
| FUN_003621-T1 | 0.00355322 | 0.01525783 | TBU03583.1 |
| FUN_003623-T1 | 0.00248221 | 0 | TBU08735.1 |
| FUN_003624-T1 | 0.00230936 | Inf | TBU08736.1 |
| FUN_003625-T1 | 0.00665233 | 2.52911102 | TBU08737.1 |
| FUN_003626-T1 | 0.00199543 | 0.25221731 | TBU08738.1 |
| FUN_003627-T1 | 0.00130753 | 0 | TBU08739.1 |
| FUN_003628-T1 | 0.00193413 | 0.05915953 | TBU08740.1 |
| FUN_003630-T1 | 0.00034136 | 0 | TBT98607.1 |
| FUN_003631-T1 | 0.00159334 | 0.02944004 | TBU05628.1 |
| FUN_003633-T1 | 0.00186874 | Inf | TBU05629.1 |
| FUN_003637-T1 | 0.00250327 | 0.13367337 | TBU02050.1 |
| FUN_003638-T1 | 0.00172043 | 0 | TBU02049.1 |
| FUN_003640-T1 | 0.00515599 | 0.20244255 | TBU07685.1 |
| FUN_003641-T1 | 0.0032607 | 0.12972113 | TBU07684.1 |
| FUN_003645-T1 | 0.00111509 | 0 | TBU06882.1 |
| FUN_003646-T1 | 0.00238004 | 0.24057962 | TBU06881.1 |
| FUN_003647-T1 | 0.00238557 | 0 | TBU06880.1 |
| FUN_003648-T1 | 0.00352969 | 0.13628189 | TBU06879.1 |
| FUN_003651-T1 | 0.00278803 | 0.06844719 | TBU00929.1 |
| FUN_003652-T1 | 0.00181719 | 0.76443334 | TBU00380.1 |
| FUN_003653-T1 | 0.00373955 | 0.16631545 | TBU08383.1 |
| FUN_003656-T1 | 0.02084367 | 0.20666476 | TBU08384.1 |
| FUN_003657-T1 | 0.00099256 | 1.27018163 | TBU08385.1 |
| FUN_003658-T1 | 0.00418261 | 0.50190836 | TBU03660.1 |
| FUN_003659-T1 | 0.00265885 | 4.11972839 | TBU03659.1 |
| FUN_003660-T1 | 0 | NA | TBT97061.1 |
| FUN_003661-T1 | 0.00256761 | 0.06872813 | TBT98313.1 |
| FUN_003662-T1 | 0.00224277 | 0.14243742 | TBU08886.1 |
| FUN_003663-T1 | 0.00045019 | Inf | TBU08885.1 |
| FUN_003664-T1 | 0.00267042 | 1.72678551 | TBT97585.1 |
| FUN_003665-T1 | 0.00301944 | 5.88396583 | TBT99630.1 |
| FUN_003690-T1 | 0.00495648 | 0.45927095 | TBU06043.1 |
| FUN_003694-T1 | 0.00363937 | 0.58554119 | TBU09003.1 |
| FUN_003695-T1 | 0.00032634 | 1.35703427 | TBU09004.1 |
| FUN_003696-T1 | 0.01458311 | 0.38464261 | TBU09005.1 |
| FUN_003716-T1 | 0.00587097 | 0 | TBU08557.1 |
| FUN_003717-T1 | 0.00704012 | 0.23110149 | TBU08558.1 |
| FUN_003718-T1 | 0.00716846 | 0.2471853 | TBU07240.1 |
| FUN_003720-T1 | 0.00013698 | Inf | TBU07241.1 |
| FUN_003724-T1 | 0.00020481 | 0 | TBU08630.1 |
| FUN_003725-T1 | 0.00439591 | 0.3874329 | TBU08629.1 |
| FUN_003726-T1 | 0.00457506 | 0.14123879 | TBU08628.1 |
| FUN_003730-T1 | 0.01405995 | 1.05314176 | TBU00854.1 |
| FUN_003732-T1 | 0.00660623 | 0.23567179 | TBU03347.1 |
| FUN_003737-T1 | 0.00545401 | 0.61472032 | TBU08961.1 |
| FUN_003738-T1 | 0.0196506 | 2.00199019 | TBU01826.1 |
| FUN_003739-T1 | 0.0043479 | 0.13612144 | TBU06110.1 |
| FUN_003746-T1 | 0 | NA | TBU00672.1 |
| FUN_003747-T1 | 0.01604788 | 0.70856381 | TBU06017.1 |
| FUN_003748-T1 | 0.00794571 | 0.26958386 | TBU06018.1 |
| FUN_003749-T1 | 0.00041758 | Inf | TBU06020.1 |
| FUN_003750-T1 | 0.00342761 | 0.26235854 | TBT98305.1 |
| FUN_003752-T1 | 0.00246992 | 0.15819492 | TBU03759.1 |
| FUN_003755-T1 | 0.00570974 | 0.16983961 | TBT98108.1 |
| FUN_003756-T1 | 0.0024335 | 0.00948518 | TBU03404.1 |
| FUN_003757-T1 | 0.00270248 | 0.34653983 | TBU03405.1 |
| FUN_003759-T1 | 0.04567332 | 0.69778046 | TBU03665.1 |
| FUN_003761-T1 | 0.00352823 | 0.36275943 | TBU06139.1 |
| FUN_003767-T1 | 0.00262971 | 1.15593154 | TBU06663.1 |
| FUN_003771-T1 | 0.00261756 | 0.17376168 | TBU09337.1 |
| FUN_003772-T1 | 0.00616922 | 0.08123611 | TBU09336.1 |
| FUN_003773-T1 | 0.00079037 | 0.75137087 | TBU09335.1 |
| FUN_003774-T1 | 0.00529783 | 0.02771429 | TBU09334.1 |
| FUN_003775-T1 | 0.00471334 | 0.74538787 | TBU09333.1 |
| FUN_003776-T1 | 0.00471802 | 0.2279066 | TBU09332.1 |
| FUN_003777-T1 | 0.01422954 | 0.58294851 | TBT99844.1 |
| FUN_003780-T1 | 0.00367641 | 0.44099911 | TBU07926.1 |
| FUN_003782-T1 | 0.00189247 | Inf | TBU07923.1 |
| FUN_003790-T1 | 0.0001044 | 0 | TBT99979.1 |
| FUN_003791-T1 | 0.00070946 | 0 | TBT97391.1 |
| FUN_003792-T1 | 0.00024701 | 0.10732554 | TBU09056.1 |
| FUN_003793-T1 | 0.01049706 | 1.7099628 | TBU09055.1 |
| FUN_003794-T1 | 0.00530928 | 0.3498603 | TBU09053.1 |
| FUN_003804-T1 | 0.00213044 | 0.32 | TBU09050.1 |
| FUN_003808-T1 | 0.0116824 | 0.3893855 | TBU09054.1 |
| FUN_003809-T1 | 0.00042501 | 0.07961783 | TBU05832.1 |
| FUN_003811-T1 | 0.00155054 | 0.48887015 | TBU01532.1 |
| FUN_003815-T1 | 0.0042008 | 1.04828257 | TBT98947.1 |
| FUN_003816-T1 | 0.00243611 | 0.06182095 | TBT99769.1 |
| FUN_003817-T1 | 0.02258343 | 0.70342158 | TBU01527.1 |
| FUN_003818-T1 | 0.00390867 | 0.42009668 | TBT99925.1 |
| FUN_003820-T1 | 0.00631507 | 0.31903217 | TBT99080.1 |
| FUN_003821-T1 | 0.00946648 | 0.8719735 | TBT99029.1 |
| FUN_003822-T1 | 0.0341556 | 1.02033548 | TBU03382.1 |
| FUN_003823-T1 | 0.00661603 | 0.13507067 | TBU03383.1 |
| FUN_003824-T1 | 0.00549311 | 0.01941419 | TBU03384.1 |
| FUN_003826-T1 | 0.03551856 | 0.52379194 | TBU03683.1 |
| FUN_003827-T1 | 0.01028483 | 0.91336481 | TBU03682.1 |
| FUN_003828-T1 | 0.00601395 | 0.16697282 | TBT99178.1 |
| FUN_003830-T1 | 0.00650061 | 0.14260739 | TBU06240.1 |
| FUN_003833-T1 | 0.00778565 | 0.13503157 | TBU07001.1 |
| FUN_003838-T1 | 0.01592215 | 0.53336286 | TBU04575.1 |
| FUN_003839-T1 | 0.01206171 | 0.94025608 | TBU07838.1 |
| FUN_003840-T1 | 0.02303843 | 0.71724497 | TBU07839.1 |
| FUN_003841-T1 | 0.00757844 | 0.45821223 | TBU07840.1 |
| FUN_003844-T1 | 0.00309291 | 0.18368834 | TBU06241.1 |
| FUN_003846-T1 | 0.00800478 | 0.07783509 | TBU06621.1 |
| FUN_003858-T1 | 0.00736328 | 0.51432672 | TBU03454.1 |
| FUN_003859-T1 | 0.00577932 | 0.0663258 | TBU08612.1 |
| FUN_003860-T1 | 0.00962573 | 0.37224476 | TBU08611.1 |
| FUN_003862-T1 | 0.00970736 | 0.41502106 | TBU08610.1 |
| FUN_003865-T1 | 0.02493166 | 1.35793249 | TBU02311.1 |
| FUN_003866-T1 | 0.00687247 | 4.14179202 | TBT99154.1 |
| FUN_003869-T1 | 0.00617232 | 0.25372558 | TBU01753.1 |
| FUN_003870-T1 | 0 | NA | TBU01754.1 |
| FUN_003871-T1 | 0.01601207 | 1.05095066 | TBU06185.1 |
| FUN_003887-T1 | 0.00086531 | 0 | TBU04299.1 |
| FUN_003888-T1 | 0.00471553 | 1.0107731 | TBU04300.1 |
| FUN_003889-T1 | 0.00513548 | 0.16973189 | TBU08101.1 |
| FUN_003890-T1 | 0.00736376 | 0.85703844 | TBU08102.1 |
| FUN_003900-T1 | 0.00397993 | 0.32083601 | TBU09164.1 |
| FUN_003901-T1 | 0.0009507 | 0.60953488 | TBU09163.1 |
| FUN_003903-T1 | 0.00180658 | 0.20556936 | TBU09162.1 |
| FUN_003904-T1 | 0.03051462 | 1.3082452 | TBU09161.1 |
| FUN_003905-T1 | 0 | NA | TBT99765.1 |
| FUN_003906-T1 | 0.0017979 | 0.37030458 | TBU09709.1 |
| FUN_003911-T1 | 0.03027295 | 0.66586332 | TBU00598.1 |
| FUN_003912-T1 | 0.00121754 | 0 | TBU00597.1 |
| FUN_003914-T1 | 0.00385731 | 0 | TBU00596.1 |
| FUN_003936-T1 | 0.0012874 | 0.141928 | TBU07950.1 |
| FUN_003937-T1 | 0.00674596 | 0.29697023 | TBU07949.1 |
| FUN_003938-T1 | 0.02534261 | 0.28650255 | TBU07948.1 |
| FUN_003942-T1 | 0.0024797 | 0 | TBU03952.1 |
| FUN_003943-T1 | 0.0009146 | 0.02680828 | TBU03953.1 |
| FUN_003944-T1 | 0.00025352 | 0 | TBU03954.1 |
| FUN_003948-T1 | 9.07E-05 | Inf | TBU00268.1 |
| FUN_003949-T1 | 0.01472205 | 0.43894187 | TBU06632.1 |
| FUN_003951-T1 | 0.00317618 | 0.58126356 | TBU06631.1 |
| FUN_003954-T1 | 0.00163712 | 0.08534681 | TBT98754.1 |
| FUN_003955-T1 | 0.0011411 | 0 | TBU08826.1 |
| FUN_003957-T1 | 0.0013974 | 0.12558245 | TBU08827.1 |
| FUN_003958-T1 | 0.00103178 | 0.16612436 | TBU08828.1 |
| FUN_003959-T1 | 0 | NA | TBU08829.1 |
| FUN_003964-T1 | 0.00480397 | 0.27581457 | TBU06549.1 |
| FUN_003967-T1 | 0.00399015 | 0.24088734 | TBT99247.1 |
| FUN_003970-T1 | 0.00493584 | 0.16870573 | TBU06696.1 |
| FUN_003971-T1 | 0.02604496 | 2.22147017 | TBU00416.1 |
| FUN_003972-T1 | 0.02246634 | 0.47943842 | TBT98234.1 |
| FUN_003973-T1 | 0.00087777 | Inf | TBU02370.1 |
| FUN_003974-T1 | 0.00256272 | 0.02827855 | TBT97021.1 |
| FUN_003975-T1 | 0.00531783 | 1.63929498 | TBU05652.1 |
| FUN_003976-T1 | 0.00324849 | 0.08872627 | TBU05653.1 |
| FUN_003977-T1 | 0.00386774 | 0.54550336 | TBU05654.1 |
| FUN_003980-T1 | 0 | NA | TBT96841.1 |
| FUN_003985-T1 | 0.00304384 | 0.15216228 | TBT99627.1 |
| FUN_003988-T1 | 0.00353152 | 0.08877694 | TBU02676.1 |
| FUN_003991-T1 | 0.00987095 | 0.27360486 | TBU05038.1 |
| FUN_003993-T1 | 0.00579501 | 0.48416693 | TBU04901.1 |
| FUN_003994-T1 | 0.00918262 | 0.86680453 | TBT99460.1 |
| FUN_003995-T1 | 0.01828776 | 0.93543439 | TBU01096.1 |
| FUN_003996-T1 | 0.00346932 | 0.14741431 | TBT97133.1 |
| FUN_003997-T1 | 0.00456493 | 2.67392679 | TBU07460.1 |
| FUN_003998-T1 | 0.0021362 | 0.05312609 | TBU07461.1 |
| FUN_003999-T1 | 0.00387402 | 0.13087831 | TBU07462.1 |
| FUN_004000-T1 | 0.0048849 | 0.54390735 | TBU07463.1 |
| FUN_004003-T1 | 0.00506571 | 0.24955134 | TBU07464.1 |
| FUN_004004-T1 | 0.0046266 | 0.48040081 | TBU07465.1 |
| FUN_004032-T1 | 0.00370002 | 0.16804131 | TBU02578.1 |
| FUN_004039-T1 | 0.00284336 | 0.1383547 | TBU07855.1 |
| FUN_004044-T1 | 0.00340925 | 0.13191097 | TBU00288.1 |
| FUN_004045-T1 | 0.00693744 | 0.10523187 | TBU00289.1 |
| FUN_004048-T1 | 0.00665643 | 0.21322245 | TBU08370.1 |
| FUN_004050-T1 | 0.00175229 | 0 | TBU08371.1 |
| FUN_004054-T1 | 0.00310701 | 0.12062953 | TBU01225.1 |
| FUN_004061-T1 | 0.00581313 | 0.89588651 | TBU03124.1 |
| FUN_004066-T1 | 0.00552456 | 0.22230021 | TBU07357.1 |
| FUN_004067-T1 | 0.00443221 | 0.06642719 | TBU07356.1 |
| FUN_004068-T1 | 0.00149221 | 0.09487168 | TBU07355.1 |
| FUN_004070-T1 | 0.00074671 | 0 | TBU07354.1 |
| FUN_004071-T1 | 0.01232314 | 0.35150947 | TBU07353.1 |
| FUN_004072-T1 | 0.00486209 | 0.10258641 | TBU07352.1 |
| FUN_004074-T1 | 0.00576652 | 0.47440318 | TBU05574.1 |
| FUN_004075-T1 | 0.00320846 | 0.01929892 | TBU05573.1 |
| FUN_004076-T1 | 0.00294663 | 0.1014064 | TBU08306.1 |
| FUN_004077-T1 | 0.00179874 | 1.17220049 | TBU08305.1 |
| FUN_004078-T1 | 0.00184871 | 0.07614302 | TBU08304.1 |
| FUN_004079-T1 | 0.00318896 | 0.20979899 | TBU08303.1 |
| FUN_004080-T1 | 0.00139844 | 0 | TBU08302.1 |
| FUN_004082-T1 | 0.00403669 | 0.18739152 | TBU08301.1 |
| FUN_004083-T1 | 0.00096248 | 0.28507834 | TBU08300.1 |
| FUN_004085-T1 | 0.00378964 | 0.08017044 | TBU08433.1 |
| FUN_004093-T1 | 0.00708963 | 0.96960928 | TBU08429.1 |
| FUN_004094-T1 | 0.00214489 | 0.11734195 | TBT98612.1 |
| FUN_004096-T1 | 0.00669472 | 0.22257879 | TBT97760.1 |
| FUN_004097-T1 | 0.02977667 | 0.63657757 | TBT98437.1 |
| FUN_004098-T1 | 0.00275653 | 0.61431257 | TBU06398.1 |
| FUN_004099-T1 | 0.00807115 | 0.43031518 | TBU06397.1 |
| FUN_004101-T1 | 0.00092768 | Inf | TBU06396.1 |
| FUN_004103-T1 | 0.00127796 | Inf | TBU09598.1 |
| FUN_004104-T1 | 0.00364326 | 0.28997972 | TBU09597.1 |
| FUN_004105-T1 | 0.00447836 | 0.15108005 | TBU09596.1 |
| FUN_004106-T1 | 0.00056593 | Inf | TBU09595.1 |
| FUN_004107-T1 | 0.00184558 | 0 | TBU09594.1 |
| FUN_004108-T1 | 0.00224014 | 1.38975343 | TBT97612.1 |
| FUN_004109-T1 | 0.0046053 | 0.20904539 | TBU02630.1 |
| FUN_004112-T1 | 0.00620072 | 0.45674789 | TBU02683.1 |
| FUN_004113-T1 | 0.00081019 | 0 | TBU02682.1 |
| FUN_004116-T1 | 0.0025943 | 0.01344036 | TBT99799.1 |
| FUN_004117-T1 | 0.00196745 | 1.60249769 | TBU07940.1 |
| FUN_004118-T1 | 0.00279679 | 0.0801902 | TBU07939.1 |
| FUN_004119-T1 | 0.00298024 | 0.05563987 | TBU07938.1 |
| FUN_004120-T1 | 0.00587144 | 0.36608849 | TBU01967.1 |
| FUN_004121-T1 | 0.00390613 | 0.07988576 | TBU00819.1 |
| FUN_004123-T1 | 0.00536149 | 0.11202042 | TBU00115.1 |
| FUN_004124-T1 | 0.00418404 | 0 | TBU07907.1 |
| FUN_004126-T1 | 0.01330096 | 0.14935273 | TBU07906.1 |
| FUN_004128-T1 | 0.00113613 | 0.05587788 | TBU07905.1 |
| FUN_004129-T1 | 0.00473915 | 0.32443687 | TBU07904.1 |
| FUN_004130-T1 | 0.00366189 | 0.24738593 | TBU00862.1 |
| FUN_004133-T1 | 0.00095848 | Inf | TBU04276.1 |
| FUN_004135-T1 | 0.0025177 | 0.49147436 | TBU03052.1 |
| FUN_004136-T1 | 0.00208741 | 0.05180573 | TBU03051.1 |
| FUN_004138-T1 | 0.00517937 | 0.28706547 | TBU03050.1 |
| FUN_004139-T1 | 0.00295341 | 0.23903896 | TBU03049.1 |
| FUN_004141-T1 | 0.01602519 | 0.53145983 | TBT98720.1 |
| FUN_004142-T1 | 0.02402867 | 1.1097297 | TBU07061.1 |
| FUN_004143-T1 | 0.00433946 | 0.24660319 | TBU07062.1 |
| FUN_004144-T1 | 0.0078776 | 0.17633869 | TBU07063.1 |
| FUN_004145-T1 | 0.004795 | 0.03796062 | TBU07064.1 |
| FUN_004146-T1 | 0.00590544 | 0.29647684 | TBU07065.1 |
| FUN_004148-T1 | 0.00310776 | 0.09060322 | TBU00900.1 |
| FUN_004150-T1 | 0.00272056 | 0 | TBU05050.1 |
| FUN_004151-T1 | 0.01233065 | 0.45143775 | TBU05049.1 |
| FUN_004152-T1 | 0.04008961 | 0.5430271 | TBU05048.1 |
| FUN_004153-T1 | 0.00382923 | 0.45140517 | TBU05935.1 |
| FUN_004157-T1 | 0.00467598 | 0.12043792 | TBU05578.1 |
| FUN_004158-T1 | 0.00165202 | 0.27327924 | TBU05577.1 |
| FUN_004161-T1 | 0.00566517 | 0.24463995 | TBU05576.1 |
| FUN_004162-T1 | 0.00425095 | 0.06905983 | TBU00924.1 |
| FUN_004168-T1 | 0.01210399 | 0.44633755 | TBU06838.1 |
| FUN_004169-T1 | 0.00905344 | 0.33932215 | TBU06837.1 |
| FUN_004170-T1 | 0.00277018 | 0.21690321 | TBU06836.1 |
| FUN_004171-T1 | 0.00183766 | 0.53671961 | TBU07370.1 |
| FUN_004173-T1 | 0.00172695 | 0.03833934 | TBU07369.1 |
| FUN_004174-T1 | 0.00249396 | 0.5020757 | TBU07368.1 |
| FUN_004175-T1 | 0.00719251 | 0.12870502 | TBU07367.1 |
| FUN_004179-T1 | 0.00316527 | 0 | TBT99223.1 |
| FUN_004180-T1 | 0.00617791 | 0.32209011 | TBT99224.1 |
| FUN_004181-T1 | 0.00288072 | 0.15923687 | TBU06711.1 |
| FUN_004183-T1 | 0.00296199 | 0.60136237 | TBU06712.1 |
| FUN_004185-T1 | 0.00375597 | Inf | TBU07673.1 |
| FUN_004186-T1 | 0.00301428 | 0.19557173 | TBU07674.1 |
| FUN_004187-T1 | 0.00554146 | 0.12292435 | TBU07675.1 |
| FUN_004190-T1 | 0.00277645 | 0.12473694 | TBU07677.1 |
| FUN_004191-T1 | 0.00150538 | 2.37945863 | TBU07678.1 |
| FUN_004192-T1 | 0.00152687 | 0.07697165 | TBU00503.1 |
| FUN_004194-T1 | 0.00465076 | 0.13585518 | TBU00502.1 |
| FUN_004195-T1 | 0.0039795 | 0.44285724 | TBU00501.1 |
| FUN_004197-T1 | 0.016053 | 1.4714234 | TBU02897.1 |
| FUN_004198-T1 | 0.00944138 | 1.0390214 | TBU02896.1 |
| FUN_004203-T1 | 0.00323352 | 0.16875708 | TBU06775.1 |
| FUN_004205-T1 | 0.01074361 | 0.25108866 | TBU06776.1 |
| FUN_004209-T1 | 0.00573477 | 0.48742864 | TBU00514.1 |
| FUN_004212-T1 | 0.00353608 | 0.09152303 | TBU03607.1 |
| FUN_004213-T1 | 0.00591398 | Inf | TBT96754.1 |
| FUN_004214-T1 | 0.0034906 | 0.19037292 | TBU03773.1 |
| FUN_004215-T1 | 0.00199771 | 0.32137258 | TBT99469.1 |
| FUN_004216-T1 | 0 | NA | TBT99468.1 |
| FUN_004217-T1 | 0.00443804 | 0.24135368 | TBU05220.1 |
| FUN_004218-T1 | 0.00249848 | 8.99456067 | TBU03506.1 |
| FUN_004225-T1 | 0.00107527 | 0.47139588 | TBU08579.1 |
| FUN_004227-T1 | 0.0045383 | 0.18967582 | TBU08578.1 |
| FUN_004229-T1 | 0.00089312 | 0 | TBT99736.1 |
| FUN_004230-T1 | 0.00032049 | 0 | TBT99735.1 |
| FUN_004234-T1 | 0.00738778 | 0.49614328 | TBU03914.1 |
| FUN_004236-T1 | 0.0030724 | 0.08161207 | TBU07005.1 |
| FUN_004237-T1 | 0.00516676 | 0.53573722 | TBU07006.1 |
| FUN_004239-T1 | 0.01381179 | 0.35599936 | TBU00753.1 |
| FUN_004241-T1 | 0.01075269 | 0.15368652 | TBU01370.1 |
| FUN_004243-T1 | 0.02058019 | 0.5771495 | TBT99678.1 |
| FUN_004245-T1 | 0.00483402 | 0.20315135 | TBU08412.1 |
| FUN_004246-T1 | 0.02783049 | 0.57772898 | TBU08411.1 |
| FUN_004247-T1 | 0.00596988 | 0.28796013 | TBU08410.1 |
| FUN_004250-T1 | 0.00389062 | 1.35507725 | TBU08870.1 |
| FUN_004255-T1 | 0.01946472 | 2.31245813 | TBU03737.1 |
| FUN_004257-T1 | 0.03911074 | 1.04028771 | TBU08935.1 |
| FUN_004259-T1 | 0.01899571 | 0.33804513 | TBU08933.1 |
| FUN_004260-T1 | 0.01620915 | 0.65028798 | TBU04350.1 |
| FUN_004264-T1 | 0.00726655 | 0.42318634 | TBT96937.1 |
| FUN_004265-T1 | 0.00986984 | 5.69608305 | TBT97035.1 |
| FUN_004267-T1 | 0.00527335 | 0.1345545 | TBT97955.1 |
| FUN_004268-T1 | 0.01989395 | 1.15170821 | TBU05606.1 |
| FUN_004269-T1 | 0.01186135 | 1.03461789 | TBU05605.1 |
| FUN_004270-T1 | 0.006851 | 0.25591483 | TBU05604.1 |
| FUN_004271-T1 | 0.00355106 | 0.10220425 | TBU05603.1 |
| FUN_004277-T1 | 0.00774523 | 0.29174711 | TBU05196.1 |
| FUN_004278-T1 | 0.00499308 | 0.23987906 | TBU08493.1 |
| FUN_004279-T1 | 0.00222837 | 0.01570458 | TBU08492.1 |
| FUN_004280-T1 | 0.00258167 | 0.10609128 | TBU08491.1 |
| FUN_004282-T1 | 0.0025574 | Inf | TBU08490.1 |
| FUN_004284-T1 | 0.00319757 | 0.06252623 | TBU08489.1 |
| FUN_004285-T1 | 0.01010028 | 0.63710551 | TBU06760.1 |
| FUN_004286-T1 | 0.00200894 | 0 | TBU06761.1 |
| FUN_004287-T1 | 0.00526652 | 0.08771679 | TBU06762.1 |
| FUN_004288-T1 | 0.00530853 | 0.30447749 | TBU06763.1 |
| FUN_004289-T1 | 0.0019397 | 0 | TBU06949.1 |
| FUN_004290-T1 | 0.00203019 | 0 | TBU06950.1 |
| FUN_004294-T1 | 0.00512581 | 0.09863615 | TBU07146.1 |
| FUN_004295-T1 | 0.0037653 | 0.0097578 | TBU07147.1 |
| FUN_004296-T1 | 0.0046226 | 0.17062623 | TBU07148.1 |
| FUN_004297-T1 | 0.0036494 | 0.33328497 | TBU01997.1 |
| FUN_004299-T1 | 0.00612639 | 0.09556413 | TBU09401.1 |
| FUN_004300-T1 | 0.00069391 | 0.03505777 | TBU09402.1 |
| FUN_004301-T1 | 0.01952394 | 1.61363334 | TBU09403.1 |
| FUN_004302-T1 | 0.01161924 | 1.00601528 | TBU09404.1 |
| FUN_004303-T1 | 0.00302668 | 0.37051057 | TBU07100.1 |
| FUN_004304-T1 | 0.00268894 | 0.73913799 | TBU07099.1 |
| FUN_004305-T1 | 0.00408674 | 0.16131248 | TBU07098.1 |
| FUN_004306-T1 | 0 | NA | TBU07097.1 |
| FUN_004307-T1 | 0.00066081 | 0.45343036 | TBU07096.1 |
| FUN_004313-T1 | 0.00119274 | 0 | TBU06808.1 |
| FUN_004314-T1 | 0.00064811 | 0.06632808 | TBU06809.1 |
| FUN_004315-T1 | 0.00213869 | 0.005902 | TBU06810.1 |
| FUN_004316-T1 | 0.00082546 | Inf | TBU06811.1 |
| FUN_004317-T1 | 0.00025006 | Inf | TBU03411.1 |
| FUN_004318-T1 | 0.00388723 | 1.0049366 | TBU03410.1 |
| FUN_004319-T1 | 0.00435959 | 8.42461946 | TBU03409.1 |
| FUN_004320-T1 | 0.0025431 | 0.7548475 | TBU04467.1 |
| FUN_004322-T1 | 0.00724392 | 0.19818459 | TBU04468.1 |
| FUN_004323-T1 | 0.00388223 | 0.09074671 | TBU07610.1 |
| FUN_004325-T1 | 0.00609057 | 0.21828652 | TBU07608.1 |
| FUN_004327-T1 | 0.00524429 | 1.34297521 | TBU07074.1 |
| FUN_004328-T1 | 0.00245167 | 0 | TBU07075.1 |
| FUN_004329-T1 | 0.00891686 | 0.80566226 | TBU07076.1 |
| FUN_004330-T1 | 0.00244277 | 0.06820684 | TBU07077.1 |
| FUN_004331-T1 | 0.00174268 | 0.08681886 | TBU07078.1 |
| FUN_004332-T1 | 0.00226458 | 0 | TBU00834.1 |
| FUN_004334-T1 | 0.00619345 | 0.20059235 | TBU08187.1 |
| FUN_004335-T1 | 0.00226604 | 0.32591196 | TBU08186.1 |
| FUN_004336-T1 | 0.00151562 | 0.11546197 | TBU08185.1 |
| FUN_004337-T1 | 0.0045002 | 0.13813193 | TBU08184.1 |
| FUN_004338-T1 | 0.00298318 | 0.12366144 | TBU08183.1 |
| FUN_004339-T1 | 0.00341268 | 0.00695648 | TBU06638.1 |
| FUN_004341-T1 | 0.00391679 | 0.3402563 | TBU06637.1 |
| FUN_004342-T1 | 0.0024065 | 0.05009753 | TBT99808.1 |
| FUN_004343-T1 | 0.00828033 | 0.45562391 | TBU04766.1 |
| FUN_004344-T1 | 0.00231241 | 0.22970493 | TBU00784.1 |
| FUN_004345-T1 | 0.00210317 | 0.39083529 | TBU04057.1 |
| FUN_004346-T1 | 0.00250451 | 0.08940304 | TBU04056.1 |
| FUN_004347-T1 | 0.00199058 | 0.12780731 | TBU03210.1 |
| FUN_004348-T1 | 0.00206365 | 0.16765242 | TBT98531.1 |
| FUN_004354-T1 | 0.00163739 | 0.16075848 | TBU04523.1 |
| FUN_004356-T1 | 0.00087977 | 0.21543946 | TBU06355.1 |
| FUN_004358-T1 | 0.0021712 | 0.34081825 | TBU00471.1 |
| FUN_004359-T1 | 0.00171739 | 0.1392847 | TBU00703.1 |
| FUN_004360-T1 | 0.00109349 | 0.04440073 | TBT98269.1 |
| FUN_004361-T1 | 0.00107527 | 0.80238501 | TBT98268.1 |
| FUN_004362-T1 | 0.00230316 | 0 | TBT97416.1 |
| FUN_004363-T1 | 0.00119164 | 0 | TBU01775.1 |
| FUN_004364-T1 | 0.00244321 | 0.07599554 | TBU01774.1 |
| FUN_004366-T1 | 0 | NA | TBU08425.1 |
| FUN_004367-T1 | 0.00335359 | 3.89176537 | TBU05845.1 |
| FUN_004368-T1 | 0.0015951 | 0.09232232 | TBU05844.1 |
| FUN_004370-T1 | 0.00161069 | 0.14453087 | TBU05843.1 |
| FUN_004371-T1 | 0.00112202 | Inf | TBU05842.1 |
| FUN_004372-T1 | 0.00114018 | 0.30452642 | TBU02938.1 |
| FUN_004374-T1 | 0.00582437 | 0.54196484 | TBU09225.1 |
| FUN_004375-T1 | 0.0016867 | 0.08719348 | TBU09224.1 |
| FUN_004376-T1 | 0.00410881 | 0.69530291 | TBU09223.1 |
| FUN_004377-T1 | 0.00208793 | 0.09456605 | TBU09222.1 |
| FUN_004378-T1 | 0.00266941 | 0.09632551 | TBU09221.1 |
| FUN_004379-T1 | 0.00157504 | 0 | TBT97779.1 |
| FUN_004381-T1 | 0.0008099 | 2.34425522 | TBU08677.1 |
| FUN_004382-T1 | 0.00333895 | 0.1024669 | TBU08676.1 |
| FUN_004383-T1 | 0.00323831 | 0.21641201 | TBU08675.1 |
| FUN_004385-T1 | 0.00447592 | 0.36238825 | TBU08674.1 |
| FUN_004402-T1 | 0.0192219 | 0.72307925 | TBU01251.1 |
| FUN_004406-T1 | 0.00198651 | 0.09006118 | TBT98667.1 |
| FUN_004407-T1 | 0.00467919 | 0.25568745 | TBU01803.1 |
| FUN_004411-T1 | 0.00393524 | 0.27317451 | TBU02698.1 |
| FUN_004412-T1 | 0.00230694 | 0.03087356 | TBU01268.1 |
| FUN_004418-T1 | 0.00905152 | 0.67044122 | TBT98002.1 |
| FUN_004422-T1 | 0.00085345 | 0.19503715 | TBU07995.1 |
| FUN_004423-T1 | 0.00451273 | 0.11073368 | TBU07993.1 |
| FUN_004424-T1 | 0.0038997 | 0.2671142 | TBU07991.1 |
| FUN_004427-T1 | 0.00369588 | 0.32865017 | TBU06076.1 |
| FUN_004428-T1 | 0.0108027 | 0.24012157 | TBU06077.1 |
| FUN_004429-T1 | 0.00450794 | 0.13613342 | TBU06078.1 |
| FUN_004430-T1 | 0.00287808 | 0.11098703 | TBU06079.1 |
| FUN_004431-T1 | 0.00462162 | 0.45109405 | TBU07500.1 |
| FUN_004432-T1 | 0.00447214 | 0.00727926 | TBU07499.1 |
| FUN_004433-T1 | 0.00054944 | 0.12151517 | TBU07498.1 |
| FUN_004434-T1 | 0.00264718 | 0.05672851 | TBU07497.1 |
| FUN_004435-T1 | 0.00237056 | 0.40323901 | TBU07496.1 |
| FUN_004437-T1 | 0.00587157 | 0.12330356 | TBU04212.1 |
| FUN_004438-T1 | 0.00560909 | 0.60131603 | TBU04213.1 |
| FUN_004439-T1 | 0.00308506 | 0.11330953 | TBU05497.1 |
| FUN_004440-T1 | 0.00118056 | 0.10779088 | TBU05496.1 |
| FUN_004446-T1 | 0.00160591 | 0.28326054 | TBU08193.1 |
| FUN_004447-T1 | 0.00100358 | 0 | TBU08192.1 |
| FUN_004448-T1 | 0.00377217 | 0.13712683 | TBU01553.1 |
| FUN_004451-T1 | 0.00112393 | 0.21023487 | TBU01458.1 |
| FUN_004452-T1 | 0.00135921 | Inf | TBU01457.1 |
| FUN_004454-T1 | 0.00059737 | 0 | TBU07958.1 |
| FUN_004460-T1 | 0.00067139 | 0 | TBU07957.1 |
| FUN_004461-T1 | 0.0024793 | 0.17824787 | TBU07956.1 |
| FUN_004462-T1 | 0.00137323 | 0.17230193 | TBT99335.1 |
| FUN_004463-T1 | 0.0013408 | 0.07146225 | TBU02080.1 |
| FUN_004465-T1 | 0.00373383 | 0.67885991 | TBU08969.1 |
| FUN_004466-T1 | 0.00262164 | 0.31107015 | TBU08970.1 |
| FUN_004467-T1 | 0.01047218 | 0.26393539 | TBU08971.1 |
| FUN_004469-T1 | 0.00108215 | 0.11563953 | TBU08972.1 |
| FUN_004471-T1 | 0.00164797 | 0.11631485 | TBU06461.1 |
| FUN_004472-T1 | 0.00038757 | 0 | TBU06460.1 |
| FUN_004473-T1 | 0.0011963 | 0.24355599 | TBU06459.1 |
| FUN_004474-T1 | 0.00081555 | 0.29465717 | TBU06458.1 |
| FUN_004475-T1 | 0.00031167 | Inf | TBU06457.1 |
| FUN_004476-T1 | 0.00670619 | 0.20749474 | TBU02460.1 |
| FUN_004477-T1 | 0.0026589 | 0.07121278 | TBU02461.1 |
| FUN_004478-T1 | 0.00365912 | 0.94914563 | TBU01520.1 |
| FUN_004480-T1 | 0.00167877 | 0.06512527 | TBU05151.1 |
| FUN_004481-T1 | 0.00286167 | 0.57020169 | TBU05150.1 |
| FUN_004482-T1 | 0.00137477 | 0 | TBU05149.1 |
| FUN_004486-T1 | 0.00081405 | 0.06840413 | TBU05962.1 |
| FUN_004487-T1 | 0.00305874 | 0.1901506 | TBU06127.1 |
| FUN_004488-T1 | 0.00217963 | 0.03927536 | TBU06128.1 |
| FUN_004489-T1 | 0.00051781 | 0.11291956 | TBU06130.1 |
| FUN_004490-T1 | 0.00299412 | 0.14120472 | TBT98020.1 |
| FUN_004491-T1 | 0.00946604 | 0.11431826 | TBU04035.1 |
| FUN_004492-T1 | 0.00273162 | 0.01493931 | TBU04034.1 |
| FUN_004494-T1 | 0.00319588 | 0.36837081 | TBU05850.1 |
| FUN_004496-T1 | 0.00296121 | 1.77581666 | TBU03399.1 |
| FUN_004498-T1 | 0.00188916 | 0.22939522 | TBU08946.1 |
| FUN_004500-T1 | 0.00261751 | 0.54214156 | TBU08945.1 |
| FUN_004501-T1 | 0.00428738 | 0.18518106 | TBU08944.1 |
| FUN_004502-T1 | 0.00547644 | 0.30176182 | TBU08943.1 |
| FUN_004503-T1 | 0.00279193 | 0.01018009 | TBU08942.1 |
| FUN_004504-T1 | 0.00321505 | 0.56054078 | TBU08941.1 |
| FUN_004505-T1 | 0.0054409 | 0.41182438 | TBU05548.1 |
| FUN_004507-T1 | 0.0071534 | 0.30001615 | TBU05371.1 |
| FUN_004508-T1 | 0.00391109 | 0.33882592 | TBU05372.1 |
| FUN_004509-T1 | 0.00420206 | 0.44148322 | TBU05373.1 |
| FUN_004514-T1 | 0 | NA | TBT98810.1 |
| FUN_004515-T1 | 0.00350294 | 0.32903901 | TBU01363.1 |
| FUN_004517-T1 | 0.00194432 | 0.10727103 | TBU01362.1 |
| FUN_004518-T1 | 0.00283968 | 0 | TBU03074.1 |
| FUN_004519-T1 | 0.00687359 | 0.41228997 | TBU03075.1 |
| FUN_004523-T1 | 0.00175936 | 0.13870095 | TBU06013.1 |
| FUN_004526-T1 | 0.00584822 | 0.30293906 | TBU06659.1 |
| FUN_004527-T1 | 0.00718894 | 0.27653549 | TBU06658.1 |
| FUN_004528-T1 | 0.00700231 | 0.57438553 | TBU06657.1 |
| FUN_004529-T1 | 0.00226678 | 0 | TBU06656.1 |
| FUN_004530-T1 | 0.01488614 | 0.53133298 | TBT99624.1 |
| FUN_004534-T1 | 0.00186343 | 0.07293879 | TBT98317.1 |
| FUN_004535-T1 | 0.00194773 | 0.16510828 | TBU01317.1 |
| FUN_004538-T1 | 0.01109692 | 0.33830741 | TBU07140.1 |
| FUN_004539-T1 | 0.00058179 | 0 | TBU07141.1 |
| FUN_004542-T1 | 0.01037986 | 0.72959904 | TBU02412.1 |
| FUN_004543-T1 | 0.0090197 | 0.92521454 | TBT99712.1 |
| FUN_004544-T1 | 0.00975498 | 0.39795158 | TBU04827.1 |
| FUN_004545-T1 | 0.00721237 | 0.31188515 | TBU04826.1 |
| FUN_004546-T1 | 0.01037861 | 0.7866495 | TBU08208.1 |
| FUN_004547-T1 | 0.0030513 | 0.2641352 | TBU08209.1 |
| FUN_004549-T1 | 0.01936415 | 0.55356577 | TBU08210.1 |
| FUN_004550-T1 | 0.02272767 | 0.24598524 | TBU08211.1 |
| FUN_004551-T1 | 0.00324888 | 0.05546595 | TBU08212.1 |
| FUN_004552-T1 | 0.00263303 | 0 | TBU05190.1 |
| FUN_004553-T1 | 0.00392535 | 5.38018504 | TBU05189.1 |
| FUN_004554-T1 | 0.00197862 | 0.51947455 | TBU05188.1 |
| FUN_004555-T1 | 0.00023503 | 0.23370787 | TBU09368.1 |
| FUN_004556-T1 | 0.00191383 | 0.58912435 | TBU09370.1 |
| FUN_004557-T1 | 0.00103789 | 0.1769751 | TBU09371.1 |
| FUN_004558-T1 | 0.00019031 | 0.24673014 | TBU09372.1 |
| FUN_004559-T1 | 0.00013741 | 0.22265625 | TBU09373.1 |
| FUN_004560-T1 | 0.00061194 | 0.33167158 | TBU09374.1 |
| FUN_004561-T1 | 0.00062986 | 0.60098032 | TBU09375.1 |
| FUN_004562-T1 | 0.00026186 | 0.05306041 | TBU09376.1 |
| FUN_004563-T1 | 0.00013525 | 0 | TBU09377.1 |
| FUN_004565-T1 | 0.00042726 | 0.11998175 | TBU06271.1 |
| FUN_004566-T1 | 0.00046921 | 0.48626841 | TBU06270.1 |
| FUN_004567-T1 | 0.00011201 | Inf | TBU06269.1 |
| FUN_004568-T1 | 0.00073817 | 0.44817927 | TBU06268.1 |
| FUN_004570-T1 | 0.00012431 | 0 | TBU07171.1 |
| FUN_004571-T1 | 0.00024518 | 0.07913961 | TBU07172.1 |
| FUN_004572-T1 | 0.00015039 | 0 | TBU07173.1 |
| FUN_004573-T1 | 0.00089312 | 0.2629336 | TBU07174.1 |
| FUN_004574-T1 | 0.00059371 | 0.20938211 | TBU07175.1 |
| FUN_004575-T1 | 0.00036959 | 0.72414252 | TBU04891.1 |
| FUN_004576-T1 | 0.00019116 | Inf | TBU04892.1 |
| FUN_004577-T1 | 0.00046728 | Inf | TBU04893.1 |
| FUN_004580-T1 | 0.000361 | 0.0877193 | TBU00825.1 |
| FUN_004582-T1 | 0.0001401 | Inf | TBU04786.1 |
| FUN_004583-T1 | 0.00037281 | 2.05654131 | TBU04785.1 |
| FUN_004584-T1 | 9.35E-05 | Inf | TBU04784.1 |
| FUN_004587-T1 | 0.00068271 | 0.22376941 | TBU08204.1 |
| FUN_004589-T1 | 0.00082472 | 3.30512962 | TBU08203.1 |
| FUN_004590-T1 | 0.00094877 | 0.37608212 | TBU08202.1 |
| FUN_004591-T1 | 0.00012116 | 0 | TBU08201.1 |
| FUN_004592-T1 | 0 | NA | TBU08200.1 |
| FUN_004593-T1 | 0.00099994 | 0 | TBU02441.1 |
| FUN_004607-T1 | 0.00152389 | 1.36942368 | TBU09644.1 |
| FUN_004610-T1 | 0.01321462 | 0.65637942 | TBU06366.1 |
| FUN_004611-T1 | 0.00294478 | 0.34345497 | TBU06367.1 |
| FUN_004612-T1 | 0.00644054 | 0.35359278 | TBU08462.1 |
| FUN_004613-T1 | 0.00214915 | 0.21585571 | TBU08461.1 |
| FUN_004615-T1 | 0.00691627 | 0.35288928 | TBU08460.1 |
| FUN_004616-T1 | 0.00335287 | 0.22893479 | TBU08459.1 |
| FUN_004617-T1 | 0.00445874 | 0.18431959 | TBU08458.1 |
| FUN_004618-T1 | 0.00401179 | 0.10123437 | TBU08457.1 |
| FUN_004620-T1 | 0.00865916 | 0.27008067 | TBU01383.1 |
| FUN_004621-T1 | 0.00393726 | 0.11096436 | TBT98451.1 |
| FUN_004622-T1 | 0.00551446 | 0.64965267 | TBT99608.1 |
| FUN_004631-T1 | 0.00402844 | 0.1768215 | TBU07219.1 |
| FUN_004637-T1 | 0.00113308 | Inf | TBU08858.1 |
| FUN_004638-T1 | 0.00265371 | 0.23405596 | TBU08859.1 |
| FUN_004639-T1 | 0.00213726 | 0.57871976 | TBU08860.1 |
| FUN_004644-T1 | 0.00239114 | 0.36964239 | TBU08034.1 |
| FUN_004645-T1 | 0.00267178 | 0.35190672 | TBU07430.1 |
| FUN_004647-T1 | 0 | NA | TBU07431.1 |
| FUN_004648-T1 | 0.0081008 | 0.33058056 | TBU07432.1 |
| FUN_004649-T1 | 0.00110269 | 0.03346445 | TBU09494.1 |
| FUN_004651-T1 | 0.00973521 | 0.24339487 | TBU09495.1 |
| FUN_004653-T1 | 0.00558782 | 0.21559505 | TBU09496.1 |
| FUN_004657-T1 | 0.00502774 | 0.79316722 | TBU09498.1 |
| FUN_004658-T1 | 0.00308024 | 0.74243217 | TBT99452.1 |
| FUN_004659-T1 | 0.01097051 | 0.23772513 | TBU08047.1 |
| FUN_004665-T1 | 0.00670069 | 0.20077707 | TBU08637.1 |
| FUN_004666-T1 | 0.00686639 | 0.2930916 | TBU08638.1 |
| FUN_004671-T1 | 0.00456487 | 0.52877978 | TBU09658.1 |
| FUN_004674-T1 | 0.00337578 | 0.01197888 | TBU06088.1 |
| FUN_004675-T1 | 0.00353606 | 0.04224085 | TBU06087.1 |
| FUN_004679-T1 | 0.00195653 | 0.02704768 | TBU06086.1 |
| FUN_004680-T1 | 0.00298173 | 0.41242796 | TBU00809.1 |
| FUN_004681-T1 | 0.02337642 | 0.69402797 | TBU05759.1 |
| FUN_004683-T1 | 0.00273983 | 3.92490766 | TBU02425.1 |
| FUN_004684-T1 | 0.00430963 | 0.09559865 | TBU02426.1 |
| FUN_004685-T1 | 0.00954594 | 0.74300537 | TBU02427.1 |
| FUN_004687-T1 | 0.00312067 | 0.07238999 | TBU01066.1 |
| FUN_004688-T1 | 0.00509623 | 0.23432304 | TBU01067.1 |
| FUN_004691-T1 | 0.00971631 | 0.32072735 | TBU07304.1 |
| FUN_004694-T1 | 0.00518706 | 5.47024259 | TBU02475.1 |
| FUN_004695-T1 | 0.00270305 | 0.24813601 | TBU02474.1 |
| FUN_004698-T1 | 0.00488217 | 0.33273085 | TBU06563.1 |
| FUN_004699-T1 | 0.0034603 | 0.13543926 | TBU07560.1 |
| FUN_004700-T1 | 0.00747995 | 0.21103502 | TBU07559.1 |
| FUN_004702-T1 | 0.00278744 | 0.24982558 | TBU07558.1 |
| FUN_004703-T1 | 0.00454545 | 0.04331493 | TBU07557.1 |
| FUN_004704-T1 | 0.00200717 | Inf | TBU07556.1 |
| FUN_004706-T1 | 0.00157298 | 0.0871556 | TBU04030.1 |
| FUN_004707-T1 | 0.00405252 | 0.04478964 | TBU04029.1 |
| FUN_004708-T1 | 0.01023656 | 1.34110317 | TBU04028.1 |
| FUN_004711-T1 | 0.00191159 | 0.1016426 | TBU02645.1 |
| FUN_004713-T1 | 0.0034351 | 0.03851912 | TBU02855.1 |
| FUN_004714-T1 | 0.00550733 | 0.2154059 | TBU03115.1 |
| FUN_004726-T1 | 0.00356669 | 0.12486161 | TBU05634.1 |
| FUN_004728-T1 | 0.00013193 | 0 | TBT99339.1 |
| FUN_004729-T1 | 0.008348 | 0.32003749 | TBU01905.1 |
| FUN_004730-T1 | 0.00199809 | 0.47470238 | TBU01904.1 |
| FUN_004735-T1 | 0.00284814 | 0.79883413 | TBU05997.1 |
| FUN_004739-T1 | 0.01389578 | 0.45497468 | TBU08025.1 |
| FUN_004742-T1 | 0.01363077 | 0.57154061 | TBU03307.1 |
| FUN_004748-T1 | 0.00610986 | 0.04938761 | TBU03930.1 |
| FUN_004749-T1 | 0.00633492 | 0.1756224 | TBT99673.1 |
| FUN_004750-T1 | 0.00240732 | 0.11783561 | TBT99672.1 |
| FUN_004752-T1 | 0.00957189 | 0.71167707 | TBU08481.1 |
| FUN_004754-T1 | 0.00187745 | 5.05562418 | TBU08482.1 |
| FUN_004755-T1 | 0.01300286 | 0.94069193 | TBU08483.1 |
| FUN_004756-T1 | 0.00181371 | Inf | TBU07531.1 |
| FUN_004761-T1 | 0.00138998 | 0.4404721 | TBU07596.1 |
| FUN_004766-T1 | 0.00184363 | 0.2522317 | TBU07254.1 |
| FUN_004767-T1 | 0.00091859 | Inf | TBU07255.1 |
| FUN_004771-T1 | 0.01599613 | 0.85675937 | TBU08572.1 |
| FUN_004772-T1 | 0.00451613 | 0.38856305 | TBU08571.1 |
| FUN_004774-T1 | 0.00266875 | 0.27528895 | TBU07266.1 |
| FUN_004775-T1 | 0.00158322 | 0.62809807 | TBU07267.1 |
| FUN_004778-T1 | 0.00103603 | Inf | TBU08551.1 |
| FUN_004780-T1 | 0.00686042 | 0.39485835 | TBU08550.1 |
| FUN_004784-T1 | 0.0016535 | 0.45023182 | TBU07050.1 |
| FUN_004785-T1 | 0.00319047 | 0.24037956 | TBU02969.1 |
| FUN_004794-T1 | 0.0085554 | 0.29314295 | TBU05783.1 |
| FUN_004795-T1 | 0.00047967 | 0.04503649 | TBU04404.1 |
| FUN_004797-T1 | 0.00136445 | 0.21522878 | TBU03631.1 |
| FUN_004808-T1 | 0.00056724 | 0.20303697 | TBU09013.1 |
| FUN_004809-T1 | 0.0014672 | 0.99147195 | TBU09014.1 |
| FUN_004810-T1 | 0.00390582 | 0.26474293 | TBU09015.1 |
| FUN_004811-T1 | 0.00214517 | 0.04297803 | TBT98917.1 |
| FUN_004813-T1 | 0.00198721 | 0.35647268 | TBU07550.1 |
| FUN_004814-T1 | 0.00092696 | 0.13693868 | TBU07549.1 |
| FUN_004816-T1 | 0.00711744 | 1.39227753 | TBU07547.1 |
| FUN_004818-T1 | 0.00198153 | 0.71908652 | TBU06468.1 |
| FUN_004819-T1 | 0.00056926 | 0.12044607 | TBU06466.1 |
| FUN_004821-T1 | 0.00046751 | 0.03799435 | TBU04996.1 |
| FUN_004824-T1 | 0.000729 | 0.14395797 | TBT98959.1 |
| FUN_004825-T1 | 0.00671583 | 0.37627912 | TBT98056.1 |
| FUN_004826-T1 | 0.00673081 | 0.26871529 | TBT96946.1 |
| FUN_004828-T1 | 0.00834646 | 0.79535322 | TBU07625.1 |
| FUN_004830-T1 | 0.03622829 | 0.88247863 | TBU07623.1 |
| FUN_004831-T1 | 0.00461725 | 0.10837566 | TBU07622.1 |
| FUN_004833-T1 | 0.00236374 | 0.03174055 | TBT99908.1 |
| FUN_004835-T1 | 0.01957541 | 0.49444152 | TBU01115.1 |
| FUN_004836-T1 | 0.0021227 | 0.16021298 | TBU07703.1 |
| FUN_004838-T1 | 0.00233326 | 0.08687134 | TBU07704.1 |
| FUN_004839-T1 | 0.00307796 | 0.1421468 | TBU07705.1 |
| FUN_004841-T1 | 0.00107527 | Inf | TBU05642.1 |
| FUN_004842-T1 | 0.00147815 | 0.01698956 | TBU05641.1 |
| FUN_004843-T1 | 0.00304287 | 0.0358958 | TBU05640.1 |
| FUN_004844-T1 | 0.01099164 | 0.73150323 | TBU06975.1 |
| FUN_004845-T1 | 0.00333141 | 0.26179657 | TBU06974.1 |
| FUN_004846-T1 | 0.00076019 | 0 | TBU06973.1 |
| FUN_004851-T1 | 0.00151119 | 1.78531534 | TBU01238.1 |
| FUN_004859-T1 | 0.00936854 | 0.85671675 | TBU02037.1 |
| FUN_004867-T1 | 0.00393954 | 0.36051477 | TBU04237.1 |
| FUN_004868-T1 | 0.00065168 | 0 | TBU06348.1 |
| FUN_004870-T1 | 0.00170761 | 0.73813595 | TBU06350.1 |
| FUN_004872-T1 | 0.00412592 | 0.26404676 | TBU04486.1 |
| FUN_004873-T1 | 0.00792747 | 0.58008181 | TBU04485.1 |
| FUN_004877-T1 | 0.00382677 | 0.04200642 | TBU08042.1 |
| FUN_004878-T1 | 0.0066932 | 0.36900118 | TBT98099.1 |
| FUN_004880-T1 | 0.00466184 | 0.28660028 | TBU01894.1 |
| FUN_004889-T1 | 0.00289606 | 0.28286787 | TBU06389.1 |
| FUN_004894-T1 | 0.00374515 | 0.46002941 | TBU09579.1 |
| FUN_004895-T1 | 0 | NA | TBU09580.1 |
| FUN_004896-T1 | 0.00247268 | 0.14200971 | TBU09581.1 |
| FUN_004897-T1 | 0.0002545 | 0.22658099 | TBU09582.1 |
| FUN_004898-T1 | 0.00651549 | 0.08547622 | TBU09583.1 |
| FUN_004904-T1 | 0.00053763 | 0.73400251 | TBU09415.1 |
| FUN_004905-T1 | 0.0033537 | 0.14085957 | TBU09416.1 |
| FUN_004907-T1 | 0.00461579 | 0.85672148 | TBT98112.1 |
| FUN_004910-T1 | 0.00146307 | 0.93655633 | TBU04845.1 |
| FUN_004911-T1 | 0.00614439 | 0.13093995 | TBU04846.1 |
| FUN_004912-T1 | 0.00539239 | 0.5797781 | TBU01975.1 |
| FUN_004913-T1 | 0.00268436 | 0.31085934 | TBU01976.1 |
| FUN_004914-T1 | 0.00946959 | 0.50985147 | TBU01977.1 |
| FUN_004915-T1 | 0.00630663 | 0.25677386 | TBU05556.1 |
| FUN_004916-T1 | 0.00121893 | 0 | TBU05557.1 |
| FUN_004947-T1 | 0.01889803 | 0.44381116 | TBT96833.1 |
| FUN_004948-T1 | 0.00036369 | Inf | TBT98488.1 |
| FUN_004950-T1 | 0.00202551 | 0.04993214 | TBU07515.1 |
| FUN_004952-T1 | 0.00309677 | 0.14390871 | TBU07516.1 |
